# Bayesian cell-type deconvolution and gene expression inference reveals tumor-microenvironment interactions

**DOI:** 10.1101/2020.01.07.897900

**Authors:** Tinyi Chu, Zhong Wang, Dana Pe’er, Charles G. Danko

**Author notes:** **<u>Address correspondence to</u>:** Charles G. Danko, Ph.D., Tinyi Chu, Ph.D.

## Abstract

Understanding the interactions between cells in their environment is a major challenge in genomics. Here we developed BayesPrism, a Bayesian method to jointly predict cellular composition and gene expression in each cell type, including heterogeneous malignant cells, from bulk RNA-seq using scRNA-seq as prior information. We conducted an integrative analysis of 1,412 bulk RNA-seq samples in primary glioblastoma, head and neck squamous cell carcinoma, and melanoma using single-cell datasets of 85 patients. We identified cell types correlated with clinical outcomes and explored spatial heterogeneity in malignant cell states and non-malignant cell type composition. We refined subtypes using gene expression in malignant cells, after excluding confounding non-malignant cell types. Finally, we identified genes whose expression in malignant cells correlated with infiltration of macrophages, T cells, fibroblasts, and endothelial cells across multiple tumor types. Our work introduces a new lens that uses scRNA-seq to accurately infer cellular composition and expression in large cohorts of bulk data.

## Introduction

Cell-cell interactions are highly complex and can strongly impact cell behavior in biological contexts, often with medical ramifications. A quintessential example is between malignant cells and diverse non-malignant cell types within the tumor microenvironment (TME)^1–3^. Numerous studies over the past two decades have revealed interactions between cells in the TME that promote diverse functions, including angiogenesis^4,5^, metastasis^6^, and immunosuppression^7,8^. Non-malignant cells can differ markedly between patients and tumor types^9–15^, and certain non-malignant cell populations are used as clinical biomarkers^16–19^ and therapeutic targets^20–25^. These studies motivate the direct measurement of cell types within tissues.

Two layers of information are critical for understanding tumor composition: (1) the proportion of each cell type, and (2) the levels of gene expression in each cell type. The rise of single-cell RNA sequencing (scRNA-seq)^26–31^ technologies has recently enabled direct, genome-wide measurement of the transcriptome in individual cells within the TME and characterization of their heterogeneity. However, the cost of scRNA-seq and requirements for high-quality tissue limits the number of patient samples that can be assayed^32,33^. Moreover, scRNA-seq is susceptible to technical biases in cell capture^32,33^, which confound the recovery of cell type composition.

As an alternative, cell type abundance can be inferred from bulk RNA-seq data using regression on a reference expression matrix constructed from a set of arbitrarily defined marker genes^17,18,34,35^. Pioneering methods for cell type deconvolution have demonstrated that it is possible to infer the abundance of multiple cell types in the TME. However, existing deconvolution methods make restrictive assumptions about the difference in distribution between the reference and bulk sample. These assumptions are often violated by both technical (e.g., different platforms are used for reference and bulk sequencing) and biological (e.g., heterogeneity in gene expression within constituent cell types^36–38^) differences between bulk and reference data. Critically, cell type deconvolution methods have not fully supported predicting gene expression in a heterogeneous population of tumor cells. Thus, existing methods fail to address key questions: How do malignant cells affect the composition of non-malignant cells in the tumor niche? Which genes are correlated with these interactions? To answer these questions, we need a model that can accurately represent cell-type fraction and cell-type-specific expression profiles in each bulk sample, and can accommodate differences between the single-cell reference and bulk.

Here, we present BayesPrism (Bayesian cell Proportion Reconstruction Inferred using Statistical Marginalization), a Bayesian model that jointly infers the posterior distribution of cell type fractions and gene expression from bulk RNA-seq data using an scRNA-seq reference as prior information. By explicitly modeling and marginalizing out the differences in gene expression between the single-cell reference and the bulk data, BayesPrism substantially outperforms leading methods in the inference of cell type fractions in both tumor and non-tumor settings. We demonstrate the utility of our approach on a large dataset of 1,412 bulk RNA-seq and 85 scRNA-seq samples in primary glioblastoma, head and neck squamous cell carcinoma, and melanoma. Our work introduces a powerful new tool for integrative analysis of bulk and scRNA-seq data.

## Results

### Bayesian inference of cell type composition and gene expression

BayesPrism uses an scRNA-seq reference to infer two statistics from each bulk RNA-seq sample: (i) the proportion of reads derived from each cell type, which we assume is proportional to the fraction of that cell type, and (ii) the expression level of genes in each cell type (**Fig. 1a**,**b, Supplementary Fig. 1, Supplementary Note 1**). The most challenging aspect of cellular deconvolution is accounting for various sources of uncertainty, including technical and biological batch variation, in gene expression between the bulk and scRNA-seq reference data. To account for these uncertainties, BayesPrism adopts a Bayesian strategy which models a prior distribution using scRNA-seq, and infers a joint posterior distribution of cell type proportion and gene expression in each cell type and bulk sample conditional on each observed bulk. As a result,the uncertainty in each estimate can be marginalized out from the joint posterior. As scRNA-seq can define multiple gene expression subtypes of the same cell type (hereafter referred to as “cell states”), BayesPrism allows users to input a refined scRNA-seq reference by labeling both cell type and cell states. Internally, BayesPrism treats cell types and states in the same manner, and returns the posterior sum over different cell states defined in the scRNA-seq dataset to represent the fraction and expression of each cell type. This strategy is useful in modeling heterogeneous cell types, including both malignant and non-malignant cell types in the TME.

**Fig. 1.**
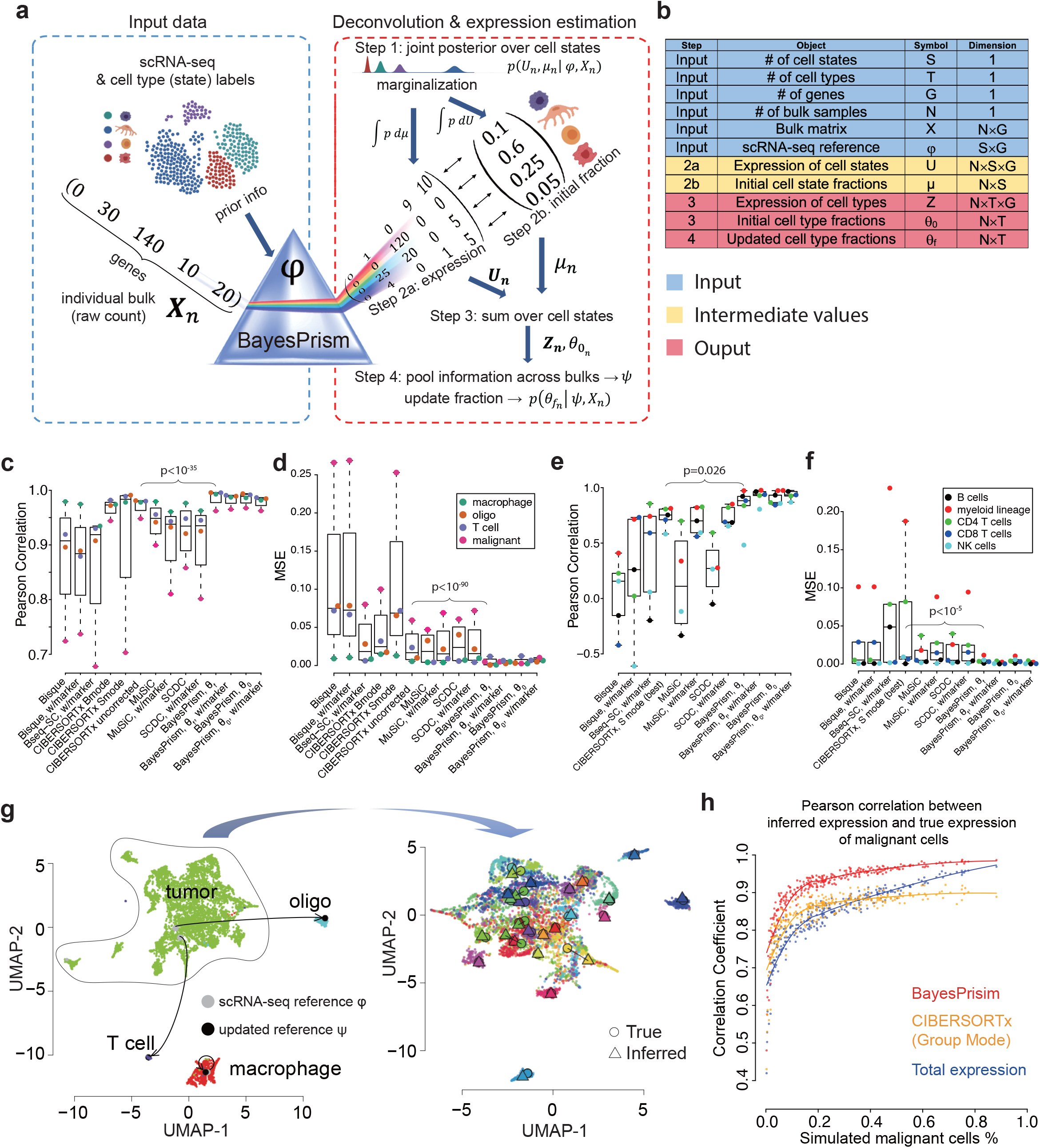
BayesPrism algorithm flow and performance validation. **a**) Algorithmist flow of the deconvolution module of BayesPrism. **b**) Variables and their dimensions shown in (**a**). **c**-**f**) Boxplots show the cell type-level Pearson’s correlation coefficient and MSE for deconvolution of pseudo bulks of GBM28 using refGBM8 (**c** and **d**), and bulk RNA-seq human whole blood samples with the ground truth measured by flow-cytometry (**e** and **f**). Boxes mark the 25th percentile (bottom of box), median (central bar), and 75th percentile (top of box). Whiskers represent extreme values within 1.5 fold of the inter quartile range. One-sided p values were shown for cell type fractions inferred by BayesPrism (updated θ using the marker free mode) and those by the second best methods ranked by the median value. T test was used for MSE and z test was performed on Fisher’s Z-transformed cell type-level correlation coefficients (see Methods). **g**) UMAP visualization shows the expression of individual cells in GBM28. The expression profiles of non-malignant cells before (gray) and after (black) information-pooling were projected onto the UMAP manifold of the scRNA-seq (left). Malignant cells in patients with greater than 10 malignant cells (N=27) were visualized on the zoomed-in UMAP (right), and are colored by patient. The inferred expression profile, shown as ○, and the averaged expression profile from scRNA-seq for each patient, shown as △, are projected onto the UMAP manifold. h) Scatter plot shows Pearson’s correlation between the average expression of malignant cells in pseudo-bulk and that estimated by BayesPrism (red) and CIBERSORTx group mode (orange) or the undeconvolved simulated bulk (blue), as a function of the fraction of malignant cells in a subsampled set (N=270).

BayesPrism is implemented in an efficient algorithm (See **Methods** and **Supplementary Note 2**) which consists of four major steps:

1. BayesPrism first infers a joint posterior distribution of the cell state proportion and gene expression, *μ*_*n*_ and *U*_*n*_, conditional on the observed single cell reference *φ* (obtained by summing over the count matrix of each cell state followed by normalizing by the total count), and read counts of bulk expression *X*_*n*_ of the *n*_*th*_ bulk sample, i.e. *p(μ*_*n*_, *U*_*n*_ | *φ, X*_*n*_), using Gibbs sampling.
2. For each bulk sample *n*, BayesPrism estimates (Step 2a) the gene expression matrix of each cell state, *U*_*n*_, and (Step 2b) the proportion of each cell state, *μ*_*n*_, by marginalizing the joint posterior and reporting the posterior mean. Explicitly modeling the cell type-specific expression value in each bulk makes BayesPrism robust to technical batch effects and biological variation between the scRNA-seq reference and the bulk (see **Supplementary Note 3** for a discussion of the mathematical intuition).
3. For each bulk sample *n*, BayesPrism estimates the gene expression matrix of each cell type, *Z*_*n*_, and the proportion of each cell type,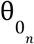, by summing up the posteriors over cell states, estimated in Step 2, within each cell type.
4. Optionally, BayesPrism updates the reference matrix *φ* by pooling information across bulk samples from *Z* to improve estimates of cell type fractions. The updated reference matrix, *ψ*, represents the multinomial distribution parameters describing the distribution of *Z*. Two strategies are used to infer *ψ* depending on the setting. First, BayesPrism can infer a maximum likelihood estimate (MLE) of cell type-specific gene expression that is unique to each bulk sample. This feature allows users to estimate gene expression in each bulk sample when there is substantial heterogeneity, as is generally the case with malignant cells from different patients^26–31^. Second, BayesPrism uses all bulk samples to summarize a maximum a posterior (MAP) estimate of cell type-specific expression. For most cell types, gene expression is reasonably similar across bulk samples^27,29,30^, and in these cases it is appropriate to share information between samples by estimating *ψ* using all bulk data. BayesPrism then uses the updated prior distribution parameterized by *ψ* to re-sample the marginal posterior of cell type composition for each bulk sample, *θ*, i.e. 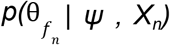. Sharing information across bulk samples results in a shrinkage property in the estimates, and provides higher accuracy for problems with batch effects.

### BayesPrism improves cell type deconvolution accuracy by accommodating uncertainty in the reference

To benchmark the robustness of cell type deconvolution against linear noise, the most common type of mathematical model for technical batch effects, we first constructed pseudo-bulk RNA-seq data that differed from the reference scRNA-seq dataset by a log-normally distributed multiplicative noise term (See Methods; **Supplementary Table 1**). We compared the Pearson correlation and mean squared error (MSE) between the ground truth and cell type proportions estimated using five different deconvolution methods^35,39–42^. BayesPrism was nearly invariant to the simulated noise and outperformed existing methods by up to an order of magnitude as noise increased (**Supplementary Fig. 2**, see **Supplementary Note 3** for mathematical arguments outlining why BayesPrism is invariant to linear noise). These data are consistent with our expectation that the Bayesian method outperforms existing methods by substantial margins when bulk data differs systematically from the scRNA-seq reference.

To assess whether BayesPrism improved deconvolution performance in a more realistic setting, we next generated pseudo-bulk data by combining reads from single cells of similar biological samples but collected using different scRNA-seq platforms. We benchmarked performance in three different settings: 1) technical batch effects with small amounts of biological variation using peripheral blood mononuclear cells (PBMCs) and mouse brain cortex from different healthy subjects (**Supplementary Fig. 3**), 2) biological variation with small amounts of technical noise using leave-one-out tests in datasets of three human cancer types generated by the same sequencing platforms (**Supplementary Fig. 4 and 5**), and 3) a mixture of technical and biological variation using glioblastoma (GBM) datasets generated from different cohorts using different sequencing platforms (**Fig. 1c-d; Supplementary Fig. 6**). When testing the effect of technical noise, we chose sequencing platforms that best recapitulate features common to bulk and scRNA-seq data modalities: full length SMART-seq2 data as a surrogate for bulk RNA-seq, and for reference scRNA-seq datasets, 3’ end-enriched tag clusters obtained using 10X (for PBMCs), sci-RNA-seq (for mouse cortex), or a microwell-based platform (for GBM). We excluded mitochondrial and ribosomal protein coding genes from all marker-free benchmarks because we found that the performance of all methods, including BayesPrism, improved when these ubiquitously expressed gene classes were excluded from the reference (**Supplementary Note 5**).

BayesPrism significantly outperformed all existing methods in all three settings in 63 of 64 tests (*p* < 0.05, see Methods). In the GBM dataset (the third setting), BayesPrism improved MSE over the next best performing method, CIBERSORTx, by ∼4-7-fold (**Supplementary Fig. 6**). BayesPrism was particularly better than CIBERSORTx in estimating the proportion of malignant cells, in which gene expression was a poor match for the reference data, consistent with our expectation that the Bayesian method will provide the highest performance advantage in the presence of substantial gene expression variation between the bulk and reference data. Separate analyses also found that BayesPrism was robust to cell types that were missing from the scRNA-seq reference (**Supplementary Fig. 7; Supplementary Note 4**), and the number of cells and scRNA-seq reference samples (**Supplementary Fig. 8**).

As a final performance benchmark, we deconvolved real bulk RNA-seq data using a ground truth obtained by orthogonal strategies. We obtained bulk RNA-seq data from 12 whole blood samples which were analyzed in parallel using flow cytometry^35^. Using PBMC scRNA-seq data as a reference, BayesPrism obtained more accurate estimates of five cell types in the bulk sample than other deconvolution methods (*p* < 5.46 × 10^−4^ on MSE; *p* < 0.03 on correlation coefficients) (**Fig. 1e-f**). Taken together, these benchmarks demonstrate that BayesPrism improves deconvolution performance in realistic settings compared with existing deconvolution methods.

### BayesPrism accurately estimates cross-patient heterogeneity in gene expression

We asked whether BayesPrism accurately recovered gene expression in heterogeneous cell types. We first focused on the recovery of gene expression in malignant cells in tumor samples. This setting is particularly challenging because heterogeneity between different patients prevents scRNA-seq reference data from accurately representing gene expression in new bulk samples^26–31^. We estimated cell types and gene expression in SMART-seq2 pseudo-bulk data from 28 GBMs, out of which 27 samples contained sufficient numbers of malignant cells for analysis (n > 10). We used a microwell-based scRNA-seq reference from 8 GBMs. As described above, we used this combination of technologies in our benchmark to mimic several of the important technical differences observed between single-cell and bulk datasets, most notably using 3’ end data to estimate expression in a full-length bulk dataset. Gene expression estimates for malignant cells in each pseudo-bulk sample (*ψ*_*mal*_) were highly similar to the known ground truth (**Fig. 1g [right]**). Next we asked how the accuracy of gene expression estimates were affected by the proportion of malignant cells. We sampled random proportions of each cell type, drawing malignant cells from a single patient. The correlation between BayesPrism gene expression estimates and the known ground truth were higher than 0.95 for tumors with >50% purity (**Fig. 1h**). Moreover, different samples simulated from the same patient produced estimates that were highly concordant with each other (**Supplementary Fig. 9**), suggesting that gene expression deconvolved by BayesPrism can accurately recover the underlying structure of gene expression in the malignant cells in bulk samples. Gene expression estimates were substantially more accurate using BayesPrism than using either CIBERSORTx or the bulk tumor with no deconvolution (**Fig. 1h; Supplementary Fig. 10**).

Next we expanded our analysis to a second cancer type, melanoma (SKCM), and examined whether BayesPrism recovered expression differences characteristic of the AXL and MITF malignant cell states. We compared BayesPrism expression estimates to AXL and MITF marker genes reported previously in the literature^27^ using a leave-one-out benchmark design. BayesPrism was able to reproduce both the AXL and MITF subtypes with similar accuracy as the original scRNA-seq data (**Supplementary Fig. 11**). Thus, BayesPrism accurately recovered sample-specific features of gene expression in heterogeneous cell types.

We asked whether BayesPrism expression estimates recovered gene expression in non-malignant cell types. Gene expression estimates aggregated across samples (*ψ*_*env*_) for macrophages, T-cells, and oligodendrocytes better matched the known ground truth than the scRNA-seq prior (**Fig. 1g [left], Supplementary Fig. 12**). To assess whether we could recover heterogeneity between patients in non-malignant cells, we used BayesPrism to estimate the gene expression of macrophages in SMART-seq2 pseudo-bulk data from 28 GBMs, using the microwell-based scRNA-seq reference from 8 GBMs in which the cell state label was provided for macrophage by clustering the scRNA-seq data. BayesPrism accurately recovered subtle variation of gene expression in macrophage cell states between simulated patients (**Supplementary Fig. 13a**). Next we sub-clustered macrophages in the scRNA-seq data used to generate pseudo-bulk and sampled random proportions of each cell type while controlling for macrophages from an individual cluster. BayesPrism accurately recovered subtle variation of gene expression in macrophage cell states between simulated pseudo-bulk samples (**Supplementary Fig. 13b-f**). Thus we conclude that BayesPrism is able to accurately recover subtle variation in gene expression between different bulk samples.

### BayesPrism reveals survival associations of immune infiltrates in three tumor types

We analyzed the proportion of cell types in 1,142 TCGA samples from three tumor types: GBM, HNSCC, and SKCM^43–45^. To maintain the highest possible accuracy, we used a scRNA-seq reference from the same tumor type in each deconvolution task^28–30^. Using these reference datasets provided estimates of 6 cell types for GBM, 10 for HNSCC, and 8 for SKCM (**Fig. 2a**). Estimates of tumor purity closely resembled those obtained using CNVs by ABSOLUTE^44–47^ and marker gene expression by ESTIMATE^47,48^, and outperformed CIBERSORTx in the majority (32/36) of comparisons (**Supplementary Fig. 14**). Likewise, estimates for the fraction of lymphocytes were correlated with those obtained by counting lymphocyte patches in H&E sections in the SKCM dataset^49^ (**Supplementary Fig. 15**). Finally, across large cohorts of tumors, non-malignant cell types had a rich correlation structure with one another that mirrored several previously described observations in the literature (**Supplementary Fig. 16, Supplementary Note 6**). Taken together, these benchmarks support the accuracy of BayesPrism in estimating cell type fractions in bulk tumor samples.

**Fig. 2.**
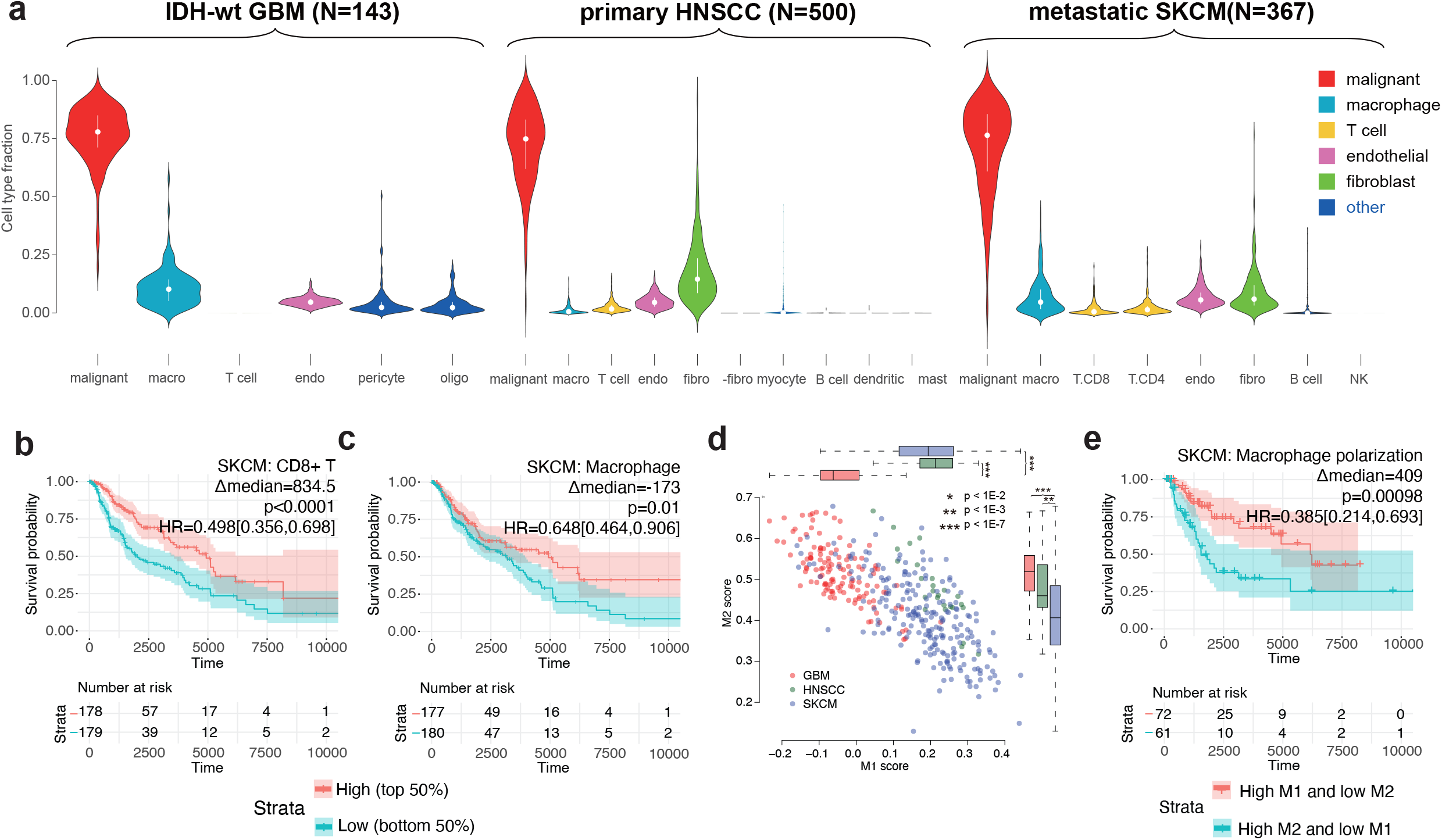
Association between prognosis and either the cell type fraction or cell state of non-malignant cells in three TCGA tumor types. **a)** Violin plots show the distribution of cell type fraction in each tumor type. Median fractions are shown by white dots and upper/lower quartiles are shown by bars. **b**-**c**) KM plots show the survival associations with **b**) T cell infiltration and **d**) macrophage infiltration in SKCM. Δmedian: median survival time in the high group - median survival time in the low group. P values were computed using the log-rank test. Hazard ratio was defined by high / low, and the 95 percentile confidence interval was shown in the square brackets. **d**) Scatter plot shows correlation between BayesPrism expression estimates in macrophages and the M1 and M2 macrophage subtype score across three tumor types. **e**) KM plots show survival associations with the M1/M2 state polarization of macrophages in SKCM.

We asked whether non-malignant cell types were correlated with patient survival. Samples contributed to TCGA have substantial differences in treatments, genetic drivers, and other confounders (**Supplementary Table 2**). To avoid confounding our analysis with known genetic or clinical covariates, we accounted for clinical features by removing subsets of samples that have a strong effect on prognosis, including IDH mutant and recurrent tumor samples in GBM, metastatic samples in HNSCC, and non-metastatic samples in SKCM. We examined the association between cell type abundance and survival using two Cox proportional hazards models by: (1) stratifying samples into *high* and *low* cell type abundance using the median value, and (2) treating cell type abundance as a continuously-valued variable (see **Methods**). In SKCM, where CD4+ and CD8+ cells were annotated separately in the reference scRNA-seq dataset, we found that CD8+ T cells had a stronger correlation with survival (HR=0.498 [0.356,0.698], **Fig. 2b** and **Supplementary Fig. 17c**), consistent with previous reports^50^. The proportion of T cells was also associated with better clinical outcomes in HNSCC, but only in the test that treated cell type abundance as a continuous variable (*p* = 0.001, Wald test, **Supplementary Fig. 17b**). BayesPrism also revealed significant positive survival associations with B cells and mast cells in HNSCC (HR=0.694 [0.509,0.948] and HR=0.668 [0.49,0.912], **Supplementary Fig. 17b**). Interestingly, the fraction of B cell and mast cells were also positively correlated (**Supplementary Fig. 16b**), suggesting that they potentially interact with each other in HNSCC and synergistically affect prognosis, consistent with reports that mast cells can promote B-cell differentiation into effector cells^51^.

The prognostic value of macrophages is more controversial than other immune cell types. Macrophage estimates by BayesPrism were positively associated with survival in SKCM (HR = 0.648 [0.464,0.906]; *p* = 0.01; log-rank test) (**Fig. 2c** and **Supplementary Fig. 17c**). In contrast, some recent studies, mostly examining other patient cohorts, have noted either no significant association or the opposite trend^50,52–55^. Intriguingly, albeit not statistically significant, we noted that high macrophage infiltration had a poor prognosis in GBM (HR = 1.18 [0.795, 1.76], **Supplementary Fig. 17a**), and no survival association in HNSCC (HR = 1.02 [0.749,1.38]) (**Supplementary Fig. 17b**). We hypothesized that differences in macrophage cell state may contribute to the differences we observed in survival associations. To test this, we used BayesPrism to estimate macrophage-specific gene expression in samples with greater than 5% macrophage content. We compared macrophage expression with marker genes characteristic of two macrophage subpopulations^34^, M1 and M2, that are believed to have different roles in the TME (See **Methods**). Macrophages from GBM had the highest M2 score and the lowest M1 score, whereas those from SKCM had the lowest M2 score and a comparably high M1 score as those from HNSCC (**Fig. 2d**). This suggests that differences in the association between macrophages and survival across the three different tumor types might be caused by macrophage polarization towards distinct transcriptional states.

We asked whether M1 / M2 macrophage polarization was associated with patient survival within each tumor type. We focused our analysis on GBM and SKCM, where >50 samples had sufficient macrophage content to estimate expression. Macrophage polarization had an extremely strong association with survival in SKCM (**Fig. 2e**). In GBM, the trend was in a consistent direction, although the log-rank test did not show statistical significance (**Supplementary Fig. 18**). Taken together, these findings highlight the importance of both macrophage content and macrophage cell state in shaping clinical outcomes across different malignancies.

### BayesPrism identifies malignant gene expression patterns correlated with cell types in the TME

Prioritizing genes that lie either upstream or downstream of interactions between malignant cells and the TME is useful for a variety of applications, such as identifying potential drug targets. We set out to develop an approach that uses the correlation between gene expression in malignant cells and the fraction of non-malignant cell types across large numbers of bulk samples to identify candidate interacting genes. Naive approaches that correlate the non-malignant cell type of interest (hereafter referred to as the “query cell type”) and the bulk tumor expression without deconvolution will result in false positive correlations from genes that are highly expressed by the query cell type. Regressing non-malignant cell type fractions out of bulk expression does not allow correlations to be estimated either, because expression estimates with each cell type will be trivially uncorrelated with cell type fraction by definition.

We therefore examined correlations between gene expression in malignant cells estimated by BayesPrism and the proportion of each query cell type. We found that BayesPrism reduced spurious correlations caused by gene expression in the query cell type (**Supplementary Fig. 19; Supplementary Note 7**). However, missing cell states from the scRNA-seq reference, a scenario often encountered in a highly heterogeneous TME, could result in transcripts that are highly expressed in a missing non-malignant cell state to be partially assigned to malignant cells by BayesPrism if the scRNA-seq of the malignant cells also show moderate expression of those genes. This partial assignment issue may cause potential false positive correlations between the estimated expression level of that gene and the fraction of the query cell type containing the missing cell states.

To further reduce potential false positive associations, we implemented two additional filters. First, we devised a likelihood ratio test to test the null hypothesis that gene expression in the query cell type alone explains the variation in the query cell fraction, effectively ameliorating false correlations introduced by cell states that may have been missed in the scRNA-seq reference (See Methods). Second, we enriched for genes intrinsic to malignant cells by selecting genes expressed at significantly higher levels in at least one malignant cell state compared to all non-malignant cell types based on the scRNA-seq reference. The combination of these two strategies ensures that we obtain a conservative set of candidate genes in which malignant cell expression correlated with non-malignant cell fraction (**Supplementary Fig. 20**).

We first asked whether we could recover known positive regulators of macrophage infiltration in IDH-wild type GBM^56,57^. Genes previously reported to have interactions all had statistically significant positive correlations with macrophage infiltration including *POSTN, ITGB1* and *LOX* (**Fig. 3a**). Likewise, several cell-surface ligands or receptors predicted to interact with macrophages by CellPhoneDB^58^, including MDK(malignant)-SORL1(macrophage), had a positive correlation with macrophage cell fraction. Thus, BayesPrism recovered multiple genes that were identified as candidates for mediating cell-cell interaction using other strategies.

**Fig. 3.**
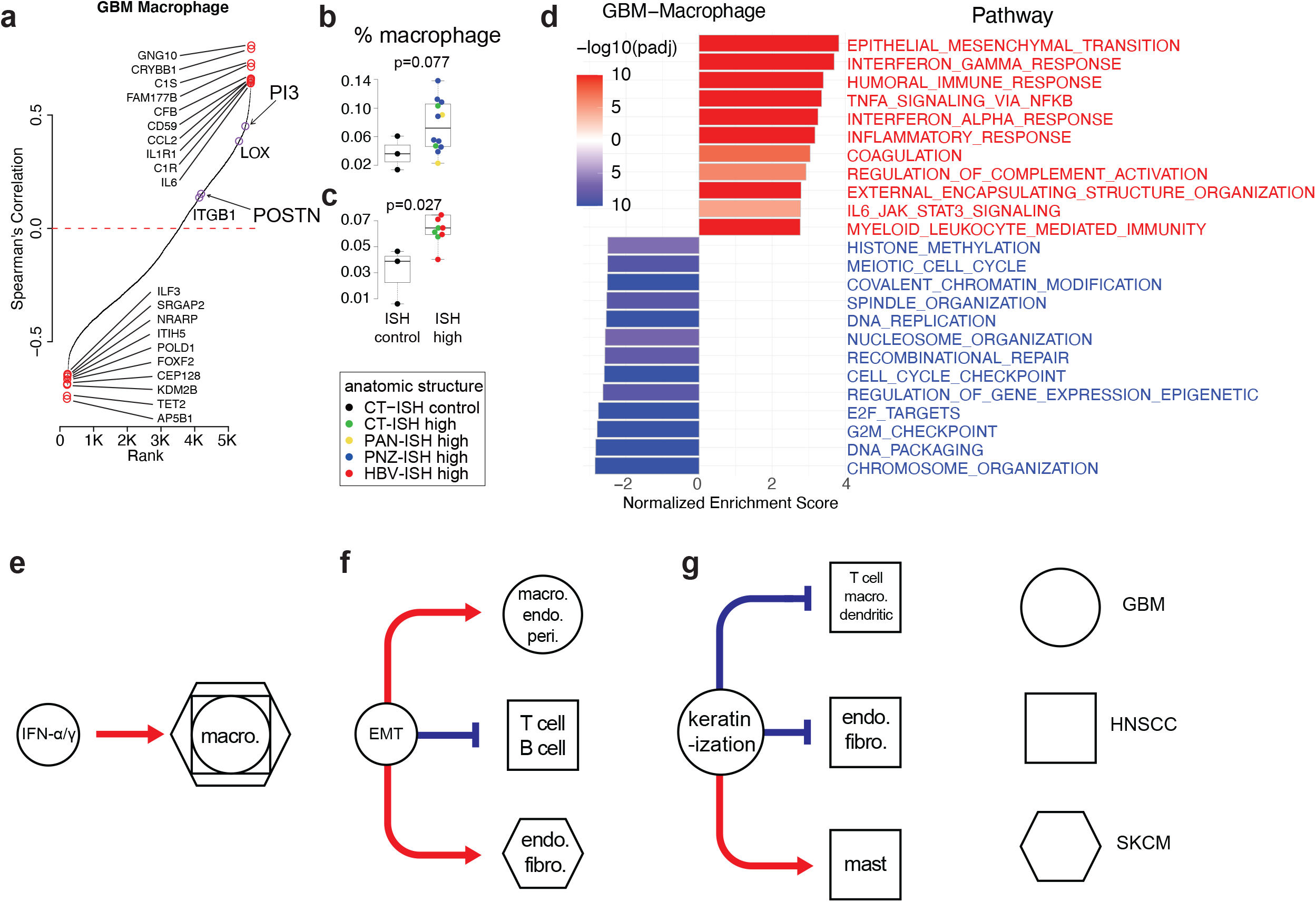
Correlation between malignant cell gene expression and non-malignant cell fraction. **a**) Rank-ordered plot shows Spearman’s rank correlation between gene expression in malignant cells inferred by BayesPrism and macrophage fraction in the TCGA-GBM dataset. Top 10 positive and negative outlier genes are marked in red. Purple circles highlight experimentally validated regulators of macrophage infiltration in GBM, or genes whose expression correlates with macrophage infiltration in IVY GAP. **b**-**c**) Boxplots show the BayesPrism inferred fraction of macrophage infiltration for regions with low (ISH-control) or high (ISH-high) expression of two target genes, PI3 and POSTN. Color indicates anatomic structures associated with the ISH experiments. **d**) Barplot shows the normalized gene set enrichment score of genes ranked by correlation with macrophage cell fraction in GBM, as computed in (**a**). Only semantically non-redundant biological processes from the top 20 most enriched in Supplementary Fig. 21a were selected for visualization. **e**-**g**) Cartoons summarize three patterns of relationship between biological processes and infiltration of non-malignant cell types. Red arrows mark positive correlations, while blue flathead arrows mark negative correlation. Shapes represent the tumor types of the non-malignant cells.

We also identified numerous other correlations with a stronger magnitude. Putative candidate interacting genes include *GNG10, CRYBB1, FAM177B, CP, GLRX, PI3*, genes involved in complement pathway (*C1S, CFB, CD59* and *C1R)*, and cytokine receptors and ligands *(CCL2, IL1R1* and *IL6)*. To validate these new correlations discovered using BayesPrism, we attempted to recapitulate them using an independent bulk RNA-seq dataset composed of 148 laser capture microdissected regions from 34 GBMs from IVY GAP^59^. We asked whether tumor regions in which malignant cells expressed high levels of candidate genes had higher macrophage infiltration. We used BayesPrism to analyze macrophage cell fraction in all 148 IVY GAP samples as the measure of macrophage cell content. Each bulk RNA-seq sample was collected adjacent to sections analyzed by in situ hybridization (ISH) for cancer stem cell markers^59^. Despite limited sample size per marker in the IVY GAP dataset, we observed higher macrophage content in ISH positive sections of *PI3* and *POSTN*, the only two genes passing our filters that were analyzed by at least 10 ISH experiments (**Fig. 3b-c, Supplementary Table 3a**, also see Methods). Thus BayesPrism identified correlations using TCGA that could be recapitulated by intratumoral heterogeneity.

We next extended our analysis to identify candidate interactions in GBM, SKCM, and HNSCC. To summarize the biological processes associated with cell-cell interactions, we performed gene set enrichment analysis^60,61^ using the correlation coefficients between candidate interacting genes and fractions of non-malignant cell types (**Fig. 3d, Supplementary Fig. 21a**). Our analyses revealed several interaction patterns. First, many of the biological processes correlated with the fraction of non-malignant cell types were discovered independently in all three tumor types. For example, interferon gamma / alpha response was positively correlated with macrophages in all three tumor types (**Fig. 3e, Supplementary Fig. 21**). As macrophages secrete interferon gamma and alpha upon activation^62^, our result likely reflects the gene expression response in malignant cells to macrophage infiltration. In addition, we also uncovered tumor type-specific aspects of malignant cell-macrophage interactions. For example, viral and type I interferon response was uniquely associated with macrophages in HNSCC (**Supplementary Fig. 21b**), potentially reflecting HPV infection in a subset of patients. In some cases, biological processes were associated with different cell types in different cancer types. Epithelial-mesenchymal transition (EMT) can play a wide variety of roles in the tumor depending on other factors in the TME^63^. EMT was positively associated with macrophages in GBM, negatively associated with lymphocytes in HNSCC, and positively associated with endothelial cells and fibroblasts in SKCM (**Fig. 3d**,**f, Supplementary Fig. 21**). Finally, some biological processes were exclusively associated with a single tumor type but with multiple cell types in that tumor. One notable example is keratinization in HNSCC, which was negatively associated with multiple non-malignant cells, except for mast cells which had a positive association (**Fig. 3g, Supplementary Fig. 21b**). Tissue stiffness affects a variety of cell phenotypes^64,65^, and thus one interpretation is that keratinization by malignant cells affects tumor stiffness to exclude certain cell types from the TME. These results highlight how BayesPrism can be used to study interactions between biological processes in malignant cells and non-malignant cell infiltration.

### BayesPrism identifies malignant gene programs through embedding learning

Evolutionary pressure pushes malignant cells to optimize for different tasks that are essential for their survival, which is done by regulating sets of co-expressed genes, known as gene programs^66^. These gene programs provide signatures that are characteristic of the heterogeneity between different patients. Gene programs can be inferred from bulk data using methods such as non-negative matrix factorization (NMF)^67,68^ and archetypal analysis^66,69^.

Although these methods have proven broadly useful to define molecular subtypes in a number of different cancers, a persistent weakness is that they often reflect differential infiltration of non-malignant cell types rather than gene expression in malignant cells^68^.

We developed a module in BayesPrism which recovered core malignant gene programs that best explain expression heterogeneity without contamination from non-malignant cell types (**Fig. 4a**). Motivated by recent observations that malignant cells in different tumors are heterogeneous mixtures of functionally distinct cell states^26,31,70^, we modelled each patient as a linear combination of gene programs. BayesPrism infers the weights of each gene in each program and each program in each tumor using the expectation maximization (EM) algorithm, such that the linear combination of all gene programs best approximates malignant cell expression in all patients (**Fig. 4a; see Methods and Supplementary Note 1**). To evaluate whether BayesPrism learned subtypes that reflect intra-tumoral heterogeneity, we identified four gene programs using the GBM28 pseudo-bulk RNA-seq dataset. BayesPrism recovered gene programs that were similar to those recently obtained by factorization of 6,863 malignant cells from 28 patients^31^ (**Fig. 4b**). Moreover, the weights of each gene program learned by BayesPrism were correlated with the fraction of cells in each tumor that represent each of the four major subtypes (**Fig. 4c-d**). Thus, in this particular case, malignant gene programs learned using BayesPrism approximate major malignant cell subpopulations in the presence of a variety of non-malignant cell types.

**Fig. 4.**
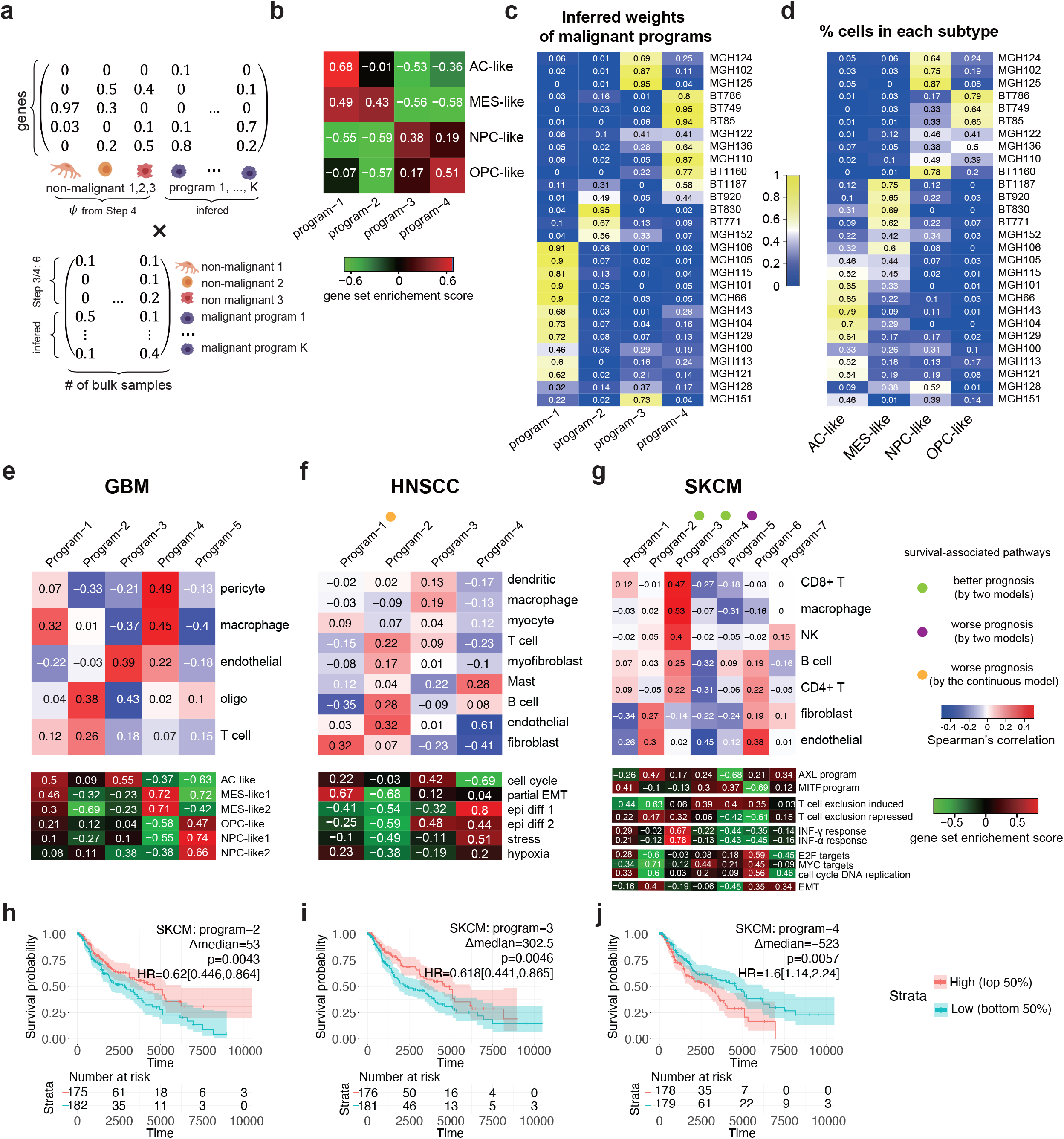
BayesPrism redefines GBM molecular subtypes after excluding expression in non-malignant cells. **a**) Cartoon illustrates the mathematical setup of the embedding learning formulated as a matrix factorization problem. **b**) Heatmap shows the gene set enrichment score for each gene program of malignant cells in GBM28 pseudo-bulk inferred by BayesPrism. Marker genes in each cluster reported by Neftel et al. (2019) are used as the gene sets. **c**) Heatmap shows the inferred weights of each gene program of malignant cells in GBM28. **d**) Heatmap shows the fraction of malignant cells assigned to each cluster in GBM28. **e**-**g**) Top panels show Spearman’s rank correlation between normalized weights of gene programs and the fraction of non-malignant cells in three TCGA tumor types, while bottom panels show their gene set enrichment score for selected gene sets. Colored dots indicate whether the normalized weights of a particular gene program was significantly associated with survival (p<0.05) by both the log-rank test in which the weights were stratified at median (green and purple dots) and a Cox proportional-hazards models in which the weights were modeled as a continuous variable (green and purple dots), or only by the continuous variable model (the orange dot). **h**-**j**) KM plots of gene programs significantly associated with patient survival by two models. Δmedian: median survival time in the high group - median survival time in the low group. P values were computed using the log-rank test. Hazard ratio was defined by high / low, and the 95 percentile confidence interval was shown in the square brackets.

### BayesPrism identifies gene programs in malignant cells from three tumor types

We applied embedding learning to GBM, HNSCC, and SKCM (**Supplementary Fig. 22-24**). We selected the number of gene programs in each tumor type based on the degree of consensus clustering of the deconvolved expression of malignant cells using NMF^71^ (GBM k = 5, HNSCC k = 4, SKCM k = 7; **Supplementary Fig. 22a**). We set the priors of gene programs based on the factors imputed by NMF, thereby incorporating information from the scRNA-seq reference into the embeddings learned by BayesPrism. BayesPrism revealed several programs that were similar to those in previous studies^31,67,68^, including GBM program 3 (classical and AC-like), program 4 (mesenchymal), and program 5 (proneural, OPC, and NPC-like) (**Fig. 4e, Supplementary Fig. 23a**). Notably, we observed that subtypes proposed to reflect contamination of the TME with non-malignant cells were not detected by BayesPrism as gene programs. For instance, the mesenchymal subtype in HNSCC was not strongly associated with any gene program (**Supplementary Fig. 23b**). This subtype was only identified by TCGA bulk analyses, but not by more recent scRNA-seq studies, and was suggested to be an artifact caused by the presence of fibroblasts^28^. This observation suggested that the embedding learning module removed the influence of non-malignant cell types resulting in gene programs that were intrinsic to malignant cells.

We next sought to characterize the biological processes associated with each gene program. We applied gene set enrichment analysis over the inferred expression of malignant cells for bulk samples specifically associated with each program (**Supplementary Fig. 23, Supplementary Fig. 24, Supplementary Table 4, Supplementary Table 5**, see Methods).

In GBM, program 4, a program positively correlated with macrophages, pericytes and endothelial cells (**Fig. 4e**), was associated with EMT, TNF-α signaling via NF-κB, IL-6/JAK/STAT3 signaling, hypoxia, angiogenesis and inflammatory responses. Program 5 was negatively correlated with macrophages (**Fig. 4e**), had the highest cell cycle score, and was strongly enriched for the gene ontology terms related to cell replication. Program 3 was associated with core cellular processes, such as regulation of nucleobase containing compound transport, neuroblast proliferation, mRNA processing, DNA conformational changes, and had a high cell cycle score. We also noted some interferon signaling associations in program 3. Overall, the associations we observed between gene programs 3-5 and biological processes frequently mirrored those reported previously^67^. Programs 1 and 2 had some notable differences from all previously described subtypes. Program 1 was similar to both AC and MES, which agrees with the recently reported stratification of the MES subtype and heterogeneity of the AC subtype in GBM^72^. It was strongly enriched for genes associated with cell respiration, and was negatively associated with endothelial cells. Interestingly, program 2 was positively correlated with oligodendrocytes, was enriched for multiple neuronal processes, and had the lowest cell cycle score. As it constitutes on average 22% of the expression of malignant cells, it is unlikely that this gene program is due to contamination of normal brain tissue. On the contrary, it may reflect the quiescent neuroblast-like malignant populations identified by scRNA-seq^30^. Notably, in several cases, ontology terms enriched in malignant gene programs reflected similar sets of genes as those which were identified using correlations between malignant cell expression and non-malignant cell type fraction (**Supplementary Note 8**).

Several of the gene programs in HNSCC and SKCM showed significant associations with patient survival. Notably, in HNSCC, Program 1 had a negative association with survival (*p* = 0.017, Wald test, **Supplementary Fig. 25a**). Program 1 was uniquely enriched for the partial EMT program (p-EMT) identified by the single cell study^28^ (**Fig. 4f**). Comparing this program to subtypes derived from bulk TCGA tumor profiles, we found that it resembled the basal subtype but not the mesenchymal subtype (**Supplementary Fig. 23b**), consistent with the characterization of the p-EMT by the scRNA-seq study^28^. Top differentially upregulated genes that marked this program also overlap signature genes of p-EMT, including *MMP10, LAMA3, LAMC2, COL17A1*, and *SEMA3C* (**Supplementary Fig. 24b, Supplementary Table 5b**). Additionally, this program was positively correlated with fibroblasts (**Fig. 4f**), which was also concordant with the previous finding by immunohistochemistry^28^. Therefore, the negative association between HNSCC program 1 and survival discovered in our work most likely reflects the same biological processes that explains increased invasiveness of cells expressing p-EMT genes, as well as the positive correlation between p-EMT scores, metastasis, and other adverse pathological features (e.g., lymph node metastases and higher nodal stage and tumor grade)^28^.

In SKCM, multiple gene programs correlated with survival (**Fig. 4h-j, Supplementary Fig. 25b**). Survival-associated gene programs in SKCM had enrichment or depletion for the AXL and MITF gene programs identified by the TCGA bulk analyses, as well as the T cell exclusion programs identified by the scRNA-seq study (**Fig. 4g**). Program 1 and 2 showed the highest MITF and AXL enrichment scores, and were negatively and positively correlated with fibroblasts respectively, which was consistent with the previous analysis on the TCGA dataset based on correlations on marker genes^27^. Program 2 was enriched for EMT, NF-κB, and hypoxia, and depleted of cell cycle genes (**Fig. 4g, Supplementary Fig. 23c**). This program represents a less replicative malignant state due to environmental stress, and was positively correlated with survival (**Fig. 4h**). Programs 3 and 4 were positively and negatively correlated with both immune cells and survival (**Fig. 4g**,**i**,**j**), coinciding with the relationship between immune infiltration and survival in SKCM (**Fig. 2b-c**). Program 3 was enriched for immune process and interferon response, and genes repressed in the T cell exclusion program, including *C4A* and *IFI27* (**Supplementary Fig. 24c, Supplementary Table 5c**), while program 4 was enriched for Myc targets and genes induced in the T cell exclusion program (**Fig. 4g, Supplementary Fig. 23c**).

Taken together, the embedding learning module provides a systematic framework to perform factor analysis by leveraging prior information from scRNA-seq data to remove confounding non-malignant cell types and incorporate information about the cell state of malignant cells.

### Spatial heterogeneity of malignant gene program and non-malignant cell infiltration in GBM

We hypothesized that the relationship between the activation of gene programs in malignant cells and the proportion of non-malignant cell types in the microenvironment display substantial intratumoral spatial heterogeneity. We deconvolved 122 bulk RNA-seq samples microdissected into five structures by IVY GAP^59^: leading edge (LE), infiltrating tumor (IT), cellular tumor (CT), microvascular proliferation (MVP) and pseudopalisading cells around necrosis (PAN) (**Fig. 5a, Supplementary Table 3b**). Notably, the TME of these distinct structures is known to differ in several respects, including blood supply, oxygen level, and immune stress^73^, all of which likely affect both cell type composition and malignant cell states. Deconvolution was conducted using the microwell GBM scRNA-seq dataset^30^, as noted for TCGA (above). Because two of the five structures (LE and IT) correspond to regions with significant amounts of normal brain tissue in which the tumor was just beginning to migrate, we also added primary neurons from a separate source^74^. The addition of neurons into the reference was useful to evaluate the relative quantity of normal brain tissue in each sample. We caution that batch effects between cell types in the reference dataset will tend to systematically overestimate neurons, although the relative rank across samples will be preserved allowing a qualitative comparison between the five structures (**Supplementary Note 9**).

**Fig. 5.**
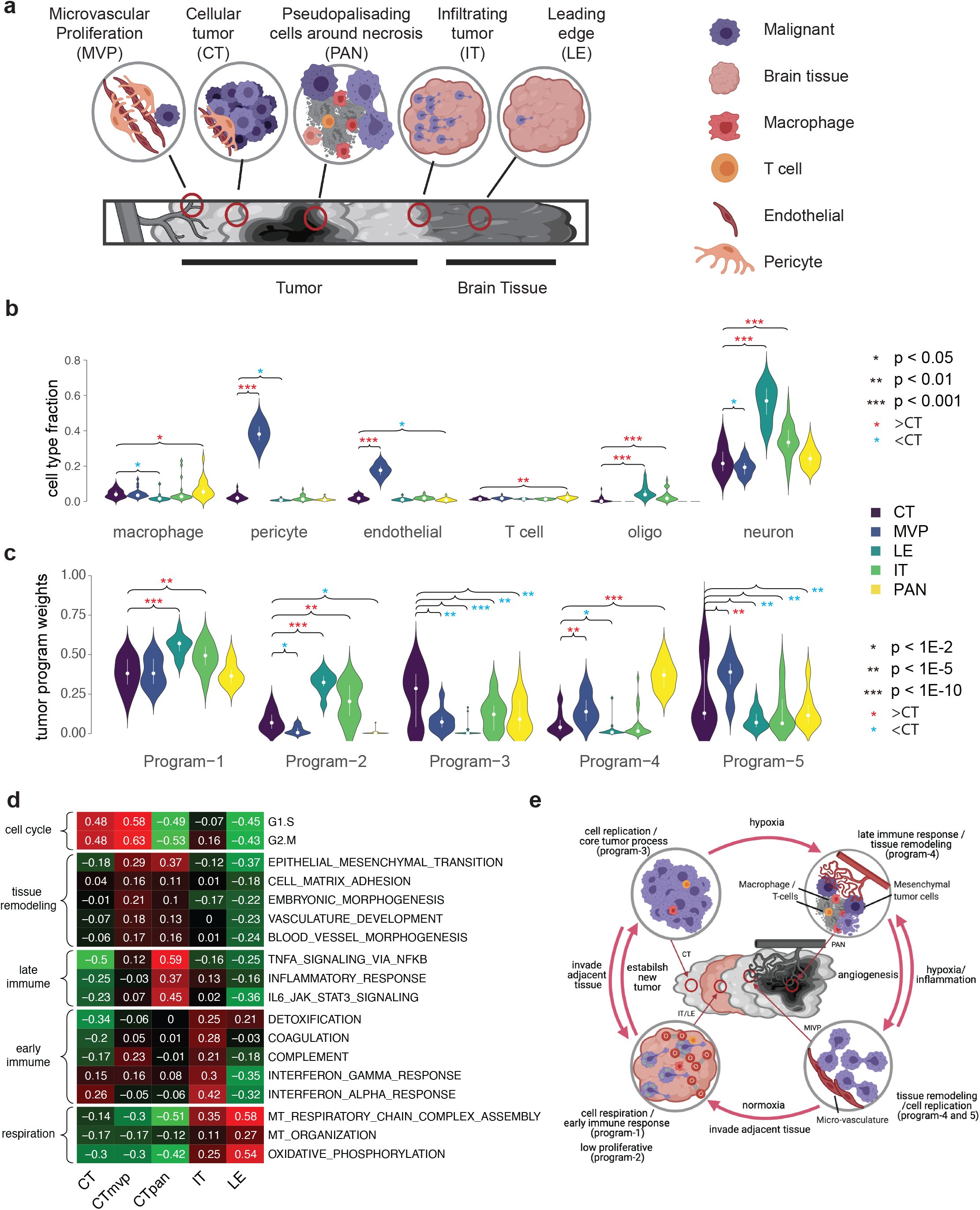
BayesPrism reveals spatial heterogeneity in GBMs. **a**) Cartoon depicts the spatial relationship between anatomic structures present in the IVY GAP samples. **b**-**c**) Violin plot shows the distribution of cell type fractions (**b**) and weights of each gene program learned from TCGA-GBM (**c**) in the inferred expression of malignant cells of each anatomic structure over 122 IVY GAP samples. Median fractions are shown by white dots and upper/lower quartiles are shown by bars. Asterisks mark significant differences between CT and other anatomic structures based on a linear mixed model. **d**) Heatmap shows the gene set enrichment score of each anatomic structure for biological processes selected from the correlation analysis shown in Fig. 3d and Supplementary Fig. 23a. **e**) A model depicting the interaction between gene programs in malignant cells and non-malignant cells in the microenvironment of GBM.

We examined which cell types and gene programs (identified using TCGA, above) were enriched in the anatomical structures investigated by IVY GAP (**Fig. 5b-c**). Notably, PAN regions were enriched for macrophages, T cells and program-4 (mesenchymal), consistent with prior observations^67^. As expected based on the presence of substantial microvasculature, MVP regions were highly enriched for endothelial cells and pericytes. LE and IT were enriched for oligodendrocytes and neurons, which is consistent with these regions being predominantly composed of normal brain tissue as malignant cells invade adjacent tissues. Extending our analysis to gene programs, we found that LE and IT were enriched for programs 1 and 2, CT was enriched for program 3, PAN regions were enriched for program-4, and MVP for programs 4 and 5.

To help interpret the enrichments in the programs obtained by BayesPrism, we next examined the gene set enrichment score in malignant cells (inferred using BayesPrism) within each IVY GAP structure for a subset of biological processes that showed evidence of substantial variation in TCGA-GBM (above) (**Fig. 5d**). We found that CT and MVP were highly proliferative, consistent with their enrichment for program 3 and 5, which were enriched for cell proliferation terms. MVP and PAN were both enriched for tissue remodeling and immune interactions (program 4), while MVP was more angiogenic and PAN was more inflammatory. Both IT and LE were highly respiratory, LE being the most respiratory and the least proliferative, explaining their enrichment for program 1. IT was also enriched for a subset of pro-inflammatory immune processes, most notably for interferon response. Taken together, our analysis shows how BayesPrism was able to link pathways and gene programs with spatial anatomical structures using the IVY GAP dataset.

## Discussion

A large body of literature now provides examples of how non-malignant cells influence malignant cell function^25,63,75^, confirming more than a century of speculation about the critical role of the TME^1^. However, our knowledge remains largely anecdotal and based mostly on work in animal models rather than human subjects. scRNA-seq has recently made it possible to measure both cell types present in the tumor and their gene expression states in a systematic manner^75^. Although scRNA-seq provides the right data modality to chart the various ways in which tumor-microenvironment interactions occur, current studies do not have a large enough sample size to address these questions in a statistically rigorous manner. In parallel, thousands of bulk RNA-seq datasets are now available that provide weak information about the entire cellular milieu in a variety of malignancies^43–45^. Here we leveraged both genomic resources by developing a rigorous statistical modeling strategy and using it to integrate scRNA-seq data from 37 thousand cells over 85 patients and 1,412 bulk RNA-seq samples, providing a new lens into both the cell type and expression in three human cancers.

Our work shows how integrative analysis of single cell and bulk RNA-seq can provide insights into disease progression. Taking GBM as an example, our joint analysis of malignant and non-malignant gene expression integrating spatial heterogeneity and large cohorts of patients in TCGA prompted us to propose a model that links malignant cell states and non-malignant cell infiltration to tumor progression (**Fig. 5e**). As malignant cells grow rapidly, they deplete nutrients in the local environment and may also encounter immune stress, leading to necrosis (**Fig. 5e; top right**). Consistent with this phase, we observed an enrichment of immune cells and mesenchymal program 4, with a higher EMT and lower respiratory activity, in PAN regions (see **Fig. 5b-c**). Malignant cells may activate these tissue remodeling pathways to promote M2 macrophage polarization and angiogenesis. As micro-vascular structure develops, malignant cells proliferate rapidly (**Fig. 5e; bottom right**), which is supported by the high cell cycle score in malignant cells near MVP (**Fig. 5d**). Proliferating cells may spark the invasion of adjacent normal brain tissues, where oxygen supply is ample. As they do so, they change their major task from rapid proliferation to respiration in order to generate stores of ATP necessary to synthesize essential molecular machinery (**Fig. 5e; bottom left**). This result is based on the enrichment of cell respiration pathways and tumor programs in the LE and IT structures (**Fig. 5d**). Finally, having accumulated sufficient cellular machinery and as the local oxygen level decreases, malignant cells then resume rapid proliferation. Taken together, this model illustrates how GBM cells optimize over multiple tasks to reshape and respond to changes in the local microenvironment^66^.

Our model relates known GBM subtypes to interpretable gene programs and their associated microenvironment, providing mechanistic insight into disease progression. We suggest that the classical-like program-3 may reflect an earlier stage of cancer growth where blood supply is ample. As the disease progresses the malignant cells outgrow the local nutrient supply, resulting in hypoxia and necrosis. This proposal explains why classical tumors recur as mesenchymal tumors in longitudinal studies more frequently than the other direction^68,76^.

BayesPrism fills several critical needs in the genomics toolbox. BayesPrism more accurately deconvolves bulk RNA-seq into the proportion of cell types than previous approaches thanks in part to the Bayesian statistical model which accommodates known differences between bulk and scRNA-seq data. Most importantly, BayesPrism is not just a deconvolution algorithm - it jointly models cell types and their average expression, which was crucial for analyses reported here. It is important to note that the gene expression and cell type fractions estimated by BayesPrism represent a mathematically optimal solution given the information from the scRNA-seq reference. In practice the accuracy of BayesPrism can be affected by missing cell states in the reference matrix, which is a general issue for all deconvolution algorithms. The expression of missing cell states in a heterogeneous TME can sometimes deviate from the prior distribution modeled by BayesPrism, resulting in a partial assignment of transcripts from the missing cell states to cell states from other cell types. Caution needs to be taken when performing correlation analysis between the posterior estimates of gene expression and cell type fraction, potentially using similar filters to those introduced here. Nevertheless we speculate that deconvolution of tumor samples will become more accurate as we collect single cell data from more patients, each presumably covering nuances in the transcriptional state. Thus we envision that BayesPrism provides a new type of lens for integrating the ever growing amount of scRNA-seq data with existing large cohorts of bulk RNA-seq data, allowing insights into tumor-microenvironment interactions.

## Acknowledgements

We thank Peter Sims for sharing the cell type annotation, Xin Bing for discussions on statistical inference, Mariano Viapiano for discussions and biological insights, and Tal Nawy for editing the manuscript. Work in this publication was supported by R01-HG009309 (NHGRI) to CGD, U2C-CA288284 (NCI) and U54-CA209975 (NCI) to DP, and P30-CA008748. T.C. thanks the Croucher Foundation for the Croucher Postdoctoral Fellowship (2019). The content is solely the responsibility of the authors and does not necessarily represent the official views of the US National Institutes of Health. Some figures were created with BioRender.com.

## Author Contributions

The project was conceived by TC with input and advice by CGD and DP. TC developed and implemented BayesPrism, conducted all analyses, and wrote the first draft of the manuscript. ZW developed the web portal. CGD supervised all work and edited the paper.

## Methods

### Overview of BayesPrism

A complete mathematical description and justification of BayesPrism is included in **Supplementary Note 1**. Here we provide a brief summary of BayesPrism and its use in this manuscript. The R package of BayesPrism can be downloaded at https://github.com/Danko-Lab/TED.git. The BayesPrism web portal can be accessed at https://dreg.dnasequence.org.

BayesPrism is comprised of two functional modules: (1) a module that infers the cell type fraction and gene expression of each cell type in each bulk RNA-seq sample (**Fig. 1a**), and (2) a module designed to identify commonly occurring malignant gene programs after removing gene expression in non-malignant cells that infiltrate the tumor (**Fig. 4a**). The second module depends on the output of the deconvolution module.

#### Definition of cell types and cell states

BayesPrism models gene expression of each cell type using a multinomial distribution. In reality, however, depending on the granularity of the cell type labels provided by the user, gene expression can be heterogeneous within each cell type, and hence show overdispersion from the multinomial distribution. This can be particularly the case for malignant cells and non-malignant cells, such as M1/M2 macrophages, in the TME, or for differences in cell cycle or copy number variation in malignant cells. To better accommodate heterogeneity in cell states, we use the concept of cell states (or cell subtypes), which can be obtained by further sub-clustering within each heterogeneous cell type. BayesPrism computes the posterior sum over the cell states to obtain the statistics for each cell type.

The following notation will be used upon describing the raw input:

Shared between bulk and scRNA-seq:

- *G* - number of genes

Bulk:

- *N* - number of bulk samples
- *X*_*ng*_ - raw count of the *g*_*th*_ gene in the *n*_*th*_ bulk sample
- *R*_*n*_ - total reads of the *n*_*th*_ bulk sample over *G* genes

scRNA-seq:

- *S* - number of cell states
- *T* - number of cell types
- *C* - number of cells
- *S*_*c*_ - the cell state of the *c*_*th*_ cell
- *T*_*c*_ - the cell type of the *c*_*th*_ cell
- *W*_*cg*_ - raw count (UMI) of the *g*_*th*_ gene in the *c*_*th*_ cell
- *R*_*c*_ - total reads (UMIs) of the *c*_*th*_ cell across *G* genes

#### Constructing the scRNA-seq reference φ

We assume that the raw count of each cell conditional on the cell state follows the multinomial distribution, where ϕ ∈ ℝ^*S*×*G*^ encodes the event probabilities of each gene *g* of cell state *s*.

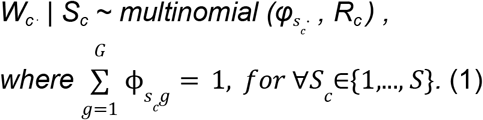

Hence, the maximum likelihood estimate of *φ*_*sg*_ is obtained by summing up read count for cells of cell state *s* and then renormalizing such that the sum across genes is 1:

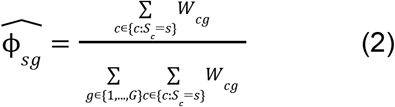

To avoid zero entries in 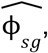, we add a pseudo count to all genes such that after renormalizing to one, the zero entries equals to the user defined pseudo.min (set to 10^−8^ by default).

#### Inferring cell state fraction and gene expression

The following notation will be used to describe this step:

- μ_*ns*_ - estimates of fraction of reads from the *s*_*th*_ cell state and the *n*_*th*_ bulk sample
- *R*_*ns*_ - total reads assigned to the *s*_*th*_ cell state in the *n*_*th*_ bulk sample
- *U*_*nsg*_ - number of reads assigned to the *g*_*th*_ gene of *s*_*th*_ cell state in the *n*_*th*_ bulk sample
- α - the hyper-parameter of dirichlet distribution, set to a small value=10^−8^, to represent a symmetric non-informative and weak prior by default.

We assume that reads from the *n*_*th*_ bulk sample are generated from the following process:

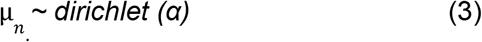

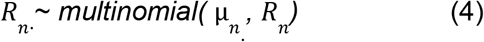

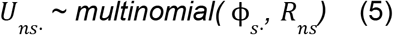

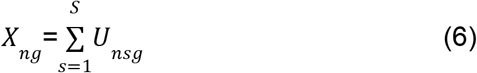

Using Gibbs sampling, we sample from the joint posterior distribution: (see **Supplementary Note 1** for the full derivation).

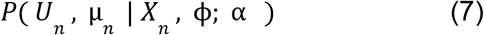

, and simultaneously their marginals:

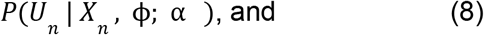

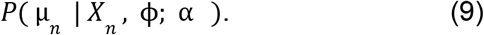

We then compute the posterior means for *U*_*n*_ and μ_*n*_, which we denote 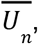, and 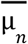.

#### Inferring cell type fraction and gene expression

As cell state is often used to approximate gene expression on a continuous manifold, it may not necessarily correspond to any actual state in the bulk. Therefore, we compute the posterior sum of the fraction and expression of each cell type over multiple cell states within a given cell type. The posterior sum is also of lower variance and more biologically interpretable than that of individual cell states. The following notation will be used to describe this step:

- 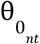-estimates of fraction of reads from the *t*_*th*_ cell type and the *n*_*th*_ bulk sample
- *Z*_*ntg*_ - number of reads assigned to the *g*_*th*_ gene of *t*_*th*_ cell type in the *n*_*th*_ bulk sample.

More formally,

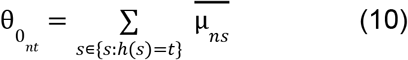

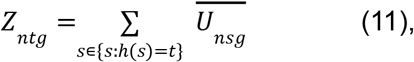

where *h* is a surjective function that maps the cell state index to the cell type index, i.e. from {1,…, *S*}to {1,…, *T*}.

#### Update the cell type fraction estimates

Under situations where 1) non-malignant cell types are of low heterogeneity across bulk samples, and 2) they are presented at substantial fractions, e.g. > 1%, in at least one bulk sample, it is beneficial to leverage the information shared across bulks to improve the estimates of cell type fractions. This is done by first constructing an updated reference matrix *ψ* to replace *φ*, and then re-estimated the posteriors of cell type fraction using *ψ*. As malignant cells are known to exhibit substantial heterogeneity, BayesPrism will not pool information of malignant cells, but instead models a sample-specific *ψ* for malignant cells. The following notation will be used to describe this step:

- 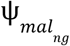-the updated reference of the *g*_*th*_ gene of the malignant cell from the *n*_*th*_ bulk
- 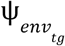-the updated reference of the *g*_*th*_ gene of the *t*_*th*_ non-malignant cell
- σ- a hyper-parameter describing a weak prior on Ψ_*env*_, set to 2 by default to represent a weak non-informative prior.
- 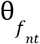 - the updated cell type fractions of the *t*_*th*_ cell type in the *n*_*th*_ bulk sample
- 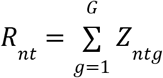 - total reads assigned to the *t*_*th*_ cell type in the *n*_*th*_ bulk sample
- 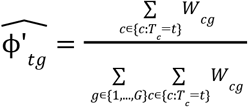- the reference matrix defined for each cell type *t*, similar to (2)

To construct Ψ_*mal*_, we assume that

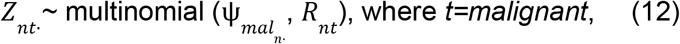

and hence the maximum likelihood estimate of Ψ_*mal*_ is:

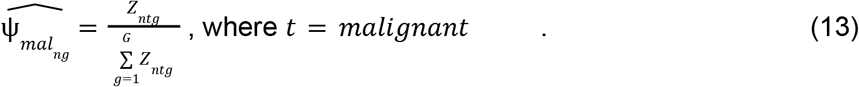

Finally we adjust 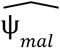 by adding a pseudo-count similar to (2).

To construct Ψ_*env*_, we pool information across all bulk samples. Considering that some non-malignant cells may be present in an extremely low amount in all bulk samples, we put a prior on Ψ_*env*_, such that the estimates of Ψ_*env*_ will be close to ϕ’_*env*_ when there is little information from the bulk. Specifically, we assume the following generative process:

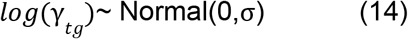

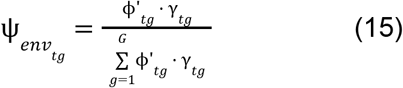

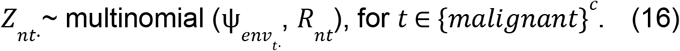

As there is no closed form solution to Ψ_*env*_, we use numerical optimization to obtain the maximum a posterior estimator 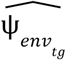. For the *n*_*th*_ bulk sample, we construct a sample specific reference φ_*ntg*_ by concatenating 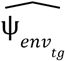 and the *n*_*th*_ row of 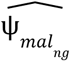. More formally,

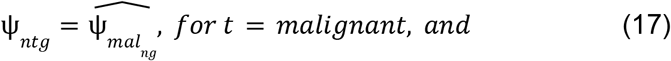

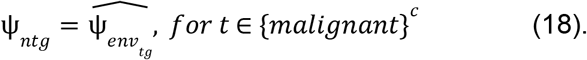

We then use the same generative process as described in (3)-(6), by replacing ϕ with the sample-specific reference Ψ, and derive the posterior similar to (9):

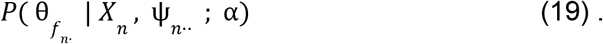

#### The embedding learning module

The second module of BayesPrism was designed to identify gene expression patterns that arise commonly among bulk RNA-seq samples after removing non-malignant cells infiltrating the tumor. BayesPrism learns the latent embeddings, called malignant bases (denoted by *η*), chosen such that their linear combination best approximates gene expression levels in malignant cells. Essentially this module tries to solve the non-negative matrix factorization (NMF) problem:

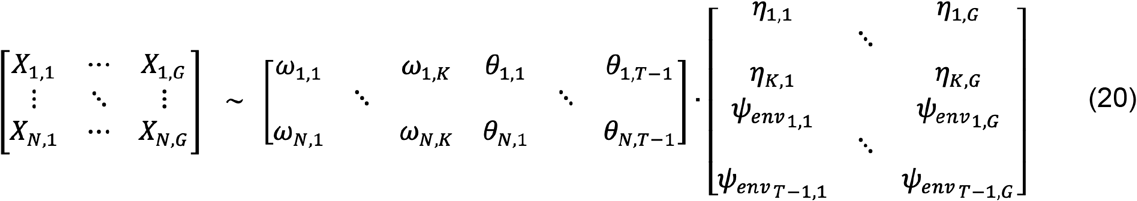

, written in short as *X*∼ υ⋅ζ, with υ∈ ℝ^*N*×*M*^ and ζ∈ ℝ^*M*×*G*^, where *M*=*K*+*T*-1. In (20), θ and Ψ_*env*_ blocks are the non-malignant components from θ and Ψ inferred by the deconvolution module in (19) and (18), and are fixed during inference. ω and ηare latent variables inferred. η has a weak prior over η_0_:

*log*(λ_*kg*_)∼ Normal(0,σ), whereσis set to 2 by default, similar to (14), and

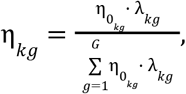

η_0_ represents a prior guess of gene programs of malignant cells. It can be supplied by users based on domain knowledge or by running clustering or NMF on expression inferred for malignant cells in (13). In addition,ωhas a weak scaled non-informative dirichlet prior:

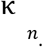 *∼ dirichlet (α)*, where *α* is set to a small value=10^−8^, similar to (3).

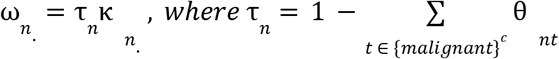

The observed read count of bulk samples, *X*, is then generated as follows:

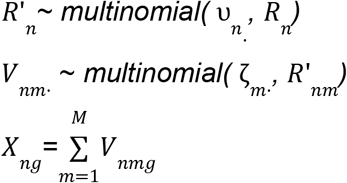

η is inferred using expectation-maximization (EM) algorithm to optimize the log posterior to obtain the *maximum a posterior* (MAP) of η while marginalizing *V* and ω:

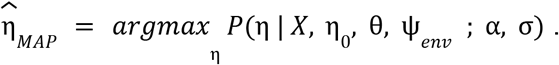

ωis then taken as the posterior mean at the 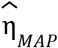:

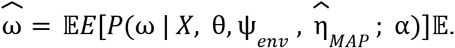

### Deconvolution of bulk RNA-seq using BayesPrism

#### Generating the reference expression profiles from scRNA-seq data

We used reference expression profiles generated from scRNA-seq data to deconvolve the bulk RNA-seq data of the corresponding tumor type. We summed up raw read counts whenever count data is available (for ^29,30^). For data where only TPM normalized data is available (scHNSCC), we summed up TPM normalized reads. To generate the reference profiles of the cell states of non-malignant cells, we first re-clustered the cell types that show substantial inter-tumoral heterogeneity based on the original scRNA-seq studies, including macrophages in GBM; fibroblasts and T cells in HNSCC; macrophages, B cells and CD4 and CD8 T cells in SKCM. To recluster, we normalized the raw reads of each cell using size factors estimated by scran^77^, and then transformed the data using log_2_(Y+0.1) for refGBM8 and log_2_(Y+1) for scSKCM and scHNSCC, with Y being the count normalized by scran. We then removed ribosomal protein coding genes and genes from chrM, chrX and chrY. Next, we performed dimensionality reduction using the randomized singular value decomposition function provided by the rsvd package^78^ at k=30, and clustered the data on the reduced dimension using PhenoGraph using the default parameters^79^. To account for the heterogeneity in malignant cells, we used sub-clusters of malignant cells generated by PhenoGraph^79^ in each individual patient, whenever malignant cells were clustered by the author (refGBM8, 60 subclusters in total for 8 patients). For datasets where malignant cells were not clustered by the original paper (scHNSCC and scSKCM), we defined the malignant cell states using the patient ID. Lastly we aggregated read counts in each cell state to generate the reference profile *φ*. We found the expression of many of the non-coding genes in TCGA were close to zero across all patients, and hence we performed the inference on protein-coding genes when deconvolving TCGA data to speed up downstream analysis. Deconvolution over all genes generated almost identical results (data not shown). In addition, genes on the sex chromosomes were also excluded in the reference to avoid sex-specific transcription states, and ribosomal protein coding and mitochondrial genes were removed to reduce batch effects. Outlier genes in the bulk, defined as those with expression greater than 1% of total reads in over 10% of bulk samples, were excluded from the deconvolution analysis.

#### Choice of hyper-parameters and retrieving the output from BayesPrism

We used the default hyper-parameters of BayesPrism to perform deconvolution: σ=2, α=10^−8^. We used the default setting for Gibbs sampling as follows: chain.length=1000, burn.in=500 and thinning=2 (i.e., we ran an MCMC chain of 1000 samples, discarded the first 500, and used every other sample to estimate parameters of interest). The maximum number of iterations of the conjugate gradient method was set to 10^5^. All cell type fractions used were the initial θ_0_. θ_f_ highly correlates with θ_0_ (R>0.98), and results from all downstream analyses were consistent whether using θ_0_ or θ_f_.

#### Statistical tests for cell type fraction

For comparisons between multiple groups, we used one-way ANOVA using the built-in function “aov” in R. For ANOVA F test statistics that passed our threshold alpha level at 0.05, we used the function TukeyHSD to perform multiple testing-corrected pairwise tests based on the studentized range statistics.

### Embedding learning analysis

To initialize the malignant gene programs (bases), we used the NMF R package^71^ to learn a linear combination that best approximates the normalized expression of malignant cells inferred by the deconvolution module of BayesPrism (res$first.gibbs.res$Zkg.tum.norm). We optimized the number of malignant bases from 2 to 12, and chose the *K* that yielded the best cophenetic score before a significant drop began. We then fixed the *K* and randomly initialized the algorithm 200 times and chose the bases that yielded the minimal residuals. The malignant bases optimized by NMF were then used as inputs for the embedding learning module of BayesPrism. Although BayesPrism does not necessarily require the use of input bases learned using external algorithms such as NMF, and can initializing bases using clustering methods when no user-defined input is used, we found that initializing bases using NMF significantly sped up the convergence of EM, and also facilitated the selection of the number of gene programs *K* when no prior information was provided.

### Performance Benchmarks

#### Benchmarks against other deconvolution tools

We benchmarked BayesPrism against CIBEROSRTx, Bseq-SC, Bisque, SCDC, and MuSiC. Details of the datasets used for the benchmark were listed in **Supplementary Table 1**. Marker genes are required for CIBERSORT-based methods including Bseq-SC and CIBERSORTx (all modes), while they are optional for all other methods including BayesPrism. For CIBERSORTx we used the online portal of CIBERSORTx (https://cibersortx.stanford.edu) to perform all the benchmarking. All parameters were used at default values, except for the “Min. Expression”, which was set to 0 for single cell references to generate a signature matrix, following the author’s recommendations for droplet-based platforms. Quantile normalization is disabled by default following the author’s recommendations for RNA-seq. For all other methods, we used the corresponding R packages. In all non-tumor sample deconvolutions, we used a single batch of scRNA-seq dataset as the reference for Bseq-SC, SCDC, and split the scRNA-seq dataset equally into two batches for each cell type for Bisque and MuSiC, as multiple batches of single cell references are required by these methods. When benchmarking Bisque we used the “ReferenceBasedDecomposition” and disabled “use.overlap”, as we did not have samples with matched scRNA-seq and bulk RNA-seq. For benchmarking deconvolution of tumor samples (GBM28 pseudo-bulk), we leveraged the information of individual patients from scRNA-seq reference (GBM8) to label biological replicates whenever possible. This includes Bseq-SC, Bisque and MuSiC. Malignant cells in each patient were used as an individual cell type. As only one patient in GBM8 contains T cells, MuSic was unable to model its variance, resulting in the missing T cell from its scRNA-seq reference. To circumvent this, we split the T cells equally into two batches.

#### Gene filtering and feature selection

When deconvolving genes without markers, ribosomal and mitochondrial genes, were removed in all benchmarks. Genes on chromosome X and Y were also removed to prevent sex-specific differences between the scRNA-seq reference and the pseudo-bulk sample. To speed up computation, we removed lowly expressed genes, by selecting genes expressed in at least 5 cells. In addition, outlier genes, defined as genes that show >1% of total reads (or normalized reads if only TPM data is available) in at least one bulk sample were removed, unless otherwise specified below. In all pseudo-bulk analysis, we defined the ground truth as fractions of total reads over all annotated genes in each cell type. We benchmarked this metric against other deconvolution tools, as the fraction of reads is proportional to the total fraction of cell counts, and hence the cell type level correlation will not be affected.

When benchmarking deconvolution using markers, we use the same set of markers generated by CIBERSORTx when applicable to ensure a fair comparison between all methods (**Fig. 1c-f; Supplementary Fig. 2**,**3**, labeled as “method name, w/ marker”). The BayesPrism package also provides options for the use of marker genes to combat cases where significant batch effects exist, e.g. when using ribosomal-depleted RNA or statistical assumptions are possibly violated, e.g. when using references collected from unmatched samples. The implementation was based on the findMarker function implemented by the scran package. Briefly, scRNA-seq reference was normalized by the median library size followed by log_2_(X+0.1) transformation. There are two types of markers defined by the findMarker function, the “all marker” (genes that are significantly differentially transcribed between one cell type and all other cell types) and “any marker” (those that are significantly differentially transcribed between one cell type and any other cell type). Empirically, we found that all markers provided the strongest robustness to batch effects as they are closer to the exclusively expressed marker genes (**Supplementary Note 3**). Statistical significance was calculated based on t-tests, and only genes upregulated in each cell type were used to define markers.

#### Linear multiplicative noise model (Supplementary Fig. 2)

We used the scRNA-seq of PBMCs collected from the first donor using 10x Chromium (v2) A^80^, as the reference, and we simulated 200 pseudo-bulk RNA-seq samples from the same dataset used to build the reference. The cell type fractions were drawn from a symmetric dirichlet distribution (α=1), and the cell count of each cell type was sampled from a multinomial distribution with the total cell number equal to the cell number in the original batch (N=3222). Cells were then sampled with replacement according to the simulated cell count of each cell type, and then aggregated by summing up the reads over the sampled single cells to make pseudo-bulks. In the simulation, no outliers from the bulk were removed.

To generate noise to the pseudo-bulks, we simulated a zero-centered log-normally distributed fold change at one particular noise level σ independently and identically distributed for each gene, which generated a vector of length equal to the number of genes. To mimic the real biological batch effects, we penalized extreme fold changes that resulted in unrealistic expression values, which was particularly frequent at high σ levels. This was done by sampling the fold change vector 10000 times and choosing the one that induced the minimal change to the total expression as measured by elemental-wise multiplying the reference expression with the fold change vector. The chosen fold change vector was then elemental-wise multiplied with the pseudo-bulks which were then rounded up to the nearest integers.

#### Cross-platform deconvolution using pseudo-bulk RNA-seq from non-tumor samples (Supplementary Fig. 3)

For PBMC data, we used the 10x Chromium (v2) dataset collected from the second donor as the reference to deconvolve pseudo-bulks generated by the Smart-seq2 from the first donor in the original paper^80^. For mouse cortex data we used the sci-RNA-seq dataset collected from the second mouse as the reference to deconvolve pseudo-bulks generated by the Smart-seq2 from the first mouse in the original paper^80^. The choice of these datasets is to represent the strongest batch effect based on the correlation shown in the Supplementary Fig. 4 of the paper by Ding et al.^80^.

#### Single-platform leave-one-out deconvolution using pseudo-bulk RNA-seq from HNSCC, SKCM (Supplementary Fig. 4) and OV (Supplementary Fig. 5)

We generated a “pseudo-bulk” RNA-seq dataset from one patient, and asked how accurately BayesPrism deconvolved expression using the reference constructed from the remaining patients. All parameters of BayesPrism and CIBERSORTx were at default. Batch effect correction was disabled for leave-one-out tests. Two cell types, “-Fibroblast” and “myocyte”, in the HNSCC scRNA-seq dataset were of very low cell number <20 cells, and only showed up in a small subset of patients (N=4), which may lead to unreliable estimates of correlation coefficients. Therefore, we excluded them during the leave-one-out test. Benchmarking the expression inference could not be done, as CIBERSORTx requires the number of mixtures to be greater than the number of reference components.

As observed with GBM, BayesPrism consistently estimated cellular proportions that were more accurate to the true values than CIBERSORTx (**Supplementary Fig. 4a-d**). As the leave-one-out test data were generated from the same sequencing platforms and processed by a uniform pipeline, they represent minimal technical batch effects, and hence the superiority in performance of BayesPrism mainly reflects its ability to account for the deviation in the reference caused by inter-tumoral biological variation.

#### Cross-platform deconvolution using pseudo-bulk RNA-seq from GBM samples (Fig. 1c-d; Supplementary Fig. 6, 8a, 9, 10, 11)

We analyzed two glioblastoma multiforme (GBM) datasets collected from different patients using different scRNA-seq platforms to represent a mixture of technical and biological variation. One scRNA-seq reference analyzed 23,793 cells from 8 patients using a microwell-based platform^30^ (GBM8), which sequenced tag clusters near the 3’ end of polyadenylated genes, similar to other high-throughput scRNA-seq methods (e.g., Drop-seq, 10x genomics, etc). A second scRNA-seq dataset was available which sequenced 7,930 cells from 28 patients using the SMART-Seq2 platform^31^ (GBM28), which sequenced full length mRNA transcripts to a high read depth in each cell, similar to most bulk RNA-seq datasets, and hence also mimicked differences between single cell and bulk.

We generated “pseudo-bulk” RNA-seq datasets from GBM28 by 1) aggreagating scRNA-seq counts for each patients (N=28), and 2) 1,350 pseudo-bulk RNA-seq samples by sampling random proportions of each cell type from a symmetric dirichlet distribution (a=1) and then aggregating reads from susampled cells using GBM28 with replacement. This dataset aims at testing BayesPrism across a wider range of different tumor compositions. The malignant cells in each simulated pseudo-bulk were restricted to an individual patient, while the non-malignant cells were drawn from all patients regardless of patient ID. 50 pseudo-bulk RNA-seq samples were simulated for each patient among 27 out of 28 GBM patients, with one sample BT1187 excluded due to only having 8 malignant cells. As raw data were TPM normalized, we rounded up the counts after summing them up across each cell. For comparison with CIBEROSRTx, we downsampled ⅕ of the 1,350 simulated samples, so that it could complete within a reasonable amount of wall time (∼30hrs) (see **Supplementary Note 2** for details). We disabled batch correction when imputing gene expression for CIBERSORTx as both S mode and B mode batch correction resulted in worse performance than when no correction was used.

To construct the scRNA-seq reference, we treat each tumor subcluster as individual cell states (N=60), while summarizing each non-malignant cell type as a single reference (N=5). When benchmarking CBIERSORTx, we used the same cell state labeling by denoting each column of the single cell reference matrix as a cell phenotype, and then summing over weights across cell states within the malignant cells. For gene expression imputation in **Fig. 1h** and **Supplementary Fig. 10**, we summed up the imputed expression values across 60 malignant cell states to get the expression of malignant cells in each sample. Only 53 genes are imputable across all malignant states references by the high resolution mode (by excluding the “1” and ‘‘NA’’ values in the CIBERSORTxHiRes_job1_PJ0XX-tumor-X_Window140.txt). Correlation coefficients in **Supplementary Fig. 10** were computed on these 53 genes for four different approaches.

To show BayesPrism also infers the expression of non-malignant cells in addition to malignant cells, we generated an additional set of references and pseudo-bulks using GBM8 and GBM28, by incorporating heterogeneity in macrophages into our simulation (**Supplementary Fig. 13**). We first clustered macrophages found in GBM28 and GBM8. We processed the scRNA-seq of macrophages in each dataset as follows. As GBM28 data is TPM normalized, we skipped the normalization step, and log_2_ transformed the data followed by removal of ribosomal protein coding and mitochondrial genes, and genes on chromosome Y. We then filtered out genes expressed in less than 10 cells. We performed dimensionality reduction using the rsvd package, using parameters k= 20, p= 15, q= 3. Phenograph was then used to cluster the macrophages over the 20 PCs imputed by SVD using the default parameter at K=30. Phenograph yielded 10 clusters for macrophages in GBM28. Similarly for macrophages in GBM8, we performed transformation, gene filtering, dimensionality reduction and clustering using the same methods and parameters as described above. We used a medium library size normalization step followed by log_2_ transformation and clustering. Phenograph yielded 11 clusters for macrophages in GBM8. For each patient (N=27) and each macrophage cluster (N=10) in GBM28, we simulated 5 pseudo-bulk datasets, constituting 1,350 samples in total. The scheme of simulation was the same as for malignant cells, above. When deconvolving pseudo-bulks, we used 60 malignant states and 11 macrophage states. The cluster purity was calculated using the “purity” function from the NMF package.

#### Real bulk RNA-seq of human whole blood with ground truth measured by flow-cytometry (Fig.1e-f)

To test the performance of BayesPrism on real bulk RNA-seq, we deconvolved 12 human whole blood samples for which the cell type composition was known using flow-cytometry. We used the same PBMC RNA-seq dataset from the CIBERSORTx paper as the reference, which was obtained from a patient with non-small cell lung cancer (NSCLC) using 10x Genomics Chromium v2 (3’ assay)^35^. The bulk PBMC dataset and scRNA-seq reference were mismatched, as the bulk RNA-seq reference was performed on whole blood and the scRNA-seq reference with PBMCs. Neutrophils present in high abundance in the whole blood sample were not represented in the reference because neutrophils are polynucleated and do not isolate with PBMCs. Missing neutrophils may inflate the fraction of other myeloid^81^ cell types that have similar expression. Thus, we inferred the proportion over a total myeloid population and used the combined fraction of monocytes and neutrophils as the ground truth in all analyses. The bulk RNA-seq of human whole blood and the scRNA-seq reference of PBMCs from non-small cell lung cancer patients were downloaded from the CIBERSORTx website at: https://cibersortx.stanford.edu/download.php. As only the S mode of CIBERSORTx produced accurate results, as shown by the authors, we did not benchmark against the uncorrected and B mode.

#### Statistical test on cell type level Pearson correlation coefficients

To test the difference in the collection of cell type level Pearson correlation coefficients between BayesPrism and other methods, we first applied the Fisher s Z-transformation to the Pearson correlation coefficient *r* (sample correlation coefficient) of each cell type using the following formula:

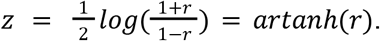

Fisher has shown that if the joint distribution of random variables *X* and *Y* is bivariate normal with correlation *ρ* (population correlation coefficient), then *z* is approximately normally distributed with mean 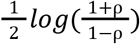 and standard error 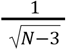, where *N* is the sample size^82,83^. As a result, the difference in r of a particular cell type *c* between method i and j, is normally distributed with mean Δ*z*_*c*(*i,j*)_ = *z*_*c,i*_ – *z*_*c,j*_, and variance 2/ (*N* – 3), and the mean of difference in *z* over *K* cell types between is normally distributed with mean 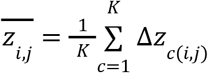, and variance 2/ (*K* × (*N* – 3)). We then performed one-sided z test with 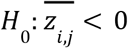, with i being BayesPrism, and j being other methods benchmarked.

### Gene set enrichment analysis

To visualize the magnitude of the enrichment of a selected set of gene sets, such as the inferred embedding in **Fig. 4b, e-g**, and **Supplementary Fig. 23**, the mean of inferred expression of malignant cells in each anatomical structure in **Fig. 5d**, we computed the gene set enrichment scores by performing the gene set variation analysis using the GSVA R package^84^. To compute the statistical significance of gene set enrichment of the correlation coefficient in **Fig. 3d, Supplementary Fig. 21**, and Wald statistics of differential gene expression in **Supplementary Table 4**, we performed gene set enrichment analysis using the fgsea R package^61^.

Details of the use of gene sets are as follows. In **Fig. 4b, e-g**, we used the marker genes of each subtype. In **Fig. 3d, Fig. 5d, Supplementary Fig. 21** and **Supplementary Table 4**, we used the gene sets from The Molecular Signatures Database (MSigDB v7.4)^60^. In **Supplementary Fig. 23** we used both subtype markers and MSigDB. Two gene sets from MSigDB were selected to be tested, the Hallmark (h.all.v7.4; see ref^85^) and the GO biological process (c5.go.bp.v7.4, see ref^86,87^).

Details of fgsea analysis in **Supplementary Table 4** are as follows. To prepare for the input of differential expression analysis, we first ranked the tumor samples according to the normalized weights of gene programs in each sample (summing over weights of malignant programs equals one). We selected the top N/K samples, with N being the number of bulk samples and K being the number of pathways. We removed the samples selected by multiple pathways to define a set of core tumor samples representing each gene program. We performed differential expression analysis between samples by comparing the inferred expression profile of malignant cells in the core samples of each gene program with that of the core samples of the remaining gene programs using DESeq2^88^, and used the Wald test statistics as the input for fgsea. The results of differentially expressed genes were shown in **Supplementary Table 5**.

### Defining Malignant Intrinsic Correlative Genes

When computing the correlation between the inferred gene expression in malignant cells and the proportion of non-malignant query cell types in **Fig. 3a** and **Supplementary Fig. 21**, there is a possibility that some of the genes might have falsely high correlation due to expression in the query cell type. This can happen when the cell states of the query cell types in the bulk are not comprehensively represented by the scRNA-seq, due to the heterogeneity of query cell types. To filter out false correlations and enrich for intrinsic genes expressed by malignant cells for the gene set enrichment analysis, we developed two filters: (1) the *malignant intrinsic filter* that selects genes that are significantly higher expressed in at least one malignant cell state compared to all non-malignant cell types / states in the scRNA-seq reference, and (2) the *regress-out filter* that selects genes that show association with the malignant cells unexplained by the expression of the query cell types. Users can customize the selection of filters and their cutoffs based on the purpose of their analysis to trade off false positives against false negatives. The details of two filters are described as follows.

#### The malignant intrinsic filter

To start with, we first normalized the raw count in each single cell using scran. This is only done for datasets refGBM8 and scSKCM, which are on the raw count scale. We then performed log2(Y+0.1) for refGBM8, log2(TPM+10) for scHNSCC, and log2(Y+1) for scSKCM, with Y being the normalized count. We used a one-sided t test (higher expression in malignant cells), and then for each malignant cell state we take the maximum p value between it and every other non-malignant cell state (refGBM8) / cell type (scSKCM and scHNSCC). We tested differential expression at the cell type level for scSKCM and scHNSCC, because SMART-seq2 has significantly fewer cell types than 10x / microwell-based platforms, which would reduce statistical power if the test was performed on the level of each non-malignant cell state. Lastly, we selected the gene if the maximum p value is less than 0.01 in at least one malignant cell state.

#### The regress-out filter

In the case of incomplete representation of non-malignant query cell states in the scRNA-seq reference, portions of reads from genes expressed higher in the non-malignant query cells may also get assigned to the malignant cells and over all non-malignant cell types, potentially causing false positive associations between their inferred expression in malignant cells and the proportion of query cell types. Likewise, it can also cause false negative associations for missing cell states with lower expression in the scRNA-seq reference. To account for these technical biases, we may take advantage of the Bayesian property of BayesPrism, in which the estimates of gene expression of one particular cell type will drift from the prior (scRNA-seq) to the posterior (bulk) as the cell type fraction increases. We developed a statistical framework by performing a likelihood ratio test on the null hypothesis of β_0(k,g)_=0 in the linear model:

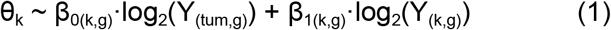

where each random variable has N observations (N=number of bulk samples). θ_k_ is the proportion of query cell type k. For each n, Y_n(tum,g)_ is Z_n(tum,g)_ normalized by the norm.to.one function, such that, 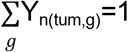, and min(Y_n(tum,g)_)=10^−8^ (the default pseudo.min) for ∀*n*. Likewise Y_(k,g)_ is the normalized expression value for non-malignant query cells. In theory, when the query cell type is presented with non-zero fractions, we can directly apply the norm.to.one function on Z_n(k,g)_ to get Y_n(k,g)_. In reality, however, unlike malignant cells, non-malignant cells can be absent or presented in an extremely low fraction. In those cases, Z_n(k,g)_ will be close to zero for ∀g, and hence directly normalizing Z_n(k,g)_ to get Y_n(k,g)_ will yield unstable estimates. To circumvent unstable estimates due to this technical reason, we developed a maximum a posterior estimator for Y_n(k,g)_ by modeling a prior distribution over Y_n(k,g)_ as follows.

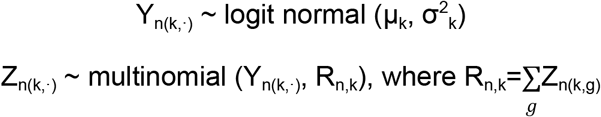

μ_k_∈ℝ^G^, and σ^2^_k_∈ℝ^G^ (diagonal covariance) are hyper-parameters of the model, fitted using a subset of samples index by the set N_0(k)_, in which Y_n(k,)_ has higher Spearman correlation with the posterior sum 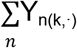 than the prior φ_k_. This strategy is to parameterize μ_k_ and σ^2^_k_ using samples with higher fractions of the query cell type without the need to manually define a cutoff, as samples with higher cell proportion tend to be more similar to the posterior than to the prior. We then set

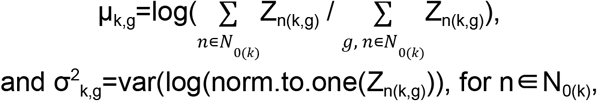

where var is the sample variance over n. As there is no closed form solution, we used the L-BFGS-B optimizer^89^ to maximize the posterior over Y_n(k,g)_. For genes with zero expression in all samples, we excluded them from the estimation of Y_(k,g)_, and replaced the linear model in (1) with θ_k_ ∼ β_0(k,g)_ · log_2_(Y_(tum,g)_), as genes of zero expression in the query cell type across all samples cannot explain the variance of the response variable.

### Choosing cell type marker genes for correlation analysis

In **Supplementary Fig. 19**, we computed the Pearson s correlation coefficient between the variance-stabilized transformed expression over a set of marker genes with the cell type fractions. Marker genes for CD4+ and CD8+ T cell and monocytes were derived from the LM6 matrix from the CIBERSORT website (https://cibersort.stanford.edu/download.php), which were based on GSE60424^90^, by assigning each gene to the cell type with the maximum expression value. Markers for oligodendrocytes, endothelial cells, pericytes, and microglias in normal brains were derived from the gene list generated by Lake et al. using normal brain snDrop-seq^91^. Only marker genes uniquely assigned to each cell type were used for the plot.

### Analysis of M1/M2 macrophage score across three tumor types

We first selected tumor samples with inferred macrophage fractions (by BayesPrism) greater than 5%. In total, 127 GBMs, 26 HNSCCs, and 225 SKCM samples were selected by this criterion. We then applied the DESeq2 variance-stabilizing transformation^88^ over the inferred macrophage gene expression profiles. We computed the Pearson correlation coefficient between the inferred macrophage expression and the log_2_ transformed expression profile of M1 and M2 macrophages from LM22 over a set of M1/M2 marker genes. The marker genes were defined as genes with the highest expression in M1/M2 macrophages across all cell types included by the LM22 reference matrix^34^.

### Analysis of anatomically resolved transcriptomics data from IVY GAP

Anonymized BAM files for each sample were downloaded from glioblastoma.alleninstitute.org, and raw counts for each gene were obtained using featureCounts^92^ using the GENCODE annotation v24lift37.

To test the statistical significance in the mean of cell type fractions (**Fig. 5b**) and gene program weights (**Fig. 5c**, obtained by applying BayesPrism to “deconvolve” the inferred expression of malignant cells using the pathway profile inferred from TCGA as the reference) across multiple anatomic structures while taking account of the multiple biological replicates of each patient, we fit a linear mixed model using the lme function from the R package nlme^93^ with a random intercept. We modeled anatomic structures as the fixed effects and patient IDs as random effects. The ground level was set to the cellular tumor (CT). We maximized the log-likelihood function by setting the method as “ML”. We used “optim” as the optimizer.

To validate candidate genes from the correlation analysis in **Fig. 3a**, we selected genes profiled in at least 10 samples from the cancer stem cells RNA-seq study of the IVY GAP (**Supplementary Table 3a**), resulting in five genes left. Two genes out of five, PI3 and POSTN, also passed the filters used to define the malignant intrinsic correlative genes, and were characterized in **Fig. 3b-c**. The p value of the regress-out filter for gene POSTN is 0.02. Although it did not pass the alpha value we used (=0.01), we still included it for visualization and validation purposes.

### Survival analysis

To avoid known clinical or genetic factors that have a strong influence on patient survival from confounding our survival analysis, we focused on the population with the greatest sample size after conditional on known confounders. These included primary IDH-1 wildtype tumors for GBM, primary HNSCC, and metastatic SKCM. We also attempted to control for HPV status in HNSCC. Although only 72 of 500 samples were annotated for HPV, we nevertheless reproduced consistent trends in a small cohort of 56 HPV negative patients (**Supplementary Fig. 17b**). Two methods have been used to test the association between survival and the feature of interest. 1) Patients were stratified into high and low groups based on the median value of the feature of interest, e.g. normalized weights of malignant gene programs or non-malignant cell fractions, and then computed the hazard ratio by fitting a Cox proportional-hazards regression model for a categorical variable denoting patient groups. 2)

The features of interest were modeled as a continuous variable by the Cox proportional-hazards model, in which we derived the statistical significance using the Wald test and examined the proportional hazards assumption using the chi-squared test for scaled Schoenfeld residuals, i.e. whether the Schoenfeld residuals were independent of time.

### Dataset used

| Dataset name | Normalization | # of cells | # of patient (scRNA-seq) / # of samples (bulk) | Clustered tumor cells | Accession ID |
| --- | --- | --- | --- | --- | --- |
| Tumor scRNA-seq |  |  |  |  |  |
| refGBM8 | Raw (UMI) | 23793 | 8 | YES | GSE103224 |
| GBM28 | TPM | 7930 | 28 | NO | GSE131928 |
| scSKCM | Raw | 6879 | 31 | NO | GSE115978 |
| scHNSCC | TPM | 5902 | 18 | NO | GSE103322 |
| scOV | TPM | 9428 (10x) | 6 | YES | GSE146026 |
| Normal scRNA-seq |  |  |  |  |  |
| PBMC 1 - 10x Chromium (v2) A | Raw (UMI) | 3222 | 1 | NA | GSE132044 |
| PBMC 2 - 10x Chromium (v2) | Raw (UMI) | 3362 | 1 | NA | GSE132044 |
| PBMC 1 - Smart-seq2 | Raw | 253 | 1 | NA | GSE132044 |
| PBMC from NSCLC patients | Raw (UMI) (converted from CPM) | 1054 | 1 | NA | https://cibersortx.stanford.edu/download.php |
| Epileptic adult human brain | Raw | 466 | 1 | NA | GSE67835 |
| Normal mouse cortex single nucleus-seq |  |  |  |  |  |
| Cortex2<br>sci-RNA-seq | Raw (UMI) | 3791 | 1 | NA | GSE132044 |
| Cortex1<br>Smart-seq2 | Raw | 295 | 1 | NA | GSE132044 |
| TCGA bulk |  |  |  |  |  |
| TCGA-GBM | Raw | NA | 169 | NA | https://portal.gdc.cancer.gov |
| TCGA-SKCM | Raw | NA | 471 | NA |  |
| TCGA-HNSC | Raw | NA | 502 | NA |  |
| Normal bulk |  |  |  |  |  |
| Whole blood<br>bulk (with<br>flow-sorted<br>ground truth) | TPM | NA | 12 | NA | https://cibersortx.stanford.edu/download.php |
| IVY GAP |  |  |  |  |  |
| IVY Anatomic<br>Structures<br>RNA-Seq | Raw | NA | 122 samples<br>across 10<br>tumors | NA | https://glioblastoma.alleninstitute.org |
| IVY Cancer<br>Stem Cells<br>RNA-Seq | Raw | NA | 148 samples<br>across 34<br>tumors | NA |  |

