## Supplementary Figures for "Bayesian cell-type deconvolution and gene expression inference reveals tumor-microenvironment interactions"

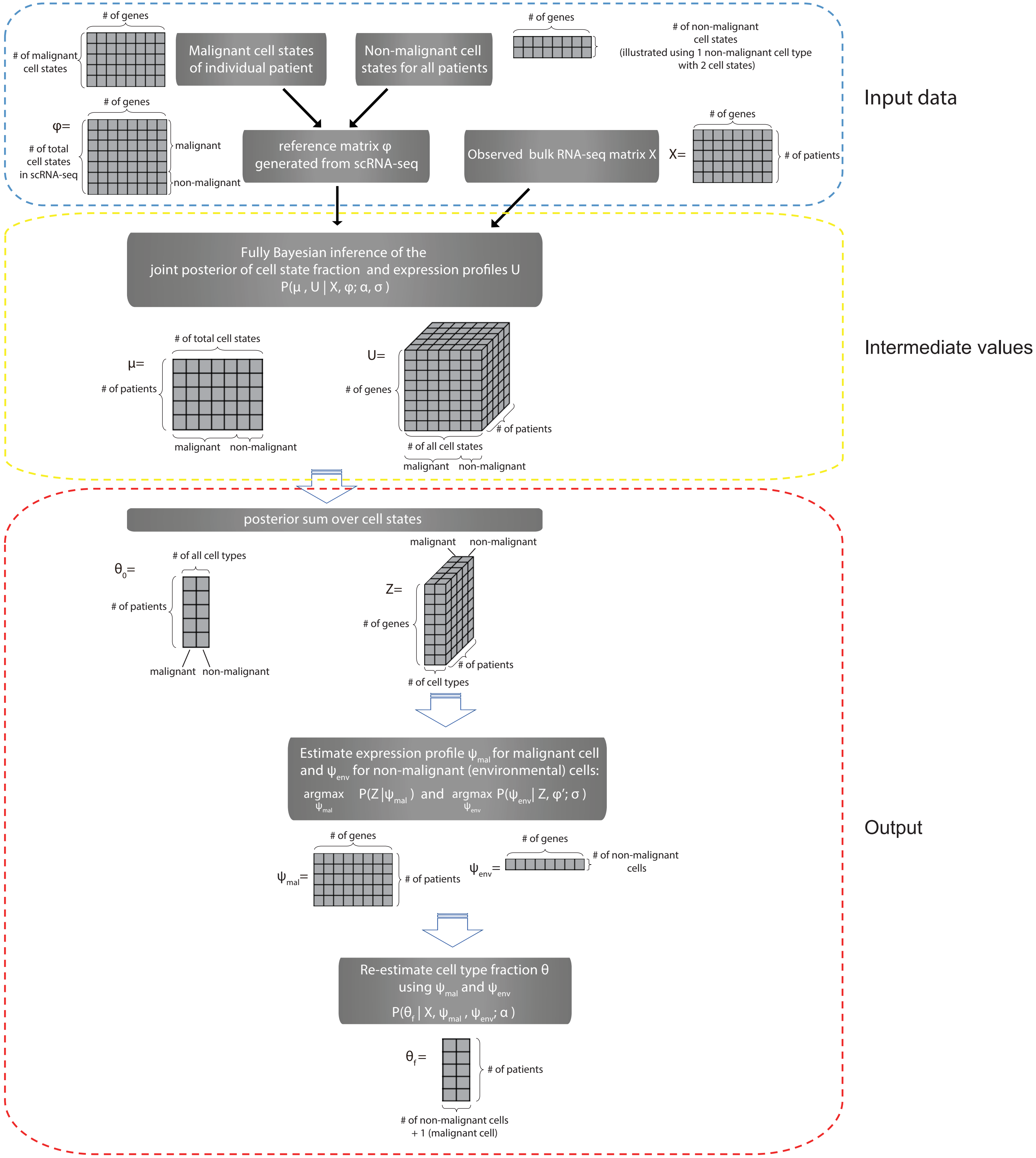


**Fig. S1 | A detailed algorithmist flow of the deconvolution module of BayesPrism.** Gray grids show the dimension of the variables used or inferred in each step.

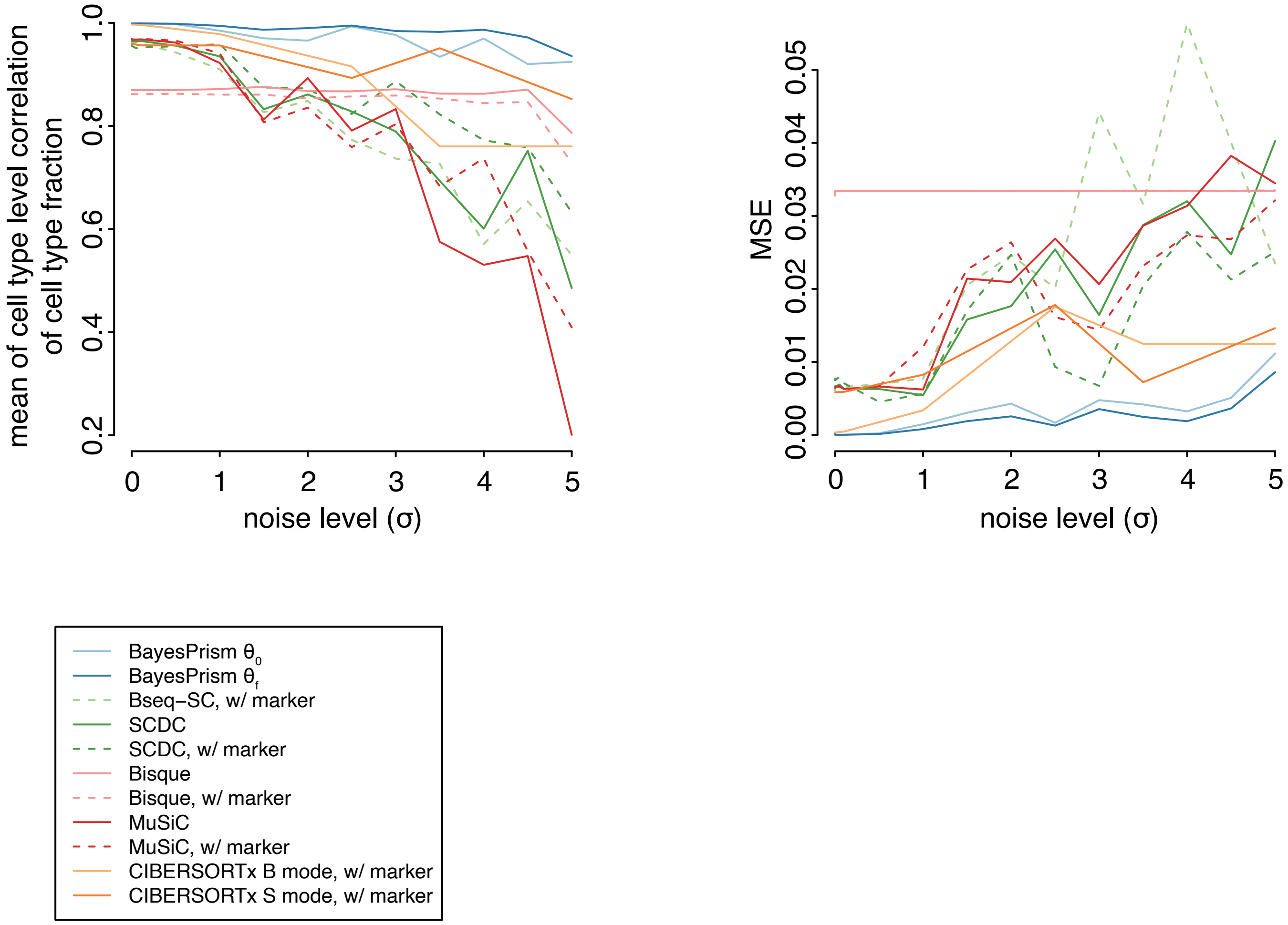


**Fig. S2 | Comparison between BayesPrism and other deconvolution methods using simulated noise.** Line plots show the mean of cell type-level Pearson's correlation coefficient of cell type fraction (left) and MSE (right) as a function of the noise level.

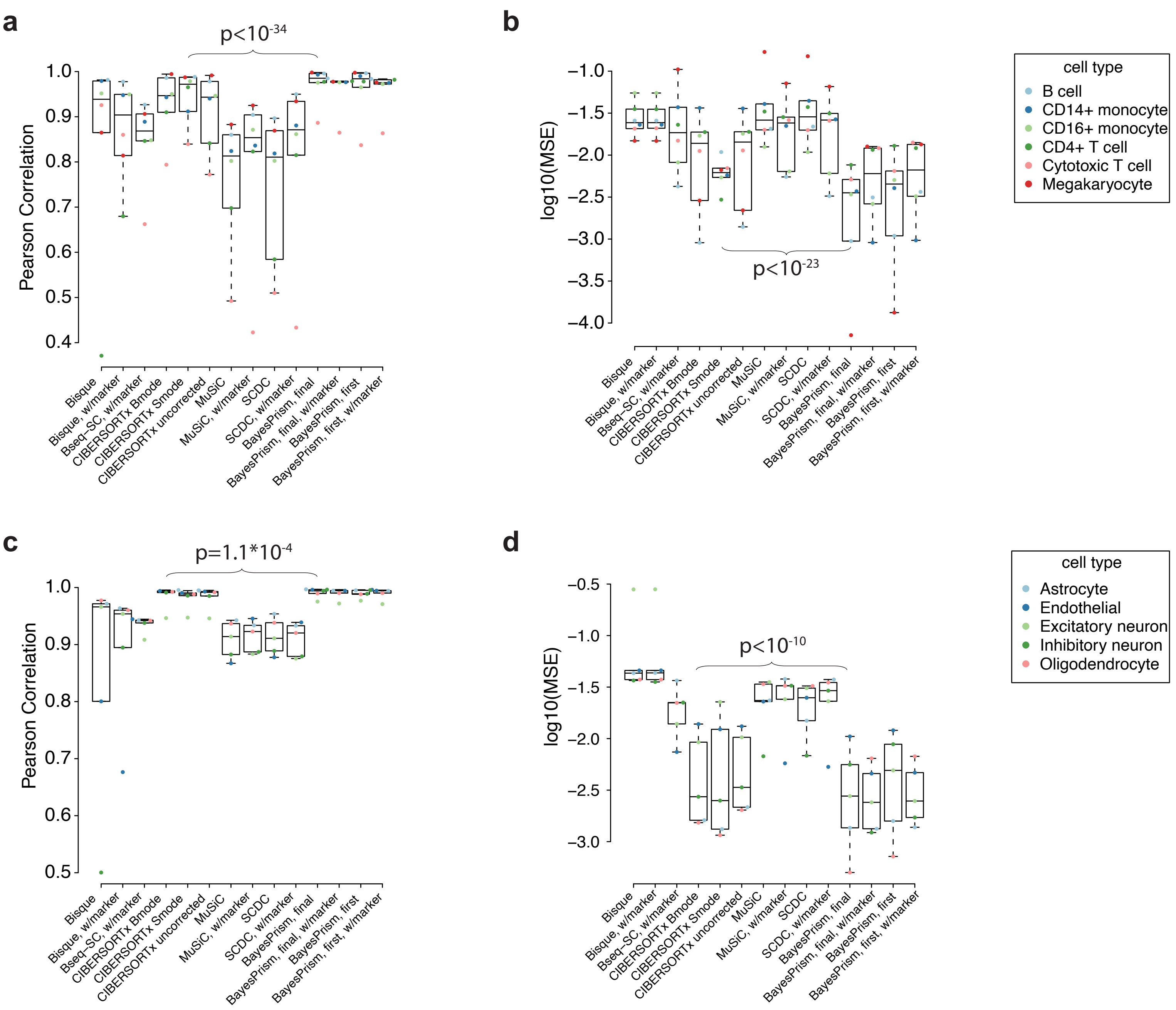


**Fig. S3 | Comparison between BayesPrism and other deconvolution methods on pseudo-bulks across different sequencing platforms and biological samples.** Boxplots show the cell type-level Pearson's correlation coefficient and MSE for the deconvolutions of pseudo-bulk human PBMC scRNA-seq (**a** and **b**) and mouse cortex single nucleus-seq (**c** and **d**). Boxes mark the 25th percentile (bottom of box), median (central bar), and 75th percentile (top of box). Whiskers represent extreme values within 1.5 fold of the inter quartile range. One-sided p values were shown for cell type fractions inferred by BayesPrism (updated θ using the marker free mode) and those by the second best methods ranked by the median value. T test was used for MSE and z test was performed on Fisher's Z-transformed cell type-level correlation coefficients (see Methods).

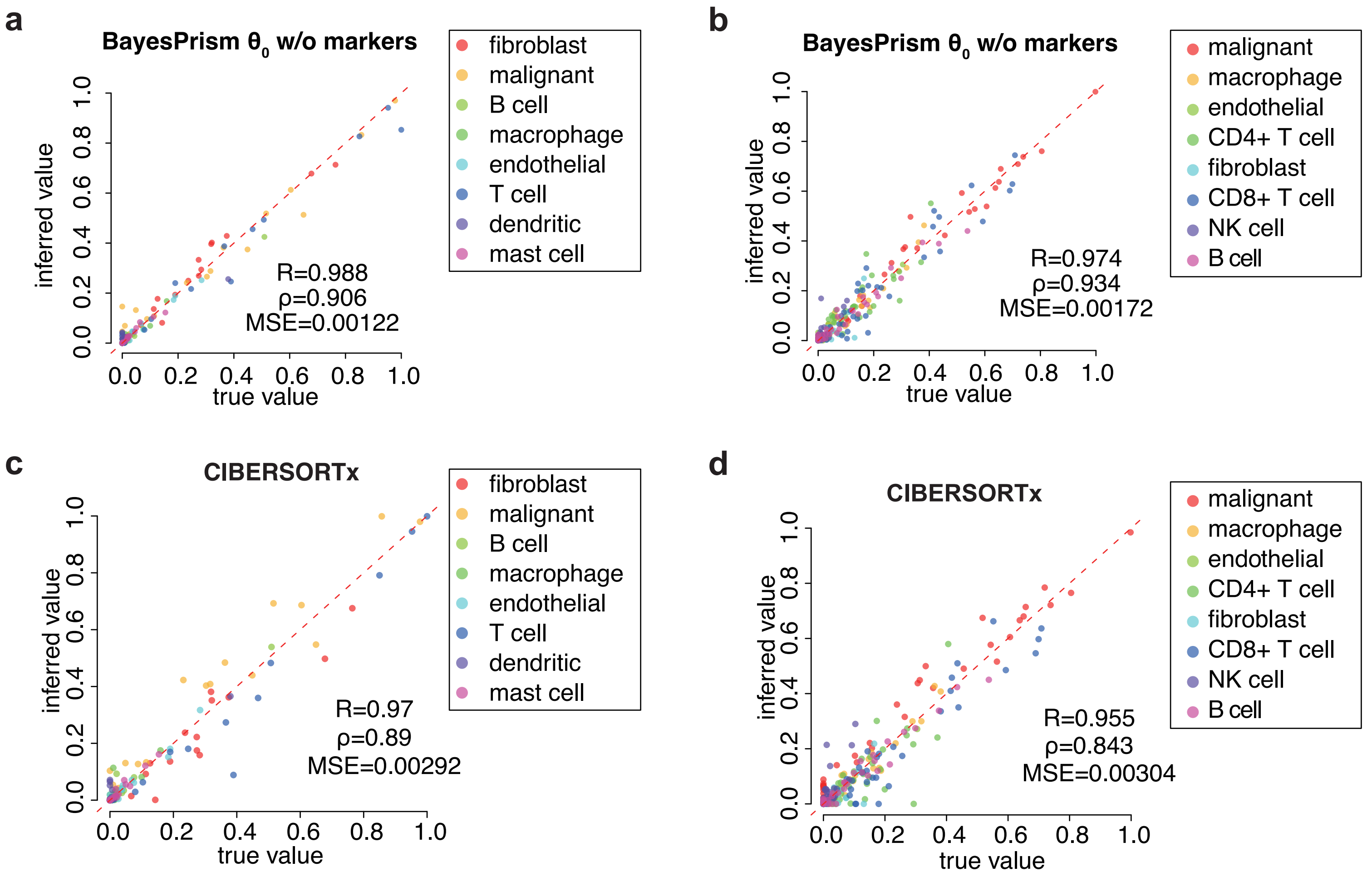


| MSE | Per cell type correlation | Total correlation |
|---|---|---|
| p=0.011 | p=0.0018 | p=3.7*10-5 |

| MSE | Per cell type correlation | Total correlation |
|---|---|---|
| p=0.00032 | p=0.63 | p=0.00076 |

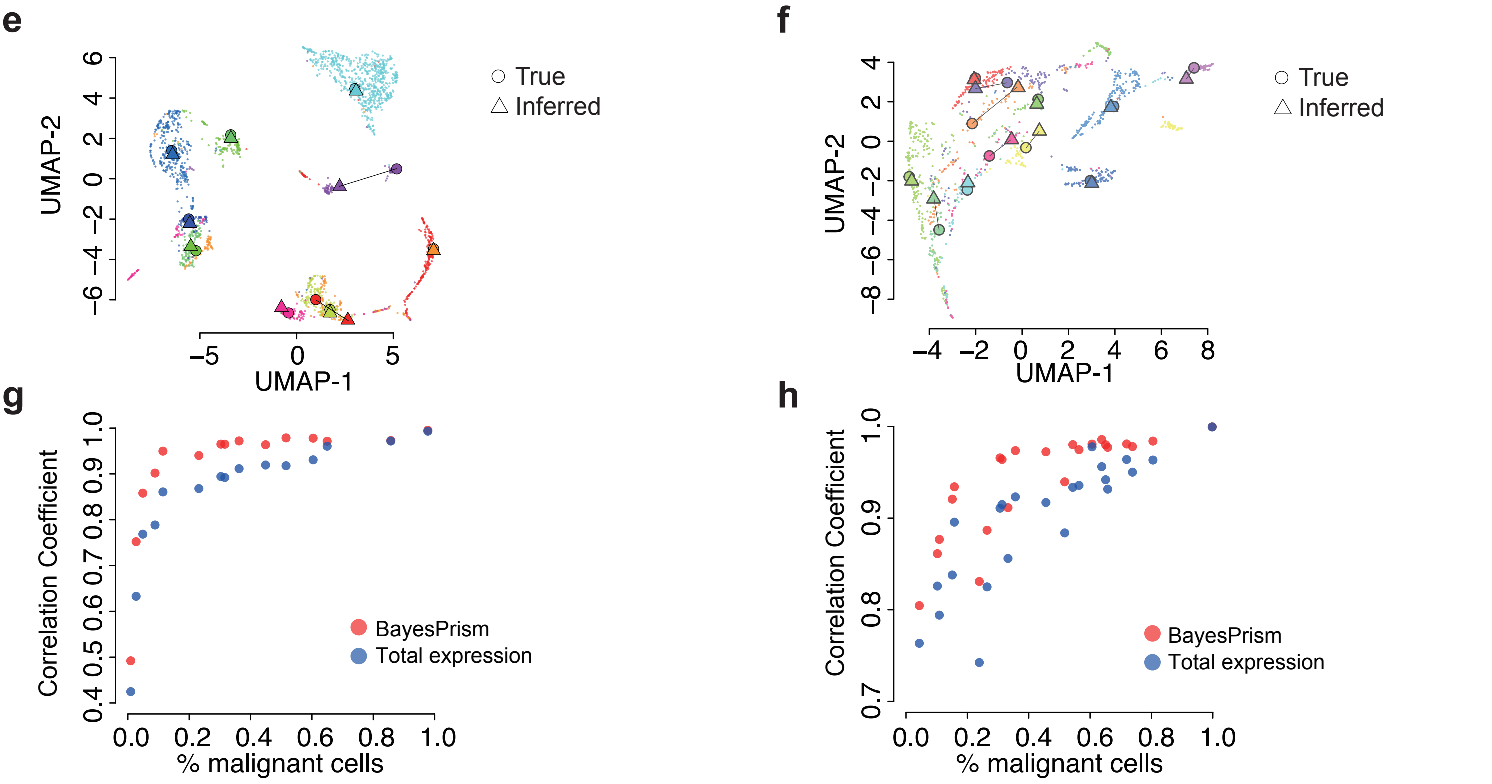


**Fig. S4 | Performance of BayesPrism in inferring cell type composition and gene expression in malignant cells on the leave-one-out pseudo-bulk data of HNSCC and SKCM. a-b**) Scatter plots show $\theta_0$ in **a**) HNSCC and **b**) SKCM versus the ground truth in pseudo-bulk. **c-d**) Scatter plots show the CIBERSORTx inferred cell type fraction in **c**) HNSCC and **d**) SKCM versus the ground truth in pseudo-bulk. Tables below show the one-sided p values for cell type fractions inferred by BayesPrism and CIBERSORTx. T test was used for MSE and z test was performed on Fisher's Z-transformed cell type-level correlation coefficients (see Methods). **e-f**) UMAP shows the expression profile of individual malignant cells in scRNA-seq of **e**) HNSCC and **f**) SKCM colored by patient ID. Patients with >50 malignant cells and cells with reads detected for >3000 genes are shown. The inferred expression profile, shown as △, and the averaged expression profile from scRNA-seq for each patient, shown as ◯, are projected onto the UMAP manifold. **g-h**) Scatter plot shows Pearson's correlation coefficient between inferred expression and that of the averaged expression from malignant cells in scRNA-seq of **g**) HNSCC and **h**) SKCM as a function of the fraction of malignant cells in each simulated data. The correlation coefficient was computed on DESeq2 variance-stabilized transformed values. Red marks the correlation inferred by BayesPrism, while blue marks that of total expression of the simulated data.

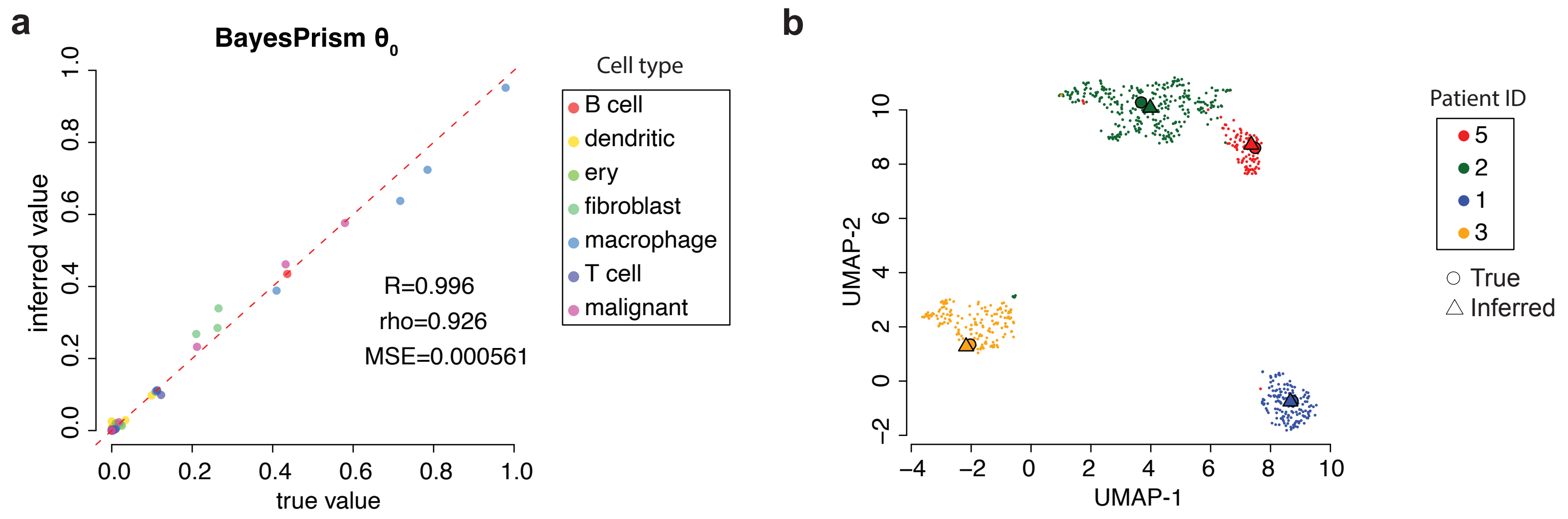


**Fig. S5 | Performance of BayesPrism in inferring cell type composition and gene expression in malignant cells on the leave-one-out pseudo-bulk data of ovarian cancer. a**) Scatter plots show the $\theta_0$ versus the ground truth in pseudo-bulk. **b**) UMAP shows the expression profile of individual malignant cells in scRNA-seq colored by patient ID. The inferred expression profile, shown as △, and the averaged expression profile from scRNA-seq for each patient, shown as ◯, are projected onto the UMAP manifold.

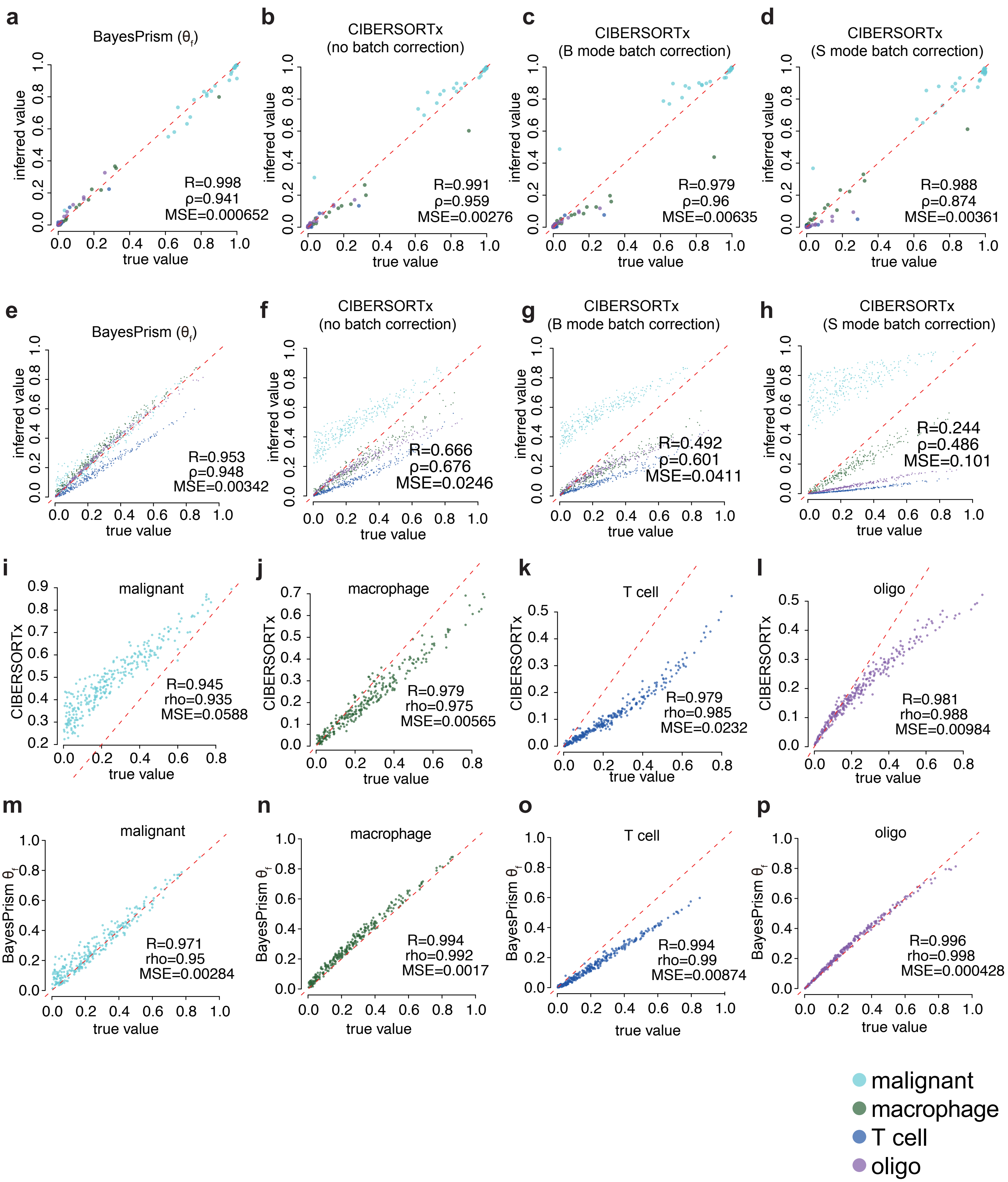


**Fig. S6 | Comparison between BayesPrism and various modes of CIBERSORTx.** Scatter plots show the inferred cell type fraction in the pseudo-bulk GBM28 (**a-d**), and a pseudo-bulk dataset containing 270 simulated samples (**e-h**). (**i-p**) Scatter plots show the performance of individual cell types in **e** and **f**.

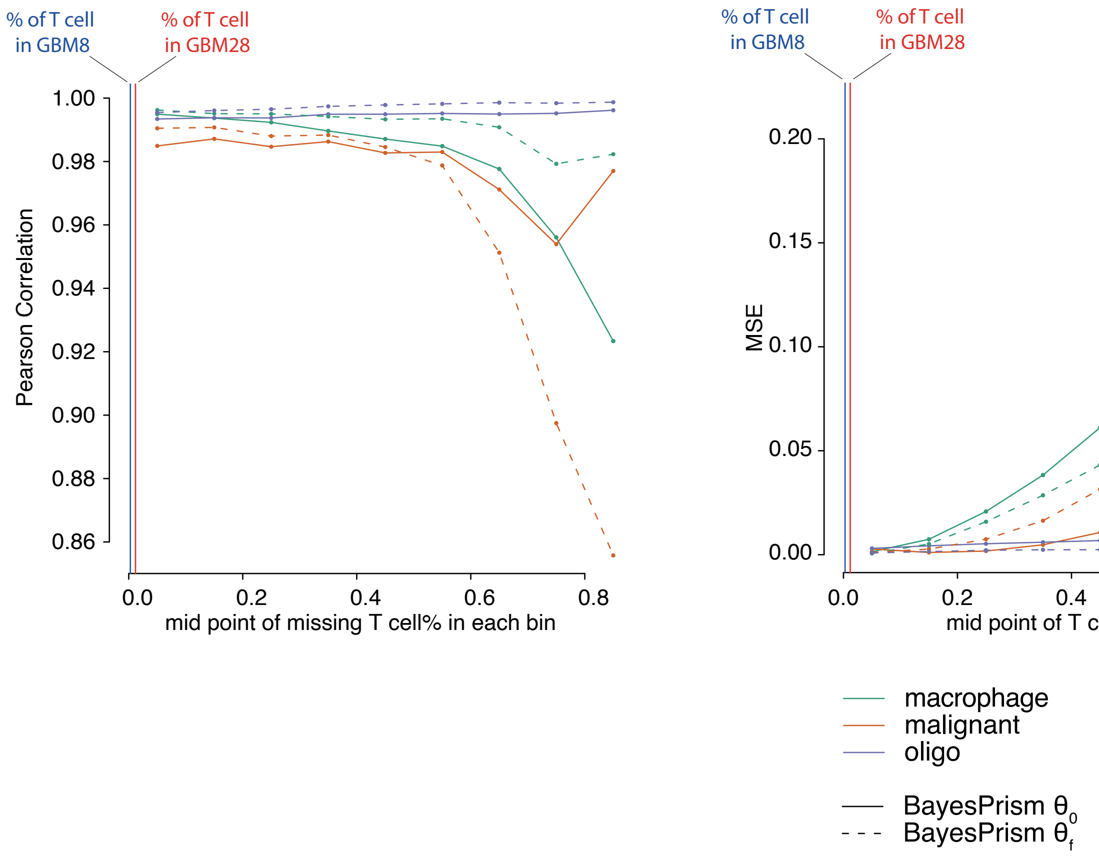


**Fig. S7 | BayesPrism is robust to missing cell types in the reference.** Line plots show the cell type-level Pearson's correlation coefficient (left) and MSE (right) for the deconvolution of simulated GBM28 (N=1350) using refGBM8 with T cells removed as the reference. The X axis marks the midpoints of each 10% width bin. Lines are colored by cell type. Solid lines represent $\theta_0$, while dashed lines represent $\theta_f$. Vertical bars mark the average observed T cell fraction in GBM8 and GBM28.

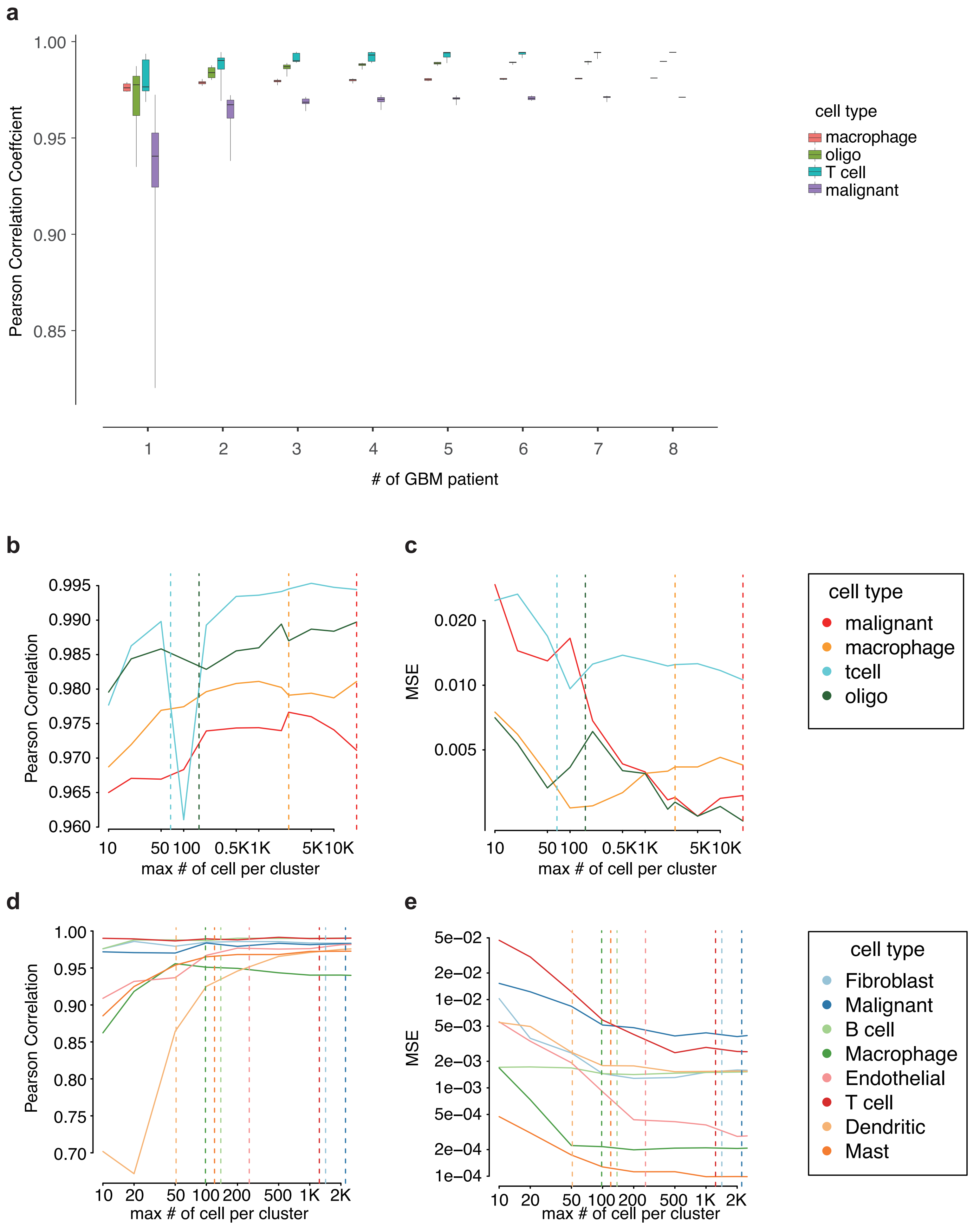


**Fig. S8 | BayesPrism is robust to a downsampled reference scRNA-seq dataset. a**) Boxplots show the distribution of cell type-level Pearson's correlation coefficient as a function of the number of downsampled patients in refGBM8, in which malignant cells are excluded from patients that are not sampled. Boxes mark the 25th percentile (bottom of box), median (central bar), and 75th percentile (top of box). Whiskers represent extreme values within 1.5 fold of the inter quartile range. **b**-**e**) Line plots show the cell type-level Pearson's correlation coefficient (left) and MSE (right) for the deconvolution of simulated GBM28 (N=1350) by refGBM8 (**b** and **c**) and HNSCC leave-one-out test using reference with downsampled single cells (**d** and **e**). The X axis marks the maximum number of cells in each cell type (or cell states of malignant cells) in the reference. Lines are colored by cell type. Dashed vertical lines mark the observed number of cells in each cell type in the original reference.

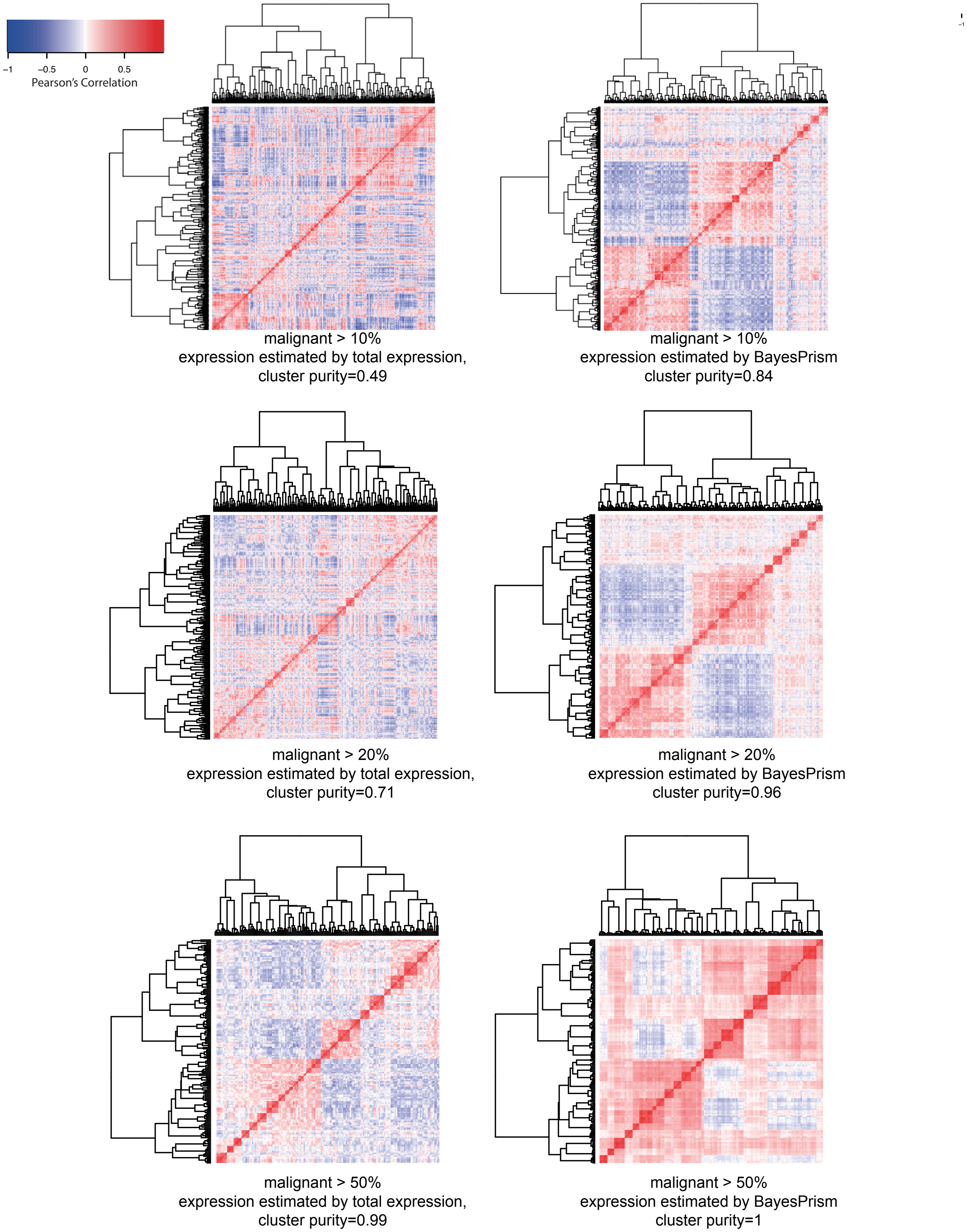


**Fig. S9 | Clustering BayesPrism expression estimates groups malignant cells from the same patient.** Heatmaps show the pairwise Pearson correlation matrix between gene expression computed for each pair of simulated pseudo-bulk samples with the fraction of malignant cells greater than 10% (**top row**), 20% (**mid row**) and 50% (**bottom row**). Simulated samples were constructed by drawing a random proportion of each non-malignant cell type, and sampling the malignant cells from one of 27 GBM patients. Vectors used for computing the Pearson correlation are of length equal to the total number of genes used to perform deconvolution, denoting zero centered variance-stabilizing transformed reads. Simulated pseudo-bulk samples are grouped by hierarchical clustering, as shown by the dendrogram. **Left column**: Correlations using total expression from bulk samples without any correction; **Right column**: Correlations over the same set of samples and genes as in the left column, but using deconvolved expression profiles for malignant cells in each sample.

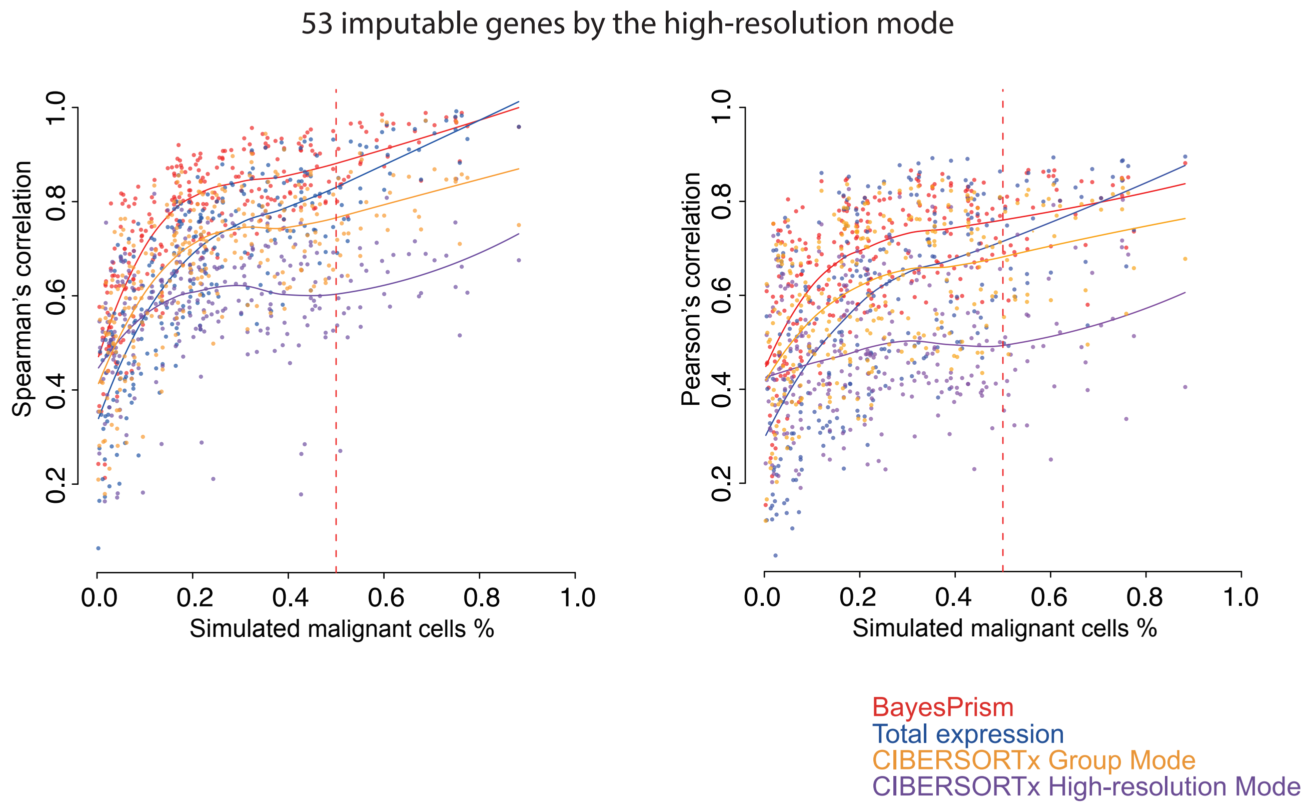


**Fig. S10 | Comparison between BayesPrism and the two different modes of expression inference by CIBERSORTx.** Scatter plot shows Spearman's correlation (left), and Pearson's correlation (right), between gene expression estimated by BayesPrism (red), total bulk (blue), CIBERSORTx group mode (orange) or CIBERSORTx high resolution mode (purple) and the average expression from malignant cells in scRNA-seq as a function of the fraction of malignant cells in the dataset containing the 270 simulated samples. The correlation coefficient was calculated on 53 imputable genes by the high-resolution mode out of the top 1000 most variable genes in malignant cells. Spearman's correlation coefficients were calculated on untransformed gene expression values, while Pearson's correlation coefficients were calculated on variance-stabilizing transformed gene expression values.

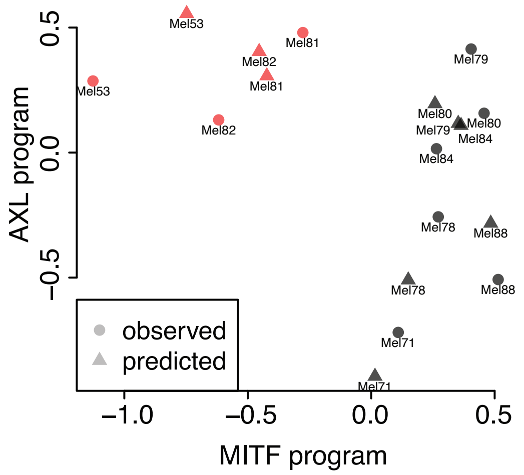


**Fig. S11 | BayesPrism accurately recovers the AXL/MITF program of malignant cells from SKCM in the leave-one-out test.** Scatter plot shows the AXL and MITF program scores, calculated by the mean of Z scores over the corresponding marker genes. △ marks the average expression of malignant cells in each patient from the scRNA-seq data, while ◯ marks the inferred expression profile of malignant cells in each patient.

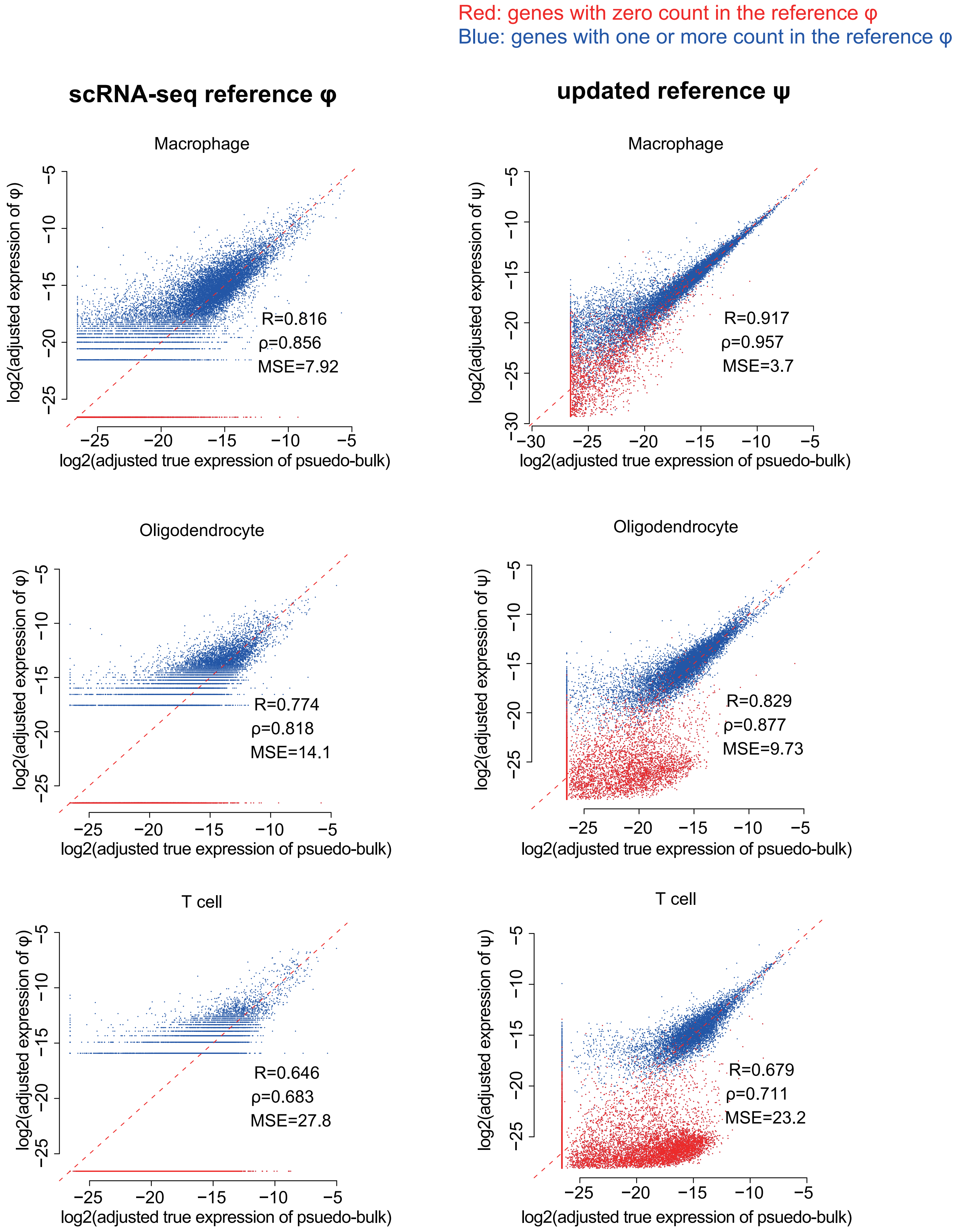


**Fig. S12 | BayesPrism improves correlations between gene expression in reference and ground truth pseudo-bulk data in non-malignant cells.** Scatter plots show $\log_2$ gene expression in non-malignant cells from the original scRNA-seq reference $\varphi$ (the left column) and the updated reference $\psi$ after the information pooling step (the right column) versus the mean expression in scRNA-seq used to generate pseudo-bulk. Genes with zero expression counts in the scRNA-seq reference are colored in red, and those with non-zero expression counts are colored in blue.

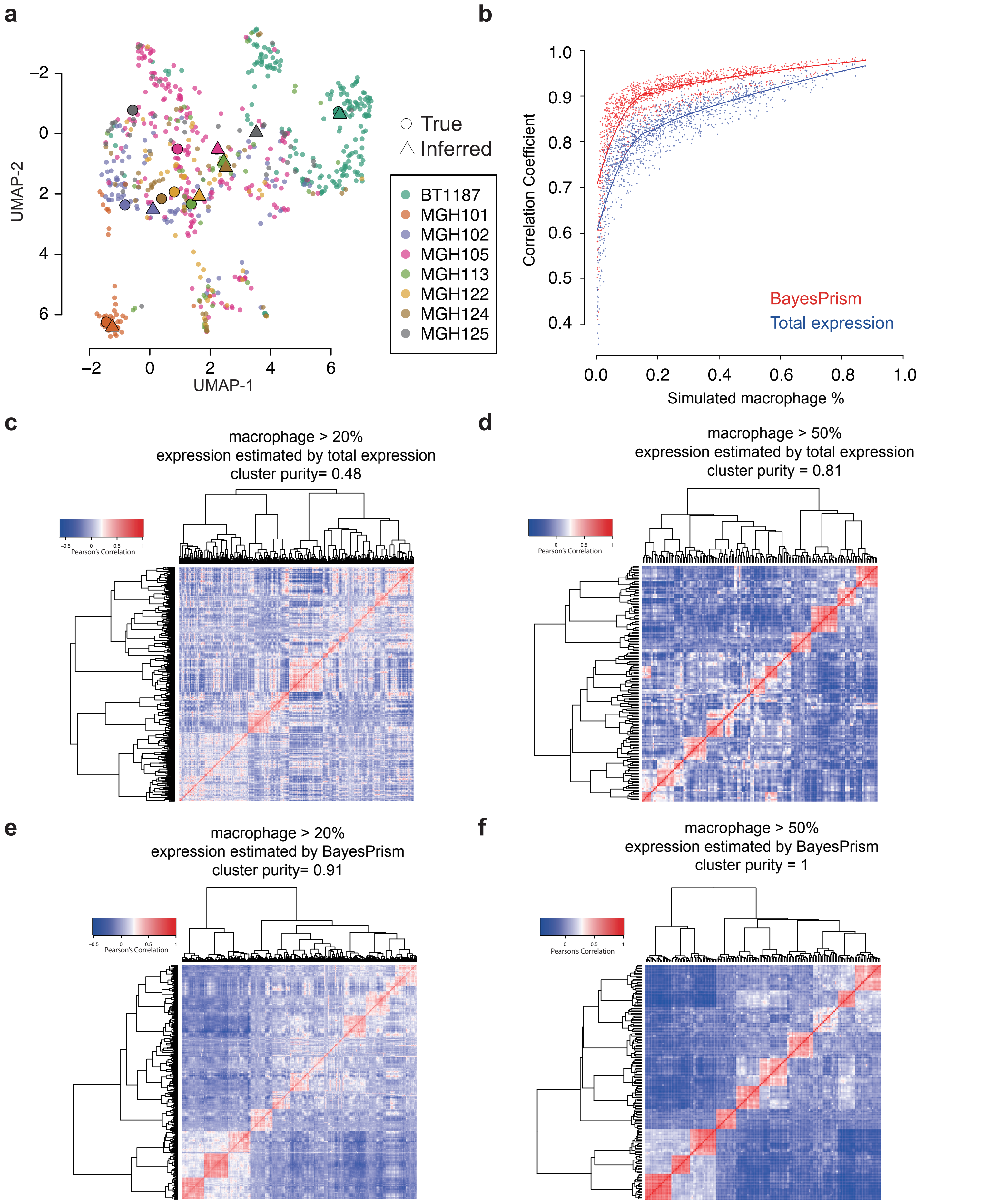


**Fig. S13** | **BayesPrism accurately recovers the heterogeneity in the expression of macrophages. a**) UMAP visualization shows the expression of individual macrophages in the pseudo-bulk GBM28 dataset. The expression profile inferred by BayesPrism, shown as △, and the averaged expression profile from scRNA-seq for each patient, shown as ◯, are projected onto the UMAP manifold. **b**) Scatter plot shows Pearson's correlation of reads summed across macrophages in the pseudo-bulk (ground truth) and gene expression deconvolved by BayesPrism (red) or undeconvovled pseudo-bulk (blue) as a function of the fraction of macrophages in the simulated pseudo-bulk (N=1,350). The correlation was computed using variance-stabilizing transformed reads. **c**-**f**) Heatmap shows the pairwise Pearson correlation matrix between gene expression computed for each pair of simulated pseudo-bulk samples with macrophage fractions greater than 20% (**c**, **e**) and 50% (**d**, **f**). Simulated samples were obtained by drawing a random proportion of each cell type from the GBM-28 dataset, while sampling macrophages from an individual macrophage sub-cluster. Vectors used to compute Pearson's correlation are of length equal to the total number of genes used to perform deconvolution, and represent zero centered variance-stabilizing transformed read counts for each gene. Simulated pseudo-bulk samples are grouped by hierarchical clustering, as shown by the dendrogram. **c**, **d**) Correlations estimated using the total gene expression without any correction. **e**, **f**) Correlations over the same set of samples and over the same set of genes as in **c** and **d**, but using BayesPrism deconvolved expression profiles for macrophages in each sample.

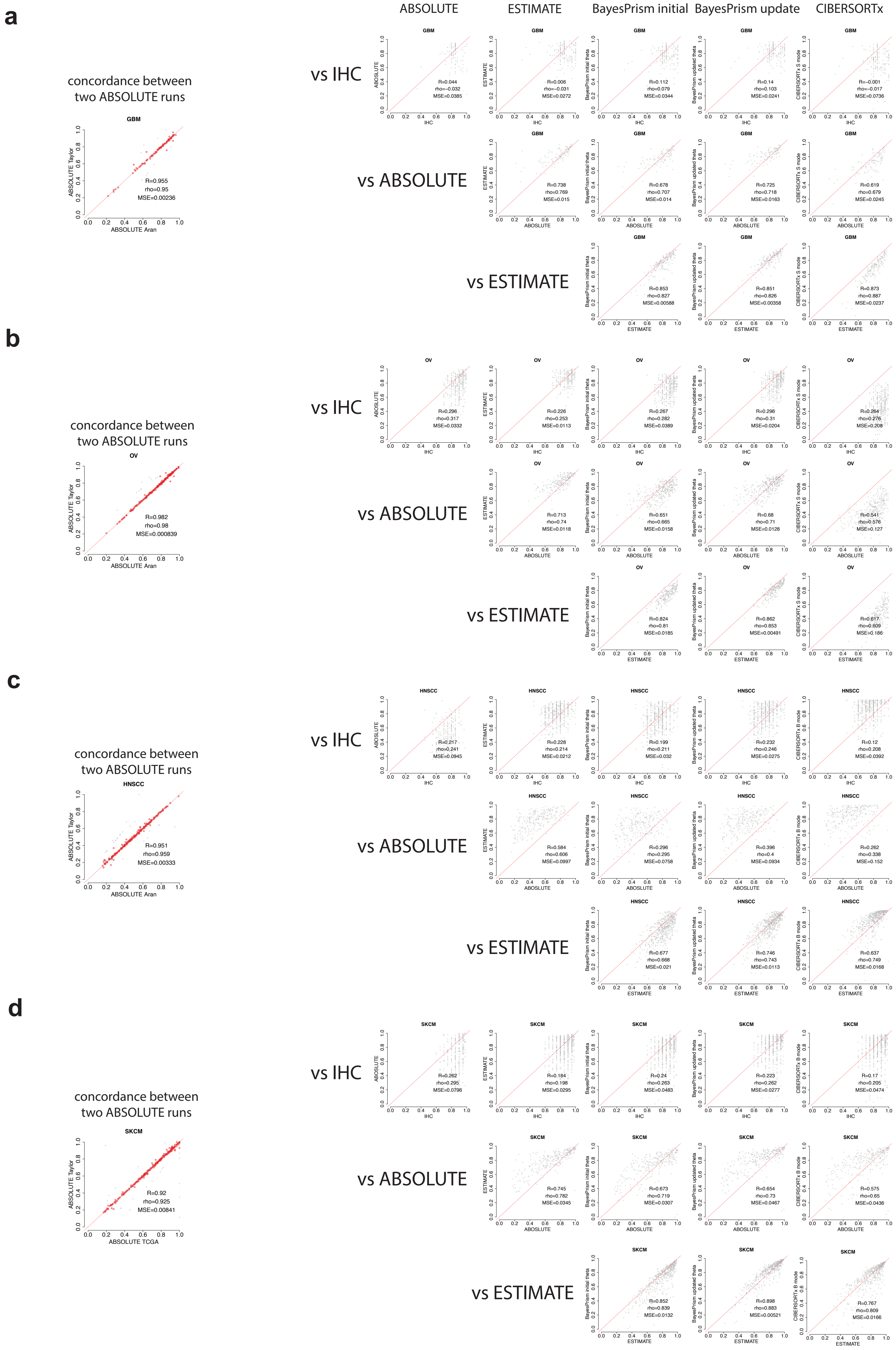

**Fig. S14 | Comparison between tumor purity inferred by BayesPrism, CIBERSORTx, ABSOLUTE, ESTIMATE and IHC.** Scatter plots show the correlation between malignant fractions inferred by each method for TCGA-GBM (**a**), TCGA-OV (**b**), TCGA-HNSCC (**c**), and TCGA-SKCM (**d**). Dashed lines mark the y=x. As the tumor purity estimated by ABSOLUTE varies between multiple sources (Aran et. al. (2015) *Nature Communications*; Taylor et. al. (2018) *Cancer Cell*; TCGAN (2015) *Cell*) (**leftmost columns**), we took the mean of ABSOLUTE scores between two sources over a subset of samples whose difference between two sources is smaller than 0.1 (marked by the red dots), and used the averaged score for comparison with other methods (**right panels**). ESTIMATE and IHC scores were both obtained from Aran et al. CIERSORTx was run using the same input reference matrix as BayesPrism, using the batch correction mode recommended by the authors (S mode for GBM and OV; B mode for SKCM and HNSCC). In HNSCC we observed that ABSOLUTE score was lower than IHC, while tumor purity estimated using transcription signal (ESTIMATE, BayesPrism and CIBERSORTx) were higher than ABSOLUTE, suggesting an underestimation of tumor purity by the DNA copy-number based method, and/or the presence of non-immune non-stromal cells (such as normal epithelial cells, which was not captured by scRNA-seq reference) in the tumor samples.

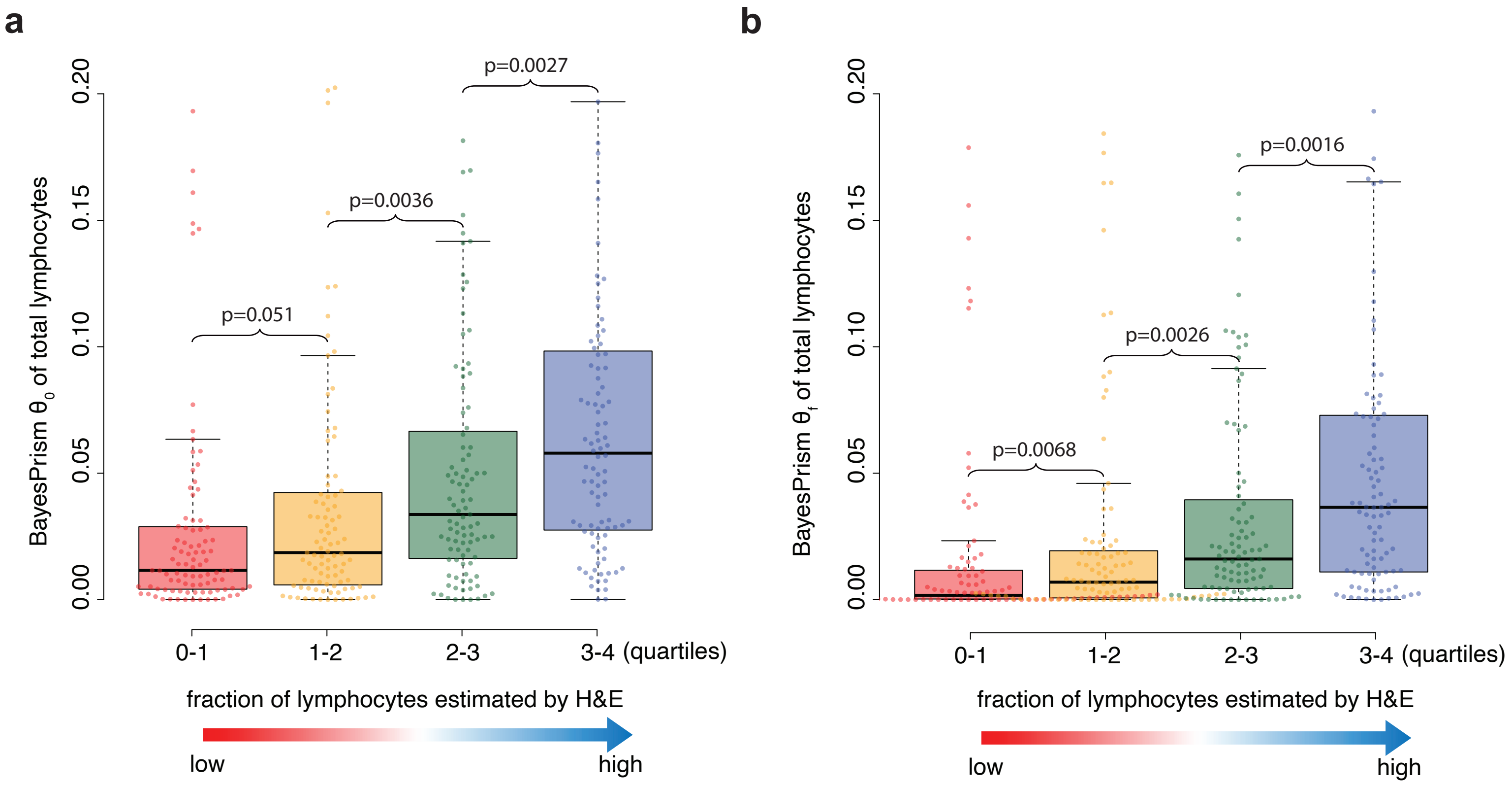


**Fig. S15 | Comparison between the total lymphocyte fraction estimated by BayesPrism and H&E.** Box plots show the distribution of $\theta_0$ (**a**) and $\theta_f$ (**b**) of total lymphocytes computed by summing across CD4+ and CD8+ T cells B cells and NK cells. Box plots are binned by the quantiles of the fraction of H&E patches classified positive for tumor infiltrating lymphocytes (TIL). P values were calculated using the one-sided Wilcox test.

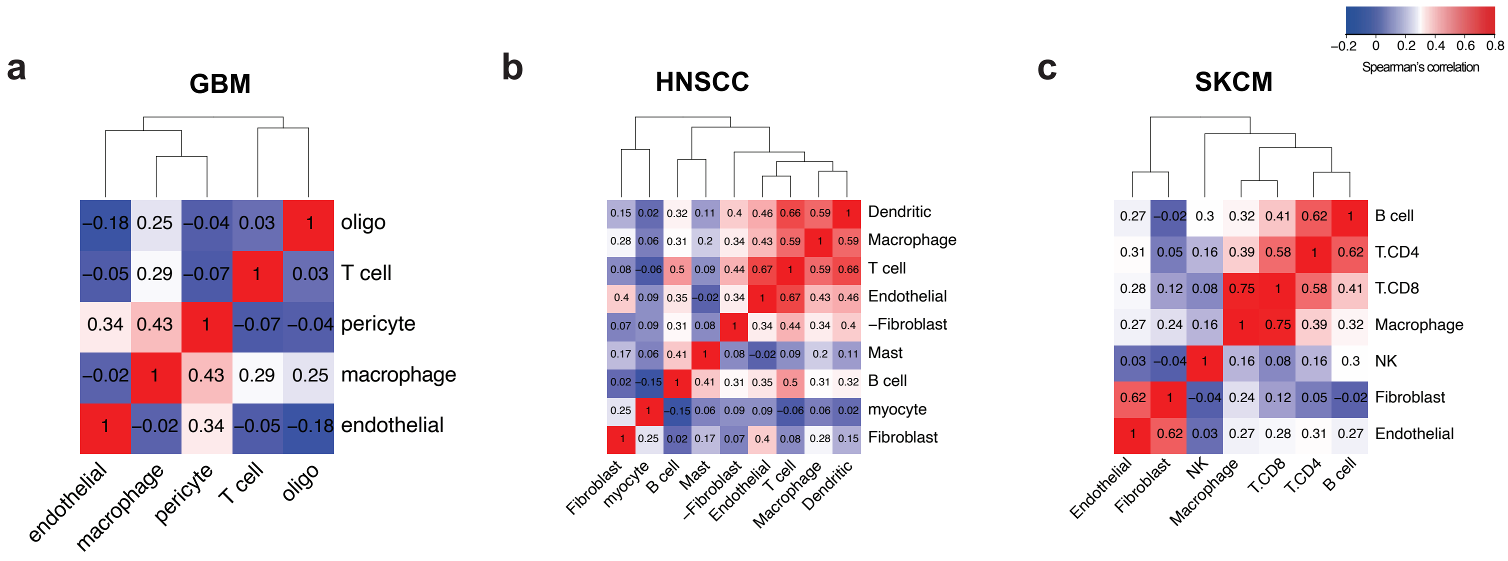


**Fig. S16 | Heatmaps show the Spearman's rank correlation between non-malignant cells in each tumor type.**

**a** **GBM**

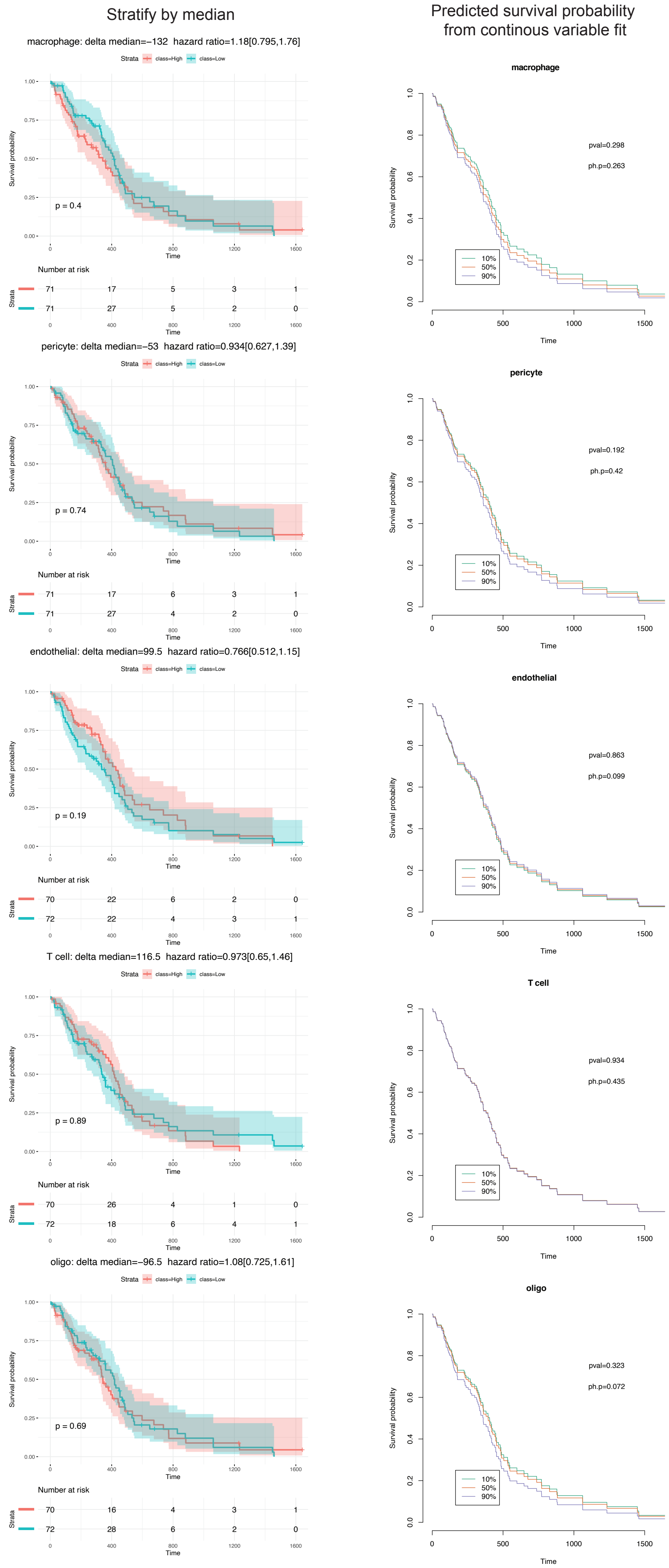

Stratify by median
Predicted survival probability from continous variable fit
macrophage: delta median=−132 hazard ratio=1.18[0.795,1.76]
p = 0.4
Number at risk
71 17 5 3 1
71 27 5 2 0
macrophage
pval=0.298
ph.p=0.263
pericyte: delta median=−53 hazard ratio=0.934[0.627,1.39]
p = 0.74
Number at risk
71 17 6 3 1
71 27 4 2 0
pericyte
pval=0.192
ph.p=0.42
endothelial: delta median=99.5 hazard ratio=0.766[0.512,1.15]
p = 0.19
Number at risk
70 22 6 2 0
72 22 4 3 1
endothelial
pval=0.863
ph.p=0.099
T cell: delta median=116.5 hazard ratio=0.973[0.65,1.46]
p = 0.89
Number at risk
70 26 4 1 0
72 18 6 4 1
T cell
pval=0.934
ph.p=0.435
oligo: delta median=−96.5 hazard ratio=1.08[0.725,1.61]
p = 0.69
Number at risk
70 16 4 3 1
72 28 6 2 0
oligo
pval=0.323
ph.p=0.072
Strata class=High class=Low
Survival probability
Time
10%
50%
90%

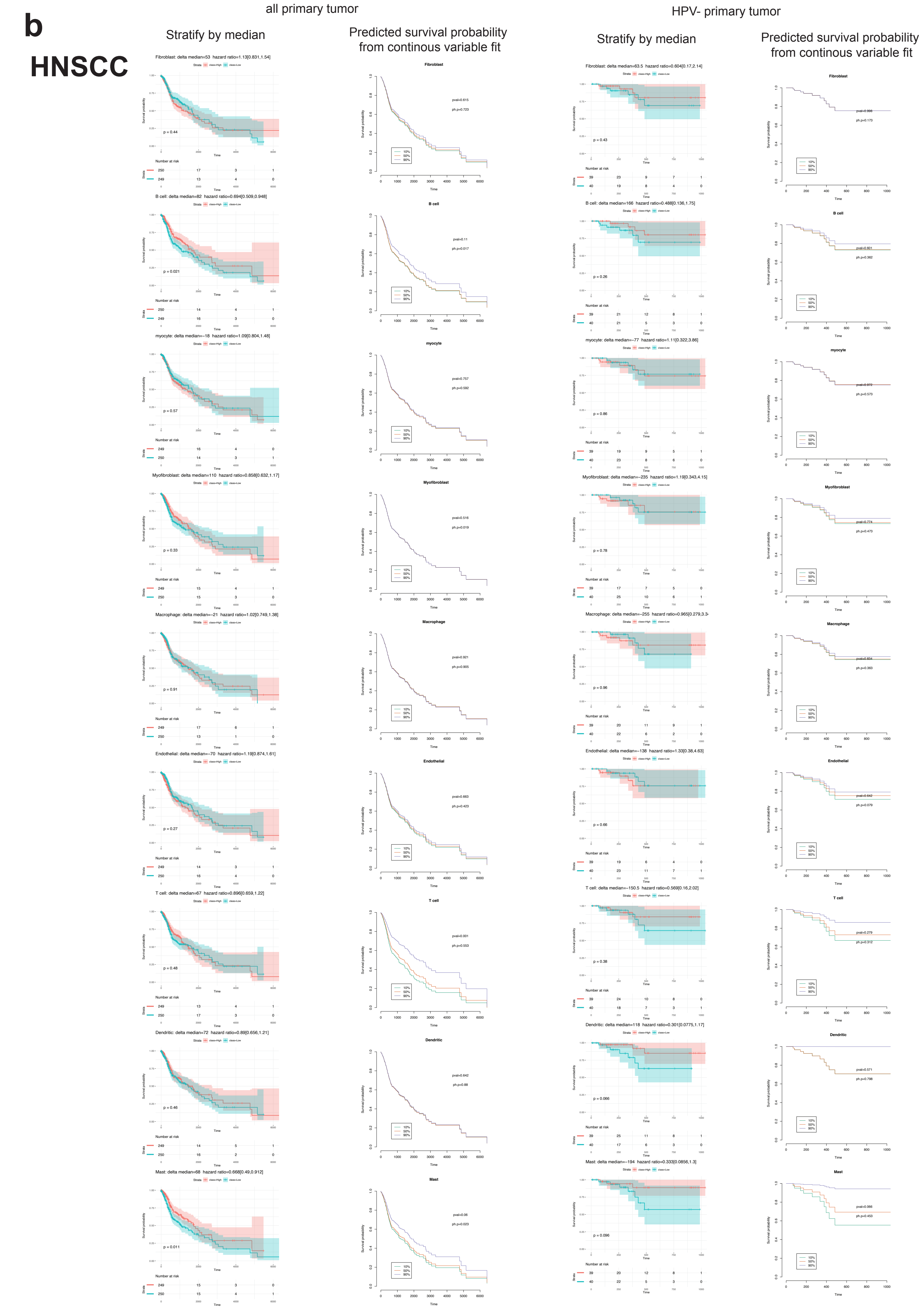

b
HNSCC
all primary tumor
HPV- primary tumor
Stratify by median
Predicted survival probability from continous variable fit
Stratify by median
Predicted survival probability from continous variable fit
Fibroblast: delta median=53 hazard ratio=1.13[0.831,1.54]
B cell: delta median=82 hazard ratio=0.694[0.509,0.948]
myocyte: delta median=-18 hazard ratio=1.09[0.804,1.48]
Myofibroblast: delta median=110 hazard ratio=0.858[0.632,1.17]
Macrophage: delta median=-21 hazard ratio=1.02[0.749,1.38]
Endothelial: delta median=-70 hazard ratio=1.19[0.874,1.61]
T cell: delta median=67 hazard ratio=0.896[0.659,1.22]
Dendritic: delta median=72 hazard ratio=0.89[0.656,1.21]
Mast: delta median=68 hazard ratio=0.668[0.49,0.912]
Fibroblast: delta median=63.5 hazard ratio=0.604[0.17,2.14]
B cell: delta median=166 hazard ratio=0.488[0.136,1.75]
myocyte: delta median=-77 hazard ratio=1.11[0.322,3.86]
Myofibroblast: delta median=-235 hazard ratio=1.19[0.343,4.15]
Macrophage: delta median=-255 hazard ratio=0.965[0.279,3.3
Endothelial: delta median=-138 hazard ratio=1.33[0.38,4.63]
T cell: delta median=-150.5 hazard ratio=0.569[0.16,2.02]
Dendritic: delta median=118 hazard ratio=0.301[0.0775,1.17]
Mast: delta median=-194 hazard ratio=0.333[0.0856,1.3]

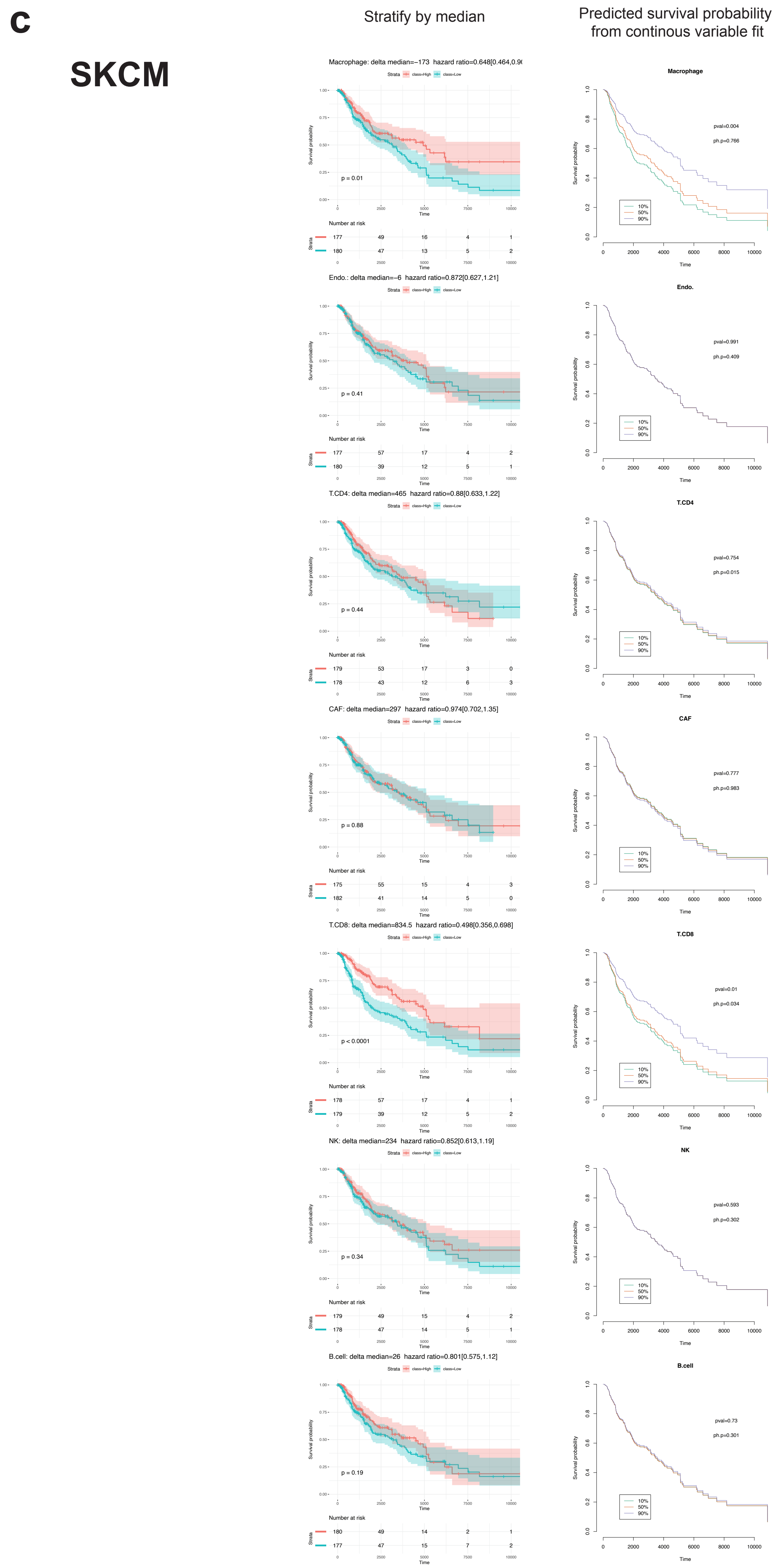


**Fig. S17 | KM plots of all cell types inferred from three tumor types across TCGA samples using median cutoff (the left column), and Cox proportional-hazards models in which the cell type fractions were modeled as a continuous variable (the right column).** The survival curves in the right column were plotted by conditioning on the cell type fraction at its 10%, 50% and 90% percentile, and then predicting survival probabilities based on the continuous variable Cox regression model, which was only used for the purpose of visualization. The log-rank test was used for the median-cutoff model. Two p values were computed for the continuous variable Cox regression model: "pval" was generated by the Wald test indicating the statistical significance of survival association, while "ph.p" was generated by the chi-squared test for scaled Schoenfeld residuals to check the proportional hazards assumption.

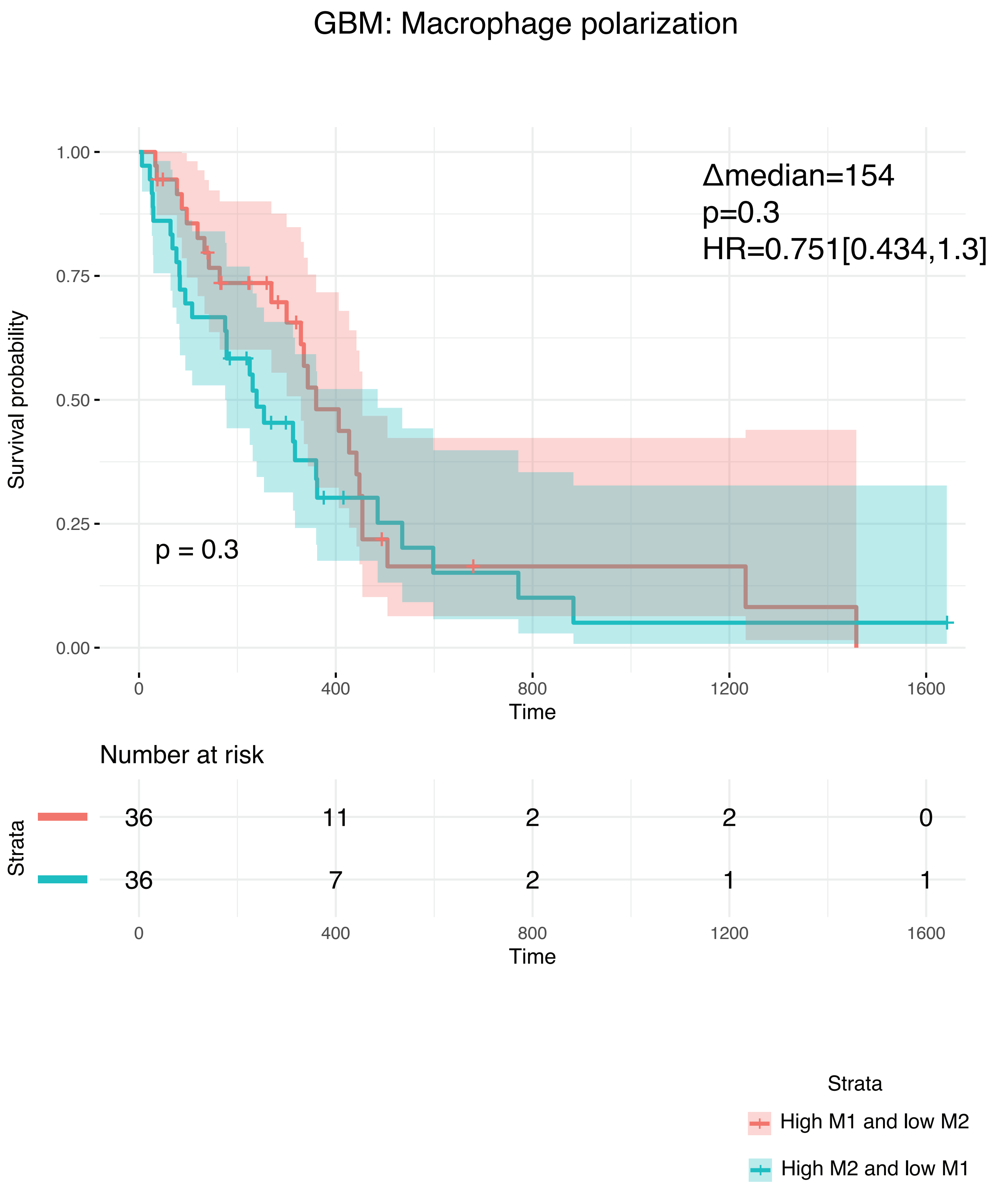


**Fig. S18 | KM plot shows the survival association with the M1/M2 state polarization of macrophages in GBM.**

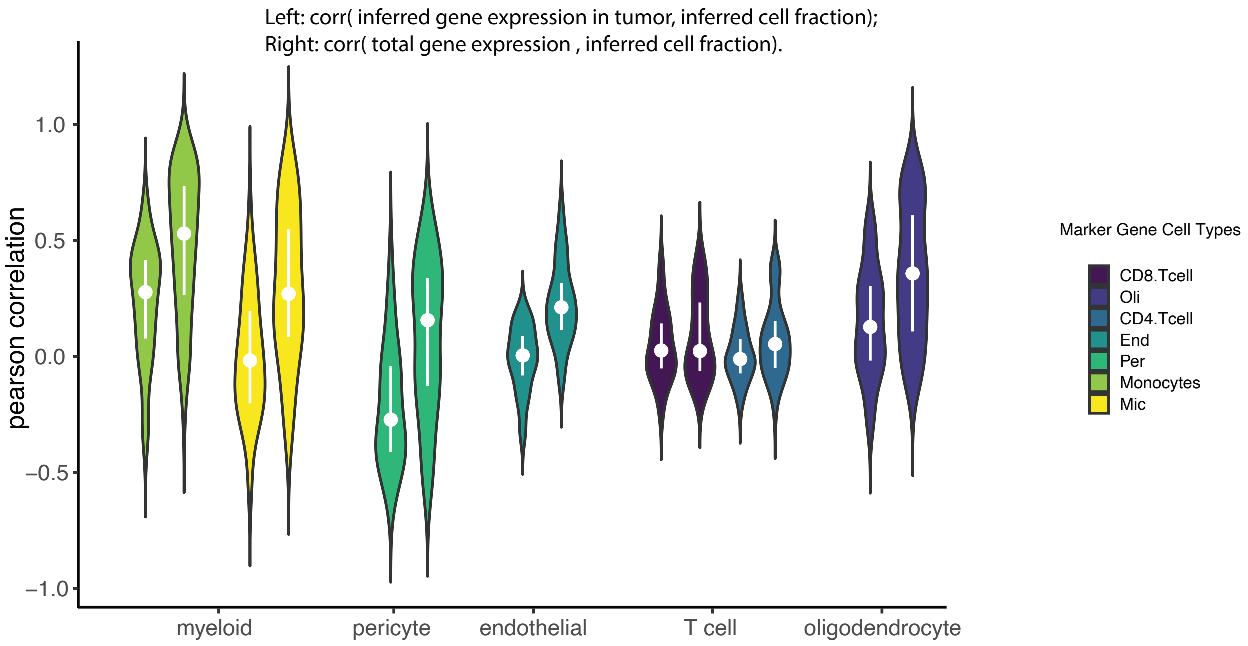


**Fig. S19 | BayesPrism removes false positive correlations from cell type marker genes.** Violin plot shows the distribution of Pearson's correlation between gene expression in malignant cells inferred by BayesPrism (left violins) or total gene expression of bulk RNA-seq (right violins) and BayesPrism predicted fractions of each cell type on their corresponding marker genes over TCGA-GBM. Median correlations are shown by white dots and upper/lower quartiles are shown by bars. Braces on the horizontal direction label the cell type fraction on which the correlations were computed. Color indicates the cell type of which the marker genes are curated from independent datasets.

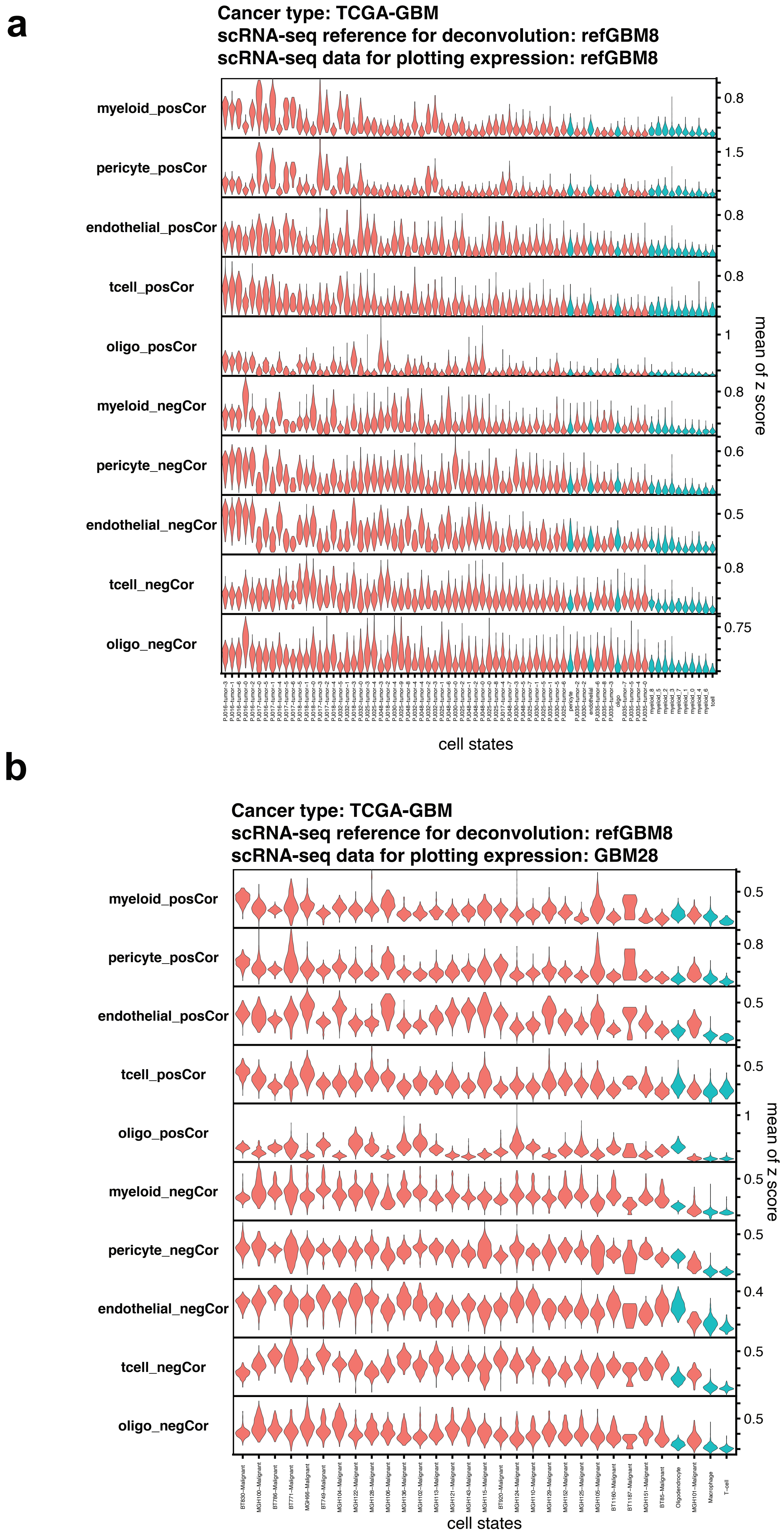
a
Cancer type: TCGA-GBM
scRNA-seq reference for deconvolution: refGBM8
scRNA-seq data for plotting expression: refGBM8
myeloid_posCor
pericyte_posCor
endothelial_posCor
tcell_posCor
oligo_posCor
myeloid_negCor
pericyte_negCor
endothelial_negCor
tcell_negCor
oligo_negCor
0.8
1.5
0.8
0.8
1
0.8
0.6
0.5
0.8
0.75
mean of z score
cell states
b
Cancer type: TCGA-GBM
scRNA-seq reference for deconvolution: refGBM8
scRNA-seq data for plotting expression: GBM28
myeloid_posCor
pericyte_posCor
endothelial_posCor
tcell_posCor
oligo_posCor
myeloid_negCor
pericyte_negCor
endothelial_negCor
tcell_negCor
oligo_negCor
0.5
0.8
0.5
0.5
1
0.5
0.5
0.4
0.5
0.5
mean of z score
cell states

**c**

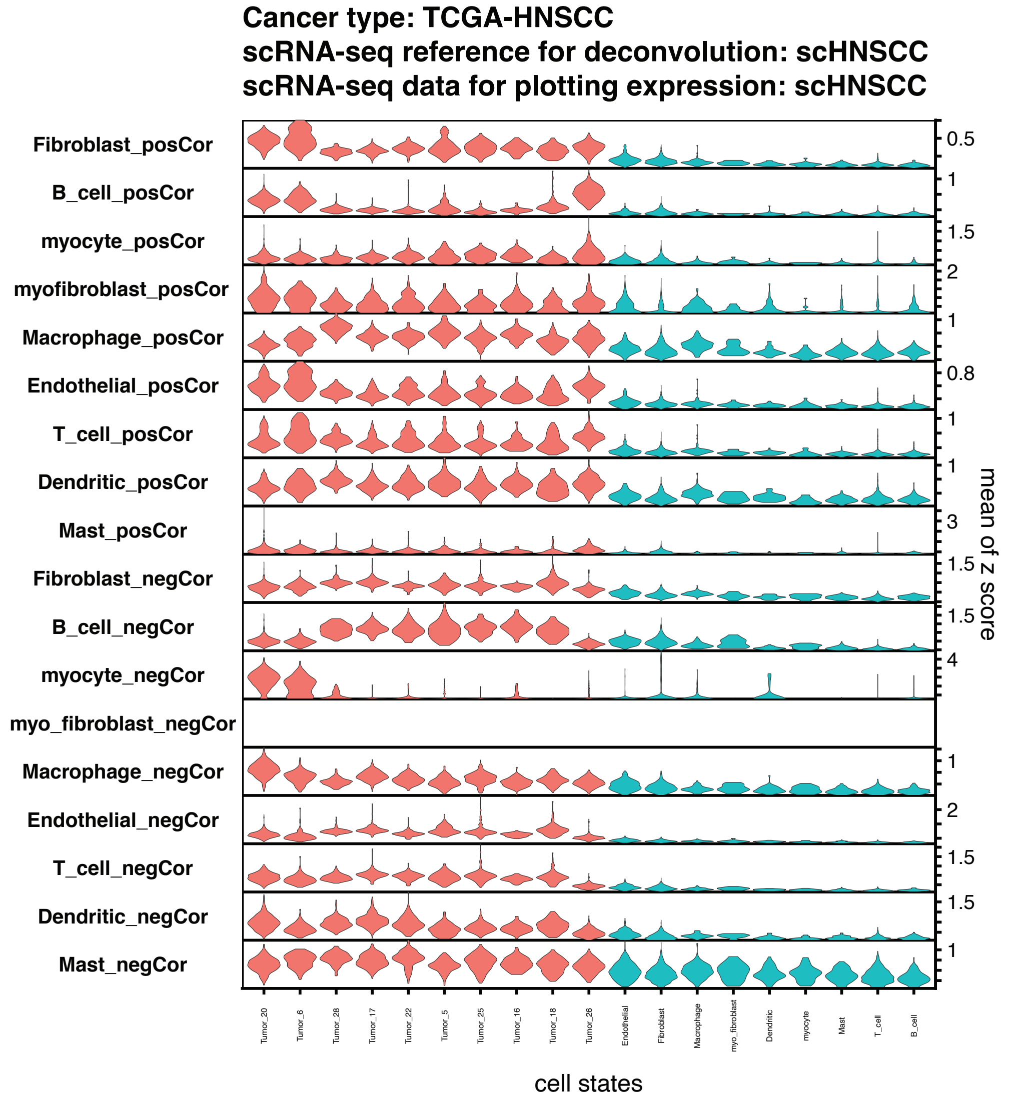


**d**

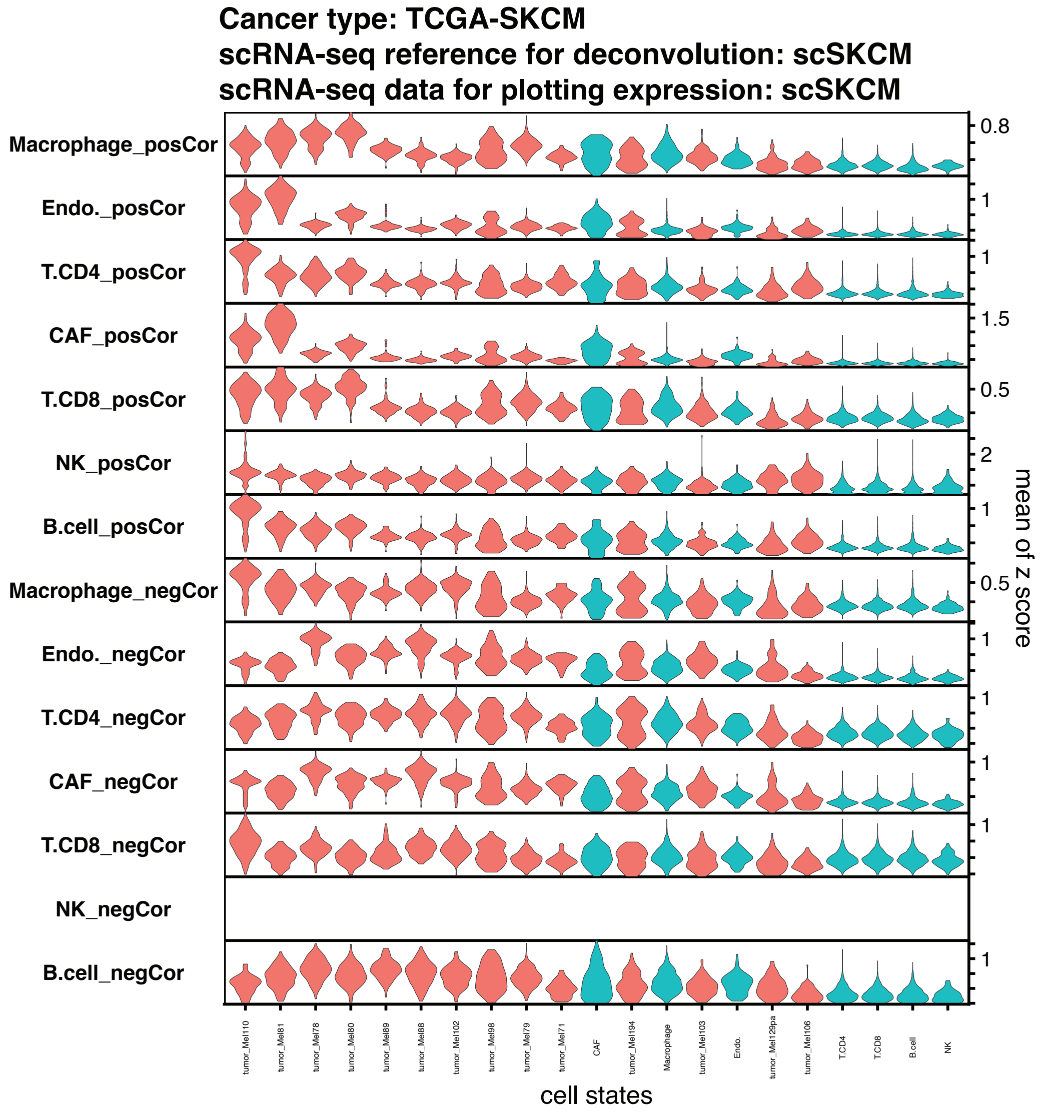


**Fig. S20 | Violin plots show the mean of z scores of expression over a gene set of interest (each row) computed using a particular scRNA-seq dataset.** Each row defines a set of genes that passed both the tumor intrinsic filter and the regress-out filter, and also correlated with the cell type fraction of a non-malignant cell type, with the absolute value of Spearman's correlation coefficient greater than 0.25. Each column denotes a cell state or cell type corresponding to the tumor intrinsic filter.

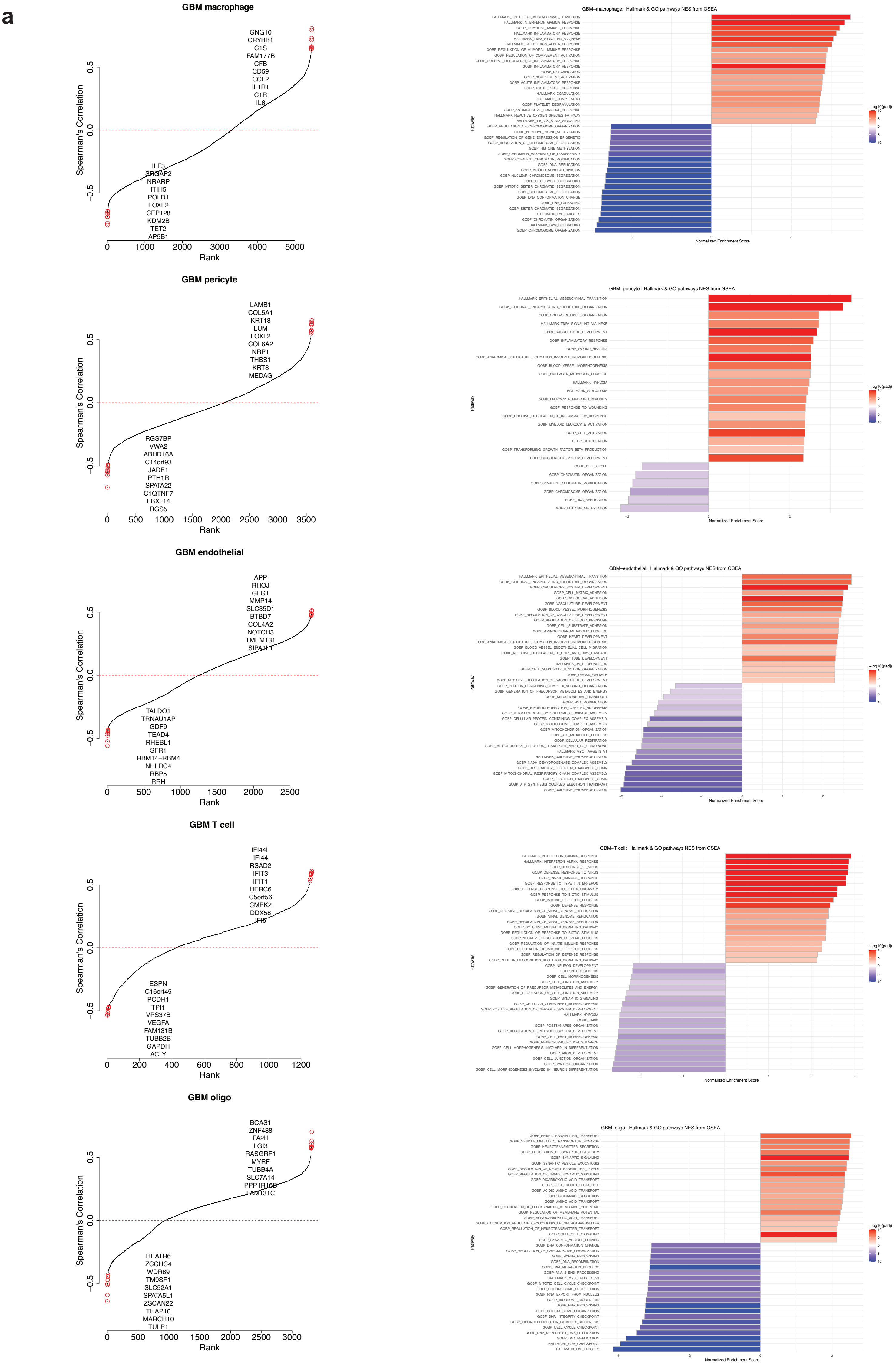

a
GBM macrophage
GNG10
CRYBB1
C1S
FAM177B
CFB
CD59
CCL2
IL1R1
C1R
IL6
ILF3
SRGAP2
NRARP
ITIH5
POLD1
FOXF2
CEP128
KDM2B
TET2
AP5B1
Spearman's Correlation
Rank
GBM−macrophage: Hallmark & GO pathways NES from GSEA
HALLMARK_EPITHELIAL_MESENCHYMAL_TRANSITION
HALLMARK_INTERFERON_GAMMA_RESPONSE
GOBP_HUMORAL_IMMUNE_RESPONSE
HALLMARK_INFLAMMATORY_RESPONSE
HALLMARK_TNFA_SIGNALING_VIA_NFKB
HALLMARK_INTERFERON_ALPHA_RESPONSE
GOBP_REGULATION_OF_HUMORAL_IMMUNE_RESPONSE
GOBP_REGULATION_OF_COMPLEMENT_ACTIVATION
GOBP_POSITIVE_REGULATION_OF_INFLAMMATORY_RESPONSE
GOBP_INFLAMMATORY_RESPONSE
GOBP_DETOXIFICATION
GOBP_COMPLEMENT_ACTIVATION
GOBP_ACUTE_INFLAMMATORY_RESPONSE
GOBP_ACUTE_PHASE_RESPONSE
HALLMARK_COAGULATION
HALLMARK_COMPLEMENT
GOBP_PLATELET_DEGRANULATION
GOBP_ANTIMICROBIAL_HUMORAL_RESPONSE
HALLMARK_REACTIVE_OXYGEN_SPECIES_PATHWAY
HALLMARK_IL6_JAK_STAT3_SIGNALING
GOBP_REGULATION_OF_CHROMOSOME_ORGANIZATION
GOBP_PEPTIDYL_LYSINE_METHYLATION
GOBP_REGULATION_OF_GENE_EXPRESSION_EPIGENETIC
GOBP_REGULATION_OF_CHROMOSOME_SEGREGATION
GOBP_HISTONE_METHYLATION
GOBP_CHROMATIN_ASSEMBLY_OR_DISASSEMBLY
GOBP_COVALENT_CHROMATIN_MODIFICATION
GOBP_DNA_REPLICATION
GOBP_MITOTIC_NUCLEAR_DIVISION
GOBP_NUCLEAR_CHROMOSOME_SEGREGATION
GOBP_CELL_CYCLE_CHECKPOINT
GOBP_MITOTIC_SISTER_CHROMATID_SEGREGATION
GOBP_CHROMOSOME_SEGREGATION
GOBP_DNA_CONFORMATION_CHANGE
GOBP_DNA_PACKAGING
GOBP_SISTER_CHROMATID_SEGREGATION
HALLMARK_E2F_TARGETS
GOBP_CHROMATIN_ORGANIZATION
HALLMARK_G2M_CHECKPOINT
GOBP_CHROMOSOME_ORGANIZATION
Pathway
Normalized Enrichment Score
−log10(padj)
GBM pericyte
LAMB1
COL5A1
KRT18
LUM
LOXL2
COL6A2
NRP1
THBS1
KRT8
MEDAG
RGS7BP
VWA2
ABHD16A
C14orf93
JADE1
PTH1R
SPATA22
C1QTNF7
FBXL14
RGS5
Spearman's Correlation
Rank
GBM−pericyte: Hallmark & GO pathways NES from GSEA
HALLMARK_EPITHELIAL_MESENCHYMAL_TRANSITION
GOBP_EXTERNAL_ENCAPSULATING_STRUCTURE_ORGANIZATION
GOBP_COLLAGEN_FIBRIL_ORGANIZATION
HALLMARK_TNFA_SIGNALING_VIA_NFKB
GOBP_VASCULATURE_DEVELOPMENT
GOBP_INFLAMMATORY_RESPONSE
GOBP_WOUND_HEALING
GOBP_ANATOMICAL_STRUCTURE_FORMATION_INVOLVED_IN_MORPHOGENESIS
GOBP_BLOOD_VESSEL_MORPHOGENESIS
GOBP_COLLAGEN_METABOLIC_PROCESS
HALLMARK_HYPOXIA
HALLMARK_GLYCOLYSIS
GOBP_LEUKOCYTE_MEDIATED_IMMUNITY
GOBP_RESPONSE_TO_WOUNDING
GOBP_POSITIVE_REGULATION_OF_INFLAMMATORY_RESPONSE
GOBP_MYELOID_LEUKOCYTE_ACTIVATION
GOBP_CELL_ACTIVATION
GOBP_COAGULATION
GOBP_TRANSFORMING_GROWTH_FACTOR_BETA_PRODUCTION
GOBP_CIRCULATORY_SYSTEM_DEVELOPMENT
GOBP_CELL_CYCLE
GOBP_CHROMATIN_ORGANIZATION
GOBP_COVALENT_CHROMATIN_MODIFICATION
GOBP_CHROMOSOME_ORGANIZATION
GOBP_DNA_REPLICATION
GOBP_HISTONE_METHYLATION
Pathway
Normalized Enrichment Score
−log10(padj)
GBM endothelial
APP
RHOJ
GLG1
MMP14
SLC35D1
BTBD7
COL4A2
NOTCH3
TMEM131
SIPA1L1
TALDO1
TRNAU1AP
GDF9
TEAD4
RHEBL1
SFR1
RBM14−RBM4
NHLRC4
RBP5
RRH
Spearman's Correlation
Rank
GBM−endothelial: Hallmark & GO pathways NES from GSEA
HALLMARK_EPITHELIAL_MESENCHYMAL_TRANSITION
GOBP_EXTERNAL_ENCAPSULATING_STRUCTURE_ORGANIZATION
GOBP_CIRCULATORY_SYSTEM_DEVELOPMENT
GOBP_CELL_MATRIX_ADHESION
GOBP_BIOLOGICAL_ADHESION
GOBP_VASCULATURE_DEVELOPMENT
GOBP_BLOOD_VESSEL_MORPHOGENESIS
GOBP_REGULATION_OF_VASCULATURE_DEVELOPMENT
GOBP_REGULATION_OF_BLOOD_PRESSURE
GOBP_CELL_SUBSTRATE_ADHESION
GOBP_AMINOGLYCAN_METABOLIC_PROCESS
GOBP_HEART_DEVELOPMENT
GOBP_ANATOMICAL_STRUCTURE_FORMATION_INVOLVED_IN_MORPHOGENESIS
GOBP_BLOOD_VESSEL_ENDOTHELIAL_CELL_MIGRATION
GOBP_NEGATIVE_REGULATION_OF_ERK1_AND_ERK2_CASCADE
GOBP_TUBE_DEVELOPMENT
HALLMARK_UV_RESPONSE_DN
GOBP_CELL_SUBSTRATE_JUNCTION_ORGANIZATION
GOBP_ORGAN_GROWTH
GOBP_NEGATIVE_REGULATION_OF_VASCULATURE_DEVELOPMENT
GOBP_PROTEIN_CONTAINING_COMPLEX_SUBUNIT_ORGANIZATION
GOBP_GENERATION_OF_PRECURSOR_METABOLITES_AND_ENERGY
GOBP_MITOCHONDRIAL_TRANSPORT
GOBP_RNA_MODIFICATION
GOBP_RIBONUCLEOPROTEIN_COMPLEX_BIOGENESIS
GOBP_MITOCHONDRIAL_CYTOCHROME_C_OXIDASE_ASSEMBLY
GOBP_CELLULAR_PROTEIN_CONTAINING_COMPLEX_ASSEMBLY
GOBP_CYTOCHROME_COMPLEX_ASSEMBLY
GOBP_MITOCHONDRION_ORGANIZATION
GOBP_ATP_METABOLIC_PROCESS
GOBP_CELLULAR_RESPIRATION
GOBP_MITOCHONDRIAL_ELECTRON_TRANSPORT_NADH_TO_UBIQUINONE
HALLMARK_MYC_TARGETS_V1
HALLMARK_OXIDATIVE_PHOSPHORYLATION
GOBP_NADH_DEHYDROGENASE_COMPLEX_ASSEMBLY
GOBP_RESPIRATORY_ELECTRON_TRANSPORT_CHAIN
GOBP_MITOCHONDRIAL_RESPIRATORY_CHAIN_COMPLEX_ASSEMBLY
GOBP_ELECTRON_TRANSPORT_CHAIN
GOBP_ATP_SYNTHESIS_COUPLED_ELECTRON_TRANSPORT
GOBP_OXIDATIVE_PHOSPHORYLATION
Pathway
Normalized Enrichment Score
−log10(padj)
GBM T cell
IFI44L
IFI44
RSAD2
IFIT3
IFIT1
HERC6
C5orf56
CMPK2
DDX58
IFI6
ESPN
C16orf45
PCDH1
TPI1
VPS37B
VEGFA
FAM131B
TUBB2B
GAPDH
ACLY
Spearman's Correlation
Rank
GBM−T cell: Hallmark & GO pathways NES from GSEA
HALLMARK_INTERFERON_GAMMA_RESPONSE
HALLMARK_INTERFERON_ALPHA_RESPONSE
GOBP_RESPONSE_TO_VIRUS
GOBP_DEFENSE_RESPONSE_TO_VIRUS
GOBP_INNATE_IMMUNE_RESPONSE
GOBP_RESPONSE_TO_TYPE_I_INTERFERON
GOBP_DEFENSE_RESPONSE_TO_OTHER_ORGANISM
GOBP_RESPONSE_TO_BIOTIC_STIMULUS
GOBP_IMMUNE_EFFECTOR_PROCESS
GOBP_DEFENSE_RESPONSE
GOBP_NEGATIVE_REGULATION_OF_VIRAL_GENOME_REPLICATION
GOBP_VIRAL_GENOME_REPLICATION
GOBP_REGULATION_OF_VIRAL_GENOME_REPLICATION
GOBP_CYTOKINE_MEDIATED_SIGNALING_PATHWAY
GOBP_REGULATION_OF_RESPONSE_TO_BIOTIC_STIMULUS
GOBP_NEGATIVE_REGULATION_OF_VIRAL_PROCESS
GOBP_REGULATION_OF_INNATE_IMMUNE_RESPONSE
GOBP_REGULATION_OF_IMMUNE_EFFECTOR_PROCESS
GOBP_REGULATION_OF_DEFENSE_RESPONSE
GOBP_PATTERN_RECOGNITION_RECEPTOR_SIGNALING_PATHWAY
GOBP_NEURON_DEVELOPMENT
GOBP_NEUROGENESIS
GOBP_CELL_MORPHOGENESIS
GOBP_CELL_JUNCTION_ASSEMBLY
GOBP_GENERATION_OF_PRECURSOR_METABOLITES_AND_ENERGY
GOBP_REGULATION_OF_CELL_JUNCTION_ASSEMBLY
GOBP_SYNAPTIC_SIGNALING
GOBP_CELLULAR_COMPONENT_MORPHOGENESIS
GOBP_POSITIVE_REGULATION_OF_NERVOUS_SYSTEM_DEVELOPMENT
HALLMARK_HYPOXIA
GOBP_TAXIS
GOBP_POSTSYNAPSE_ORGANIZATION
GOBP_REGULATION_OF_NERVOUS_SYSTEM_DEVELOPMENT
GOBP_CELL_PART_MORPHOGENESIS
GOBP_NEURON_PROJECTION_GUIDANCE
GOBP_CELL_MORPHOGENESIS_INVOLVED_IN_DIFFERENTIATION
GOBP_AXON_DEVELOPMENT
GOBP_CELL_JUNCTION_ORGANIZATION
GOBP_SYNAPSE_ORGANIZATION
GOBP_CELL_MORPHOGENESIS_INVOLVED_IN_NEURON_DIFFERENTIATION
Pathway
Normalized Enrichment Score
−log10(padj)
GBM oligo
BCAS1
ZNF488
FA2H
LGI3
RASGRF1
MYRF
TUBB4A
SLC7A14
PPP1R16B
FAM131C
HEATR6
ZCCHC4
WDR89
TM9SF1
SLC52A1
SPATA5L1
ZSCAN22
THAP10
MARCH10
TULP1
Spearman's Correlation
Rank
GBM−oligo: Hallmark & GO pathways NES from GSEA
GOBP_NEUROTRANSMITTER_TRANSPORT
GOBP_VESICLE_MEDIATED_TRANSPORT_IN_SYNAPSE
GOBP_NEUROTRANSMITTER_SECRETION
GOBP_REGULATION_OF_SYNAPTIC_PLASTICITY
GOBP_SYNAPTIC_SIGNALING
GOBP_SYNAPTIC_VESICLE_EXOCYTOSIS
GOBP_REGULATION_OF_NEUROTRANSMITTER_LEVELS
GOBP_REGULATION_OF_TRANS_SYNAPTIC_SIGNALING
GOBP_DICARBOXYLIC_ACID_TRANSPORT
GOBP_LIPID_EXPORT_FROM_CELL
GOBP_ACIDIC_AMINO_ACID_TRANSPORT
GOBP_GLUTAMATE_SECRETION
GOBP_AMINO_ACID_TRANSPORT
GOBP_REGULATION_OF_POSTSYNAPTIC_MEMBRANE_POTENTIAL
GOBP_REGULATION_OF_MEMBRANE_POTENTIAL
GOBP_MONOCARBOXYLIC_ACID_TRANSPORT
GOBP_CALCIUM_ION_REGULATED_EXOCYTOSIS_OF_NEUROTRANSMITTER
GOBP_REGULATION_OF_NEUROTRANSMITTER_TRANSPORT
GOBP_CELL_CELL_SIGNALING
GOBP_SYNAPTIC_VESICLE_PRIMING
GOBP_DNA_CONFORMATION_CHANGE
GOBP_REGULATION_OF_CHROMOSOME_ORGANIZATION
GOBP_NCRNA_PROCESSING
GOBP_DNA_RECOMBINATION
GOBP_DNA_METABOLIC_PROCESS
GOBP_RNA_3_END_PROCESSING
HALLMARK_MYC_TARGETS_V1
GOBP_MITOTIC_CELL_CYCLE_CHECKPOINT
GOBP_CHROMOSOME_SEGREGATION
GOBP_RNA_EXPORT_FROM_NUCLEUS
GOBP_RIBOSOME_BIOGENESIS
GOBP_RNA_PROCESSING
GOBP_CHROMOSOME_ORGANIZATION
GOBP_DNA_INTEGRITY_CHECKPOINT
GOBP_RIBONUCLEOPROTEIN_COMPLEX_BIOGENESIS
GOBP_CELL_CYCLE_CHECKPOINT
GOBP_DNA_DEPENDENT_DNA_REPLICATION
GOBP_DNA_REPLICATION
HALLMARK_G2M_CHECKPOINT
HALLMARK_E2F_TARGETS
Pathway
Normalized Enrichment Score
−log10(padj)

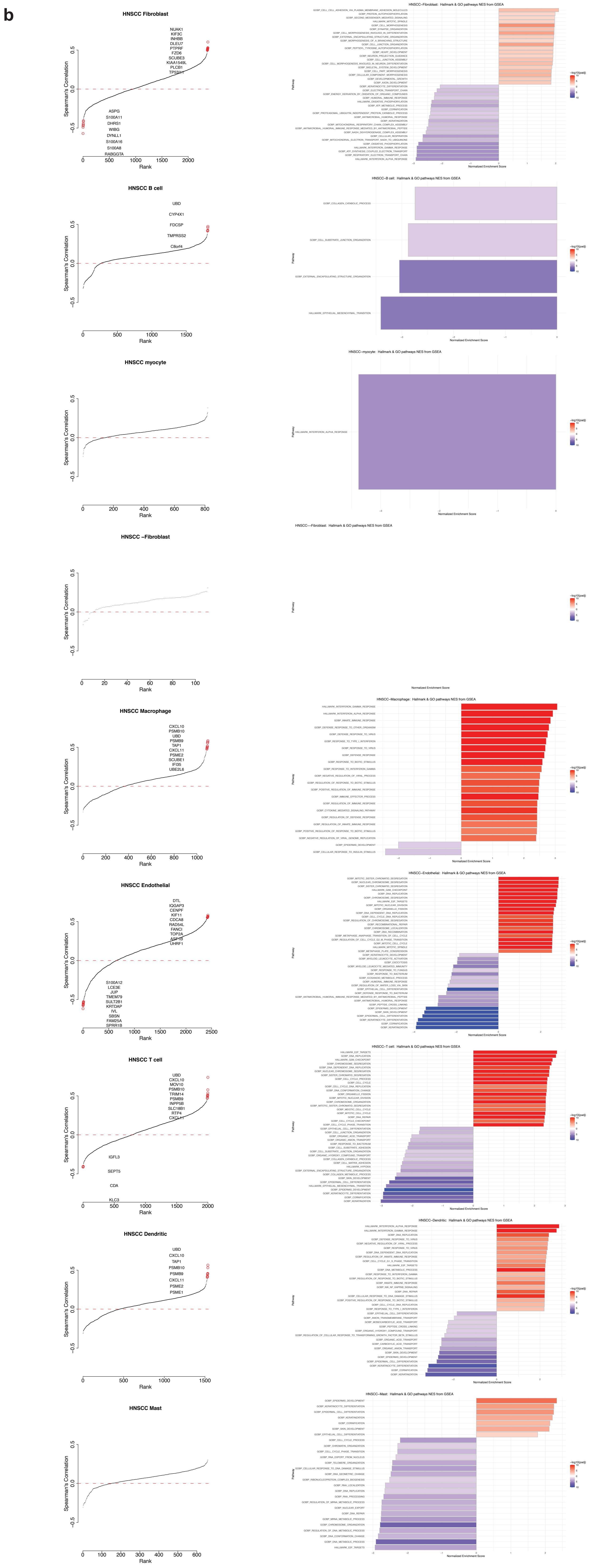
b
HNSCC Fibroblast
NUAK1
KIF3C
INHBB
DLEU7
PTPRF
FZD6
SCUBE3
KIAA1549L
PLCB1
TP53I11
ASPG
S100A11
DHRS1
WIBG
DYNLL1
S100A16
S100A8
RABGGTA
HNSCC B cell
UBD
CYP4X1
FDCSP
TMPRSS2
C8orf4
HNSCC myocyte
HNSCC –Fibroblast
HNSCC Macrophage
CXCL10
PSMB10
UBD
PSMB9
TAP1
CXCL11
PSME2
SCUBE1
IFI35
UBE2L6
HNSCC Endothelial
DTL
IQGAP3
CENPF
KIF11
CDCA8
RAD54L
FANCI
TOP2A
ASF1B
UHRF1
S100A12
LCE3E
JUP
TMEM79
SULT2B1
KRTDAP
IVL
SBSN
FAM25A
SPRR1B
HNSCC T cell
UBD
CXCL10
MOV10
PSMB10
TRIM14
PSMB9
INPP5B
SLC18B1
RTP4
CXCL11
IGFL3
SEPT5
CDA
KLC3
HNSCC Dendritic
UBD
CXCL10
TAP1
PSMB10
PSMB9
CXCL11
PSME2
PSME1
HNSCC Mast
Spearman's Correlation
Rank
HNSCC–Fibroblast: Hallmark & GO pathways NES from GSEA
HNSCC–B cell: Hallmark & GO pathways NES from GSEA
HNSCC–myocyte: Hallmark & GO pathways NES from GSEA
HNSCC—Fibroblast: Hallmark & GO pathways NES from GSEA
HNSCC–Macrophage: Hallmark & GO pathways NES from GSEA
HNSCC–Endothelial: Hallmark & GO pathways NES from GSEA
HNSCC–T cell: Hallmark & GO pathways NES from GSEA
HNSCC–Dendritic: Hallmark & GO pathways NES from GSEA
HNSCC–Mast: Hallmark & GO pathways NES from GSEA
Pathway
Normalized Enrichment Score
–log10(padj)

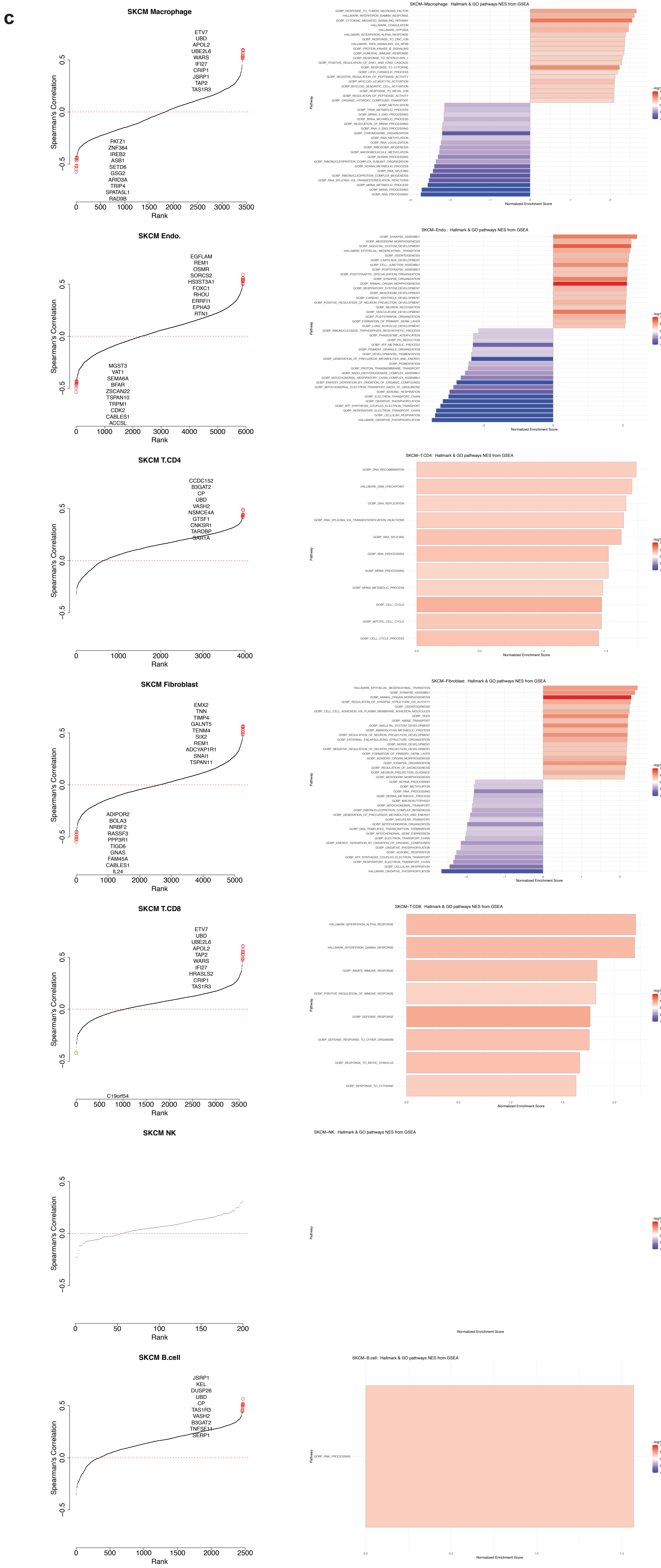


**Fig. S21 | Correlation between malignant cell gene expression and non-malignant cell fraction in three tumor types: TCGA-GBM (a), TCGA-HNSCC (b), and TCGA-SKCM (c).** Rank-ordered plots show Spearman's rank correlation between gene expression in malignant cells inferred by BayesPrism and fractions of each non-malignant cell type in each tumor type. The top 10 positive and negative outlier genes with correlations > 0.4 are marked in red. Barplot shows the normalized gene set enrichment score (NES) for gene sets with the top 20 highest absolute NES, using correlations shown in the corresponding rank-ordered plot as the input.

a

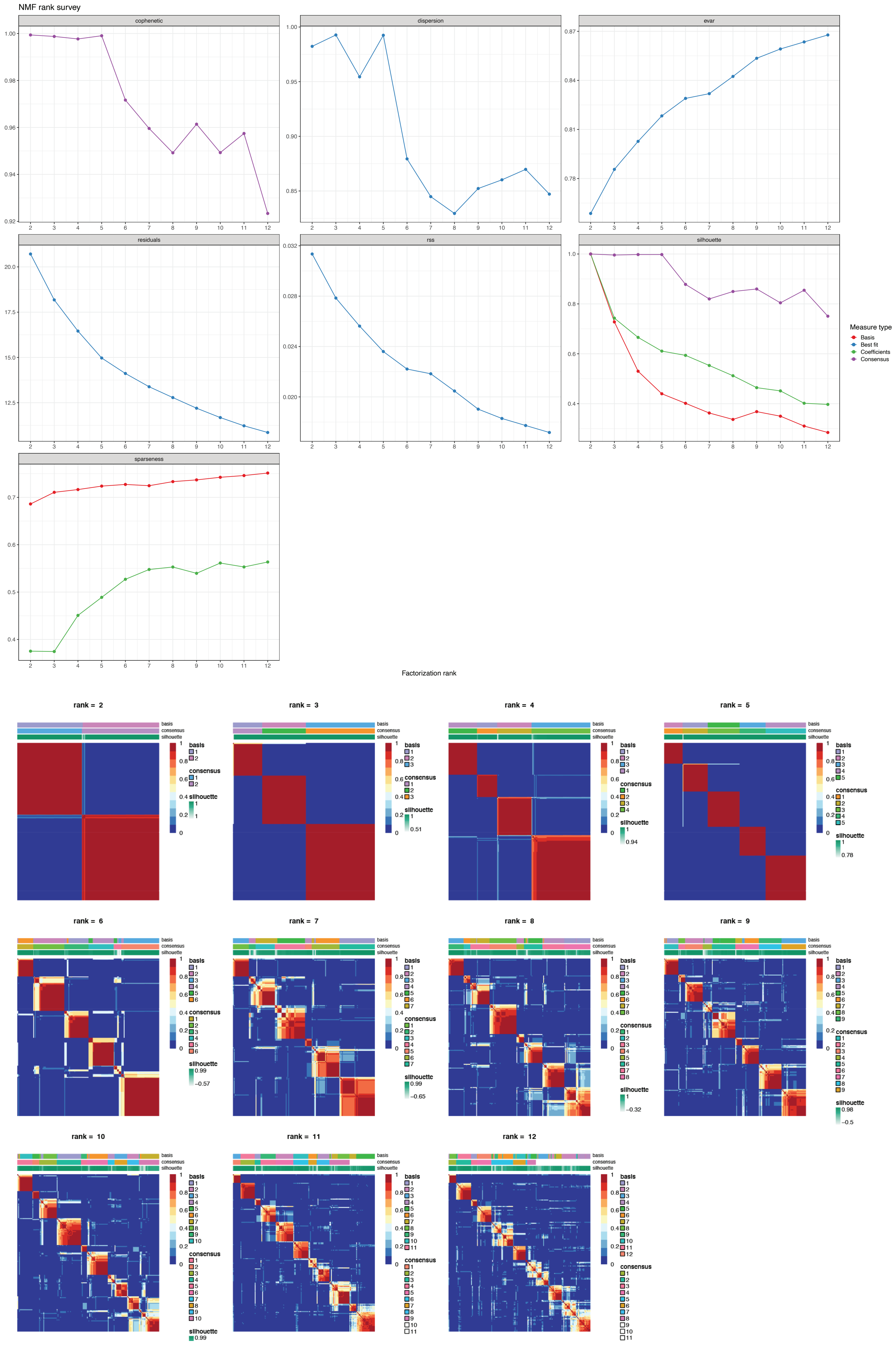

NMF rank survey
cophenetic
dispersion
evar
residuals
rss
silhouette
sparseness
Measure type
Basis
Best fit
Coefficients
Consensus
Factorization rank
rank = 2
rank = 3
rank = 4
rank = 5
rank = 6
rank = 7
rank = 8
rank = 9
rank = 10
rank = 11
rank = 12
basis
consensus
silhouette

b

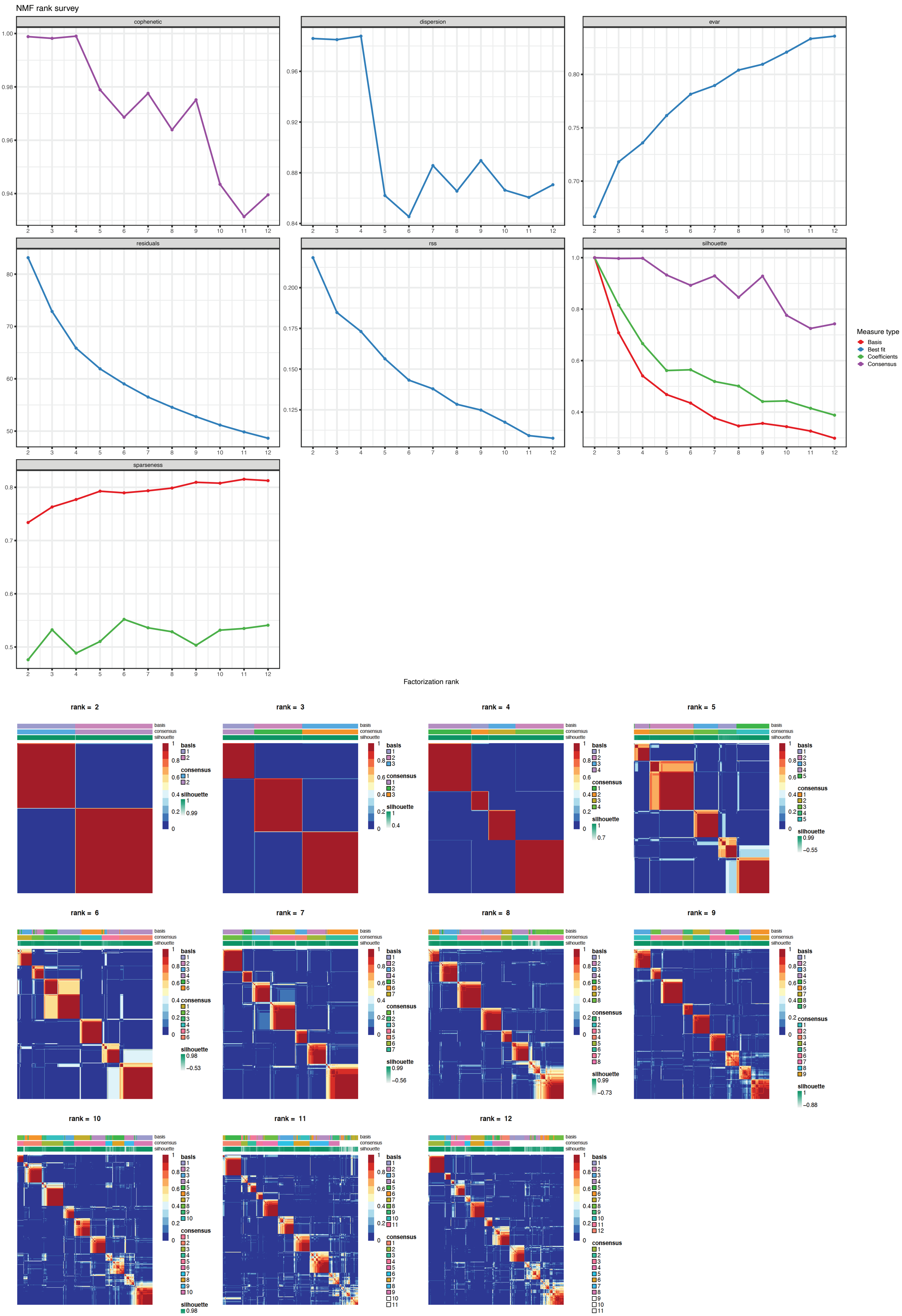

NMF rank survey
cophenetic
dispersion
evar
residuals
rss
silhouette
sparseness
Measure type
Basis
Best fit
Coefficients
Consensus
Factorization rank
rank = 2
rank = 3
rank = 4
rank = 5
rank = 6
rank = 7
rank = 8
rank = 9
rank = 10
rank = 11
rank = 12
basis
consensus
silhouette

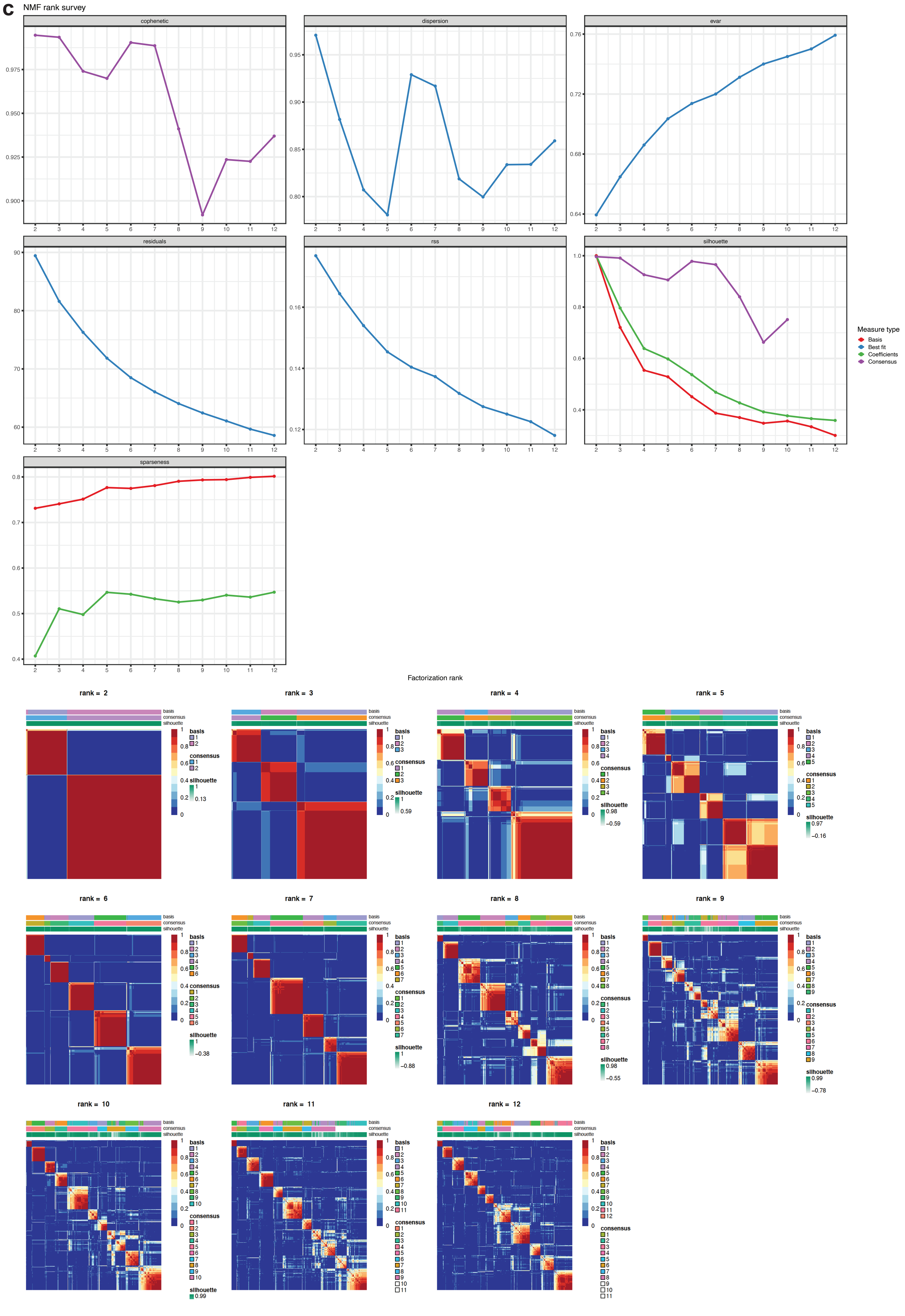


**Fig. S22 | Choosing the number of gene programs and initializing tumor basis $\eta_0$ using NMF factorization and consensus clustering for embedding learning in three tumor types: TCGA-GBM (a), TCGA-HNSCC (b), and TCGA-SKCM (c).** Line plots show various metrics on consensus clustering as a function of the number of gene programs. Heatmaps show the consensus clustering matrix of different choices of number of gene programs.

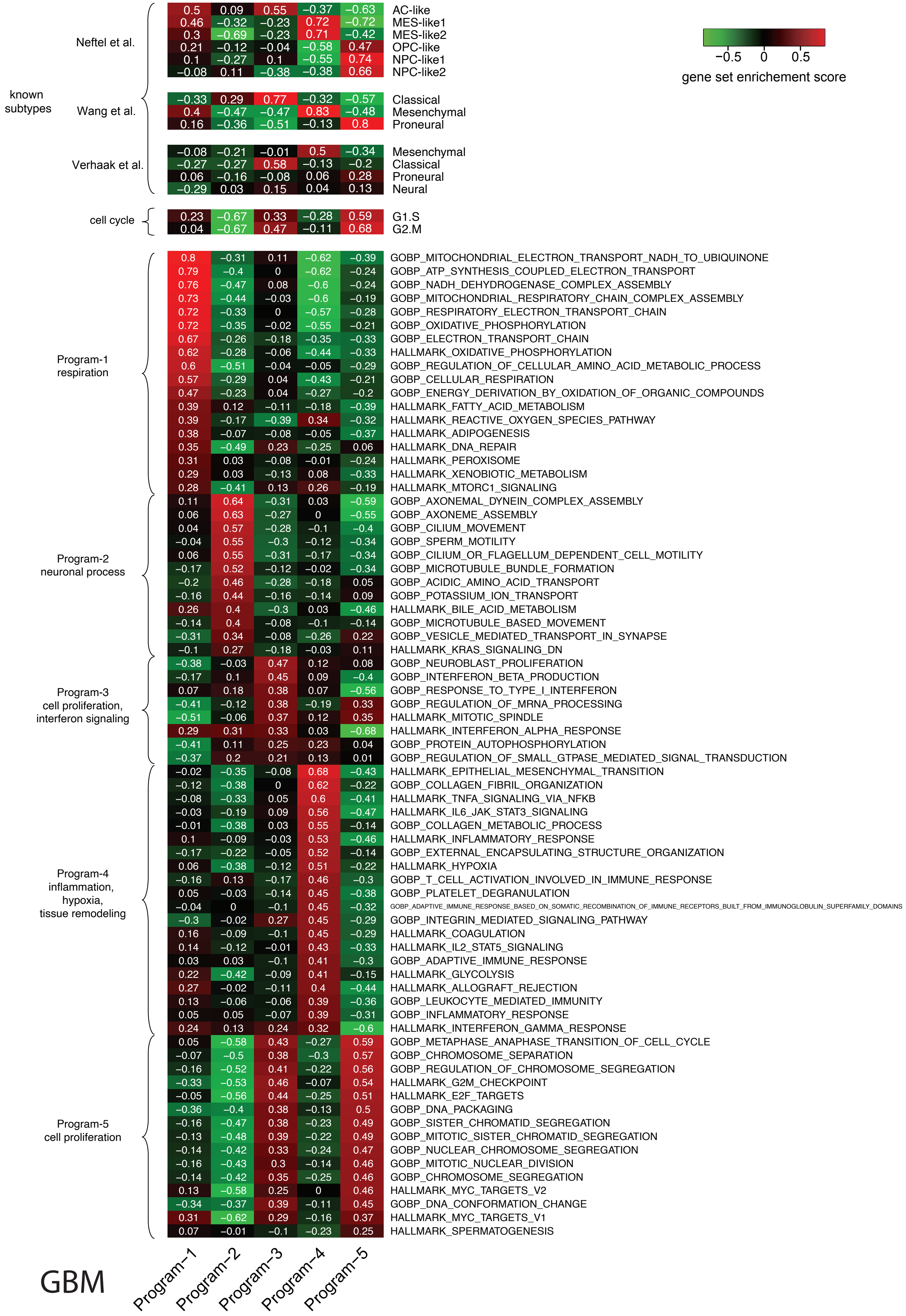

known subtypes
Neftel et al.
0.5 0.09 0.55 −0.37 −0.63 AC-like
0.46 −0.32 −0.23 0.72 −0.72 MES-like1
0.3 −0.69 −0.23 0.71 −0.42 MES-like2
0.21 −0.12 −0.04 −0.58 0.47 OPC-like
0.1 −0.27 0.1 −0.55 0.74 NPC-like1
−0.08 0.11 −0.38 −0.38 0.66 NPC-like2
Wang et al.
−0.33 0.29 0.77 −0.32 −0.57 Classical
0.4 −0.47 −0.47 0.83 −0.48 Mesenchymal
0.16 −0.36 −0.51 −0.13 0.8 Proneural
Verhaak et al.
−0.08 −0.21 −0.01 0.5 −0.34 Mesenchymal
−0.27 −0.27 0.58 −0.13 −0.2 Classical
0.06 −0.16 −0.08 0.06 0.28 Proneural
−0.29 0.03 0.15 0.04 0.13 Neural
cell cycle
0.23 −0.67 0.33 −0.28 0.59 G1.S
0.04 −0.67 0.47 −0.11 0.68 G2.M
Program-1 respiration
0.8 −0.31 0.11 −0.62 −0.39 GOBP_MITOCHONDRIAL_ELECTRON_TRANSPORT_NADH_TO_UBIQUINONE
0.79 −0.4 0 −0.62 −0.24 GOBP_ATP_SYNTHESIS_COUPLED_ELECTRON_TRANSPORT
0.76 −0.47 0.08 −0.6 −0.24 GOBP_NADH_DEHYDROGENASE_COMPLEX_ASSEMBLY
0.73 −0.44 −0.03 −0.6 −0.19 GOBP_MITOCHONDRIAL_RESPIRATORY_CHAIN_COMPLEX_ASSEMBLY
0.72 −0.33 0 −0.57 −0.28 GOBP_RESPIRATORY_ELECTRON_TRANSPORT_CHAIN
0.72 −0.35 −0.02 −0.55 −0.21 GOBP_OXIDATIVE_PHOSPHORYLATION
0.67 −0.26 −0.18 −0.35 −0.33 GOBP_ELECTRON_TRANSPORT_CHAIN
0.62 −0.28 −0.06 −0.44 −0.33 HALLMARK_OXIDATIVE_PHOSPHORYLATION
0.6 −0.51 −0.04 −0.05 −0.29 GOBP_REGULATION_OF_CELLULAR_AMINO_ACID_METABOLIC_PROCESS
0.57 −0.29 0.04 −0.43 −0.21 GOBP_CELLULAR_RESPIRATION
0.47 −0.23 0.04 −0.27 −0.2 GOBP_ENERGY_DERIVATION_BY_OXIDATION_OF_ORGANIC_COMPOUNDS
0.39 0.12 −0.11 −0.18 −0.39 HALLMARK_FATTY_ACID_METABOLISM
0.39 −0.17 −0.39 0.34 −0.32 HALLMARK_REACTIVE_OXYGEN_SPECIES_PATHWAY
0.38 −0.07 −0.08 −0.05 −0.37 HALLMARK_ADIPOGENESIS
0.35 −0.49 0.23 −0.25 0.06 HALLMARK_DNA_REPAIR
0.31 0.03 −0.08 −0.01 −0.24 HALLMARK_PEROXISOME
0.29 0.03 −0.13 0.08 −0.33 HALLMARK_XENOBIOTIC_METABOLISM
0.28 −0.41 0.13 0.26 −0.19 HALLMARK_MTORC1_SIGNALING
Program-2 neuronal process
0.11 0.64 −0.31 0.03 −0.59 GOBP_AXONEMAL_DYNEIN_COMPLEX_ASSEMBLY
0.06 0.63 −0.27 0 −0.55 GOBP_AXONEME_ASSEMBLY
0.04 0.57 −0.28 −0.1 −0.4 GOBP_CILIUM_MOVEMENT
−0.04 0.55 −0.3 −0.12 −0.34 GOBP_SPERM_MOTILITY
0.06 0.55 −0.31 −0.17 −0.34 GOBP_CILIUM_OR_FLAGELLUM_DEPENDENT_CELL_MOTILITY
−0.17 0.52 −0.12 −0.02 −0.34 GOBP_MICROTUBULE_BUNDLE_FORMATION
−0.2 0.46 −0.28 −0.18 0.05 GOBP_ACIDIC_AMINO_ACID_TRANSPORT
−0.16 0.44 −0.16 −0.14 0.09 GOBP_POTASSIUM_ION_TRANSPORT
0.26 0.4 −0.3 0.03 −0.46 HALLMARK_BILE_ACID_METABOLISM
−0.14 0.4 −0.08 −0.1 −0.14 GOBP_MICROTUBULE_BASED_MOVEMENT
−0.31 0.34 −0.08 −0.26 0.22 GOBP_VESICLE_MEDIATED_TRANSPORT_IN_SYNAPSE
−0.1 0.27 −0.18 −0.03 0.11 HALLMARK_KRAS_SIGNALING_DN
Program-3 cell proliferation, interferon signaling
−0.38 −0.03 0.47 0.12 0.08 GOBP_NEUROBLAST_PROLIFERATION
−0.17 0.1 0.45 0.09 −0.4 GOBP_INTERFERON_BETA_PRODUCTION
0.07 0.18 0.38 0.07 −0.56 GOBP_RESPONSE_TO_TYPE_I_INTERFERON
−0.41 −0.12 0.38 −0.19 0.33 GOBP_REGULATION_OF_MRNA_PROCESSING
−0.51 −0.06 0.37 0.12 0.35 HALLMARK_MITOTIC_SPINDLE
0.29 0.31 0.33 0.03 −0.68 HALLMARK_INTERFERON_ALPHA_RESPONSE
−0.41 0.11 0.25 0.23 0.04 GOBP_PROTEIN_AUTOPHOSPHORYLATION
−0.37 0.2 0.21 0.13 0.01 GOBP_REGULATION_OF_SMALL_GTPASE_MEDIATED_SIGNAL_TRANSDUCTION
Program-4 inflammation, hypoxia, tissue remodeling
−0.02 −0.35 −0.08 0.68 −0.43 HALLMARK_EPITHELIAL_MESENCHYMAL_TRANSITION
−0.12 −0.38 0 0.62 −0.22 GOBP_COLLAGEN_FIBRIL_ORGANIZATION
−0.08 −0.33 0.05 0.6 −0.41 HALLMARK_TNFA_SIGNALING_VIA_NFKB
−0.03 −0.19 0.09 0.56 −0.47 HALLMARK_IL6_JAK_STAT3_SIGNALING
−0.01 −0.38 0.03 0.55 −0.14 GOBP_COLLAGEN_METABOLIC_PROCESS
0.1 −0.09 −0.03 0.53 −0.46 HALLMARK_INFLAMMATORY_RESPONSE
−0.17 −0.22 −0.05 0.52 −0.14 GOBP_EXTERNAL_ENCAPSULATING_STRUCTURE_ORGANIZATION
0.06 −0.38 −0.12 0.51 −0.22 HALLMARK_HYPOXIA
−0.16 0.13 −0.17 0.46 −0.3 GOBP_T_CELL_ACTIVATION_INVOLVED_IN_IMMUNE_RESPONSE
0.05 −0.03 −0.14 0.45 −0.38 GOBP_PLATELET_DEGRANULATION
−0.04 0 −0.1 0.45 −0.32 GOBP_ADAPTIVE_IMMUNE_RESPONSE_BASED_ON_SOMATIC_RECOMBINATION_OF_IMMUNE_RECEPTORS_BUILT_FROM_IMMUNOGLOBULIN_SUPERFAMILY_DOMAINS
−0.3 −0.02 0.27 0.45 −0.29 GOBP_INTEGRIN_MEDIATED_SIGNALING_PATHWAY
0.16 −0.09 −0.1 0.45 −0.29 HALLMARK_COAGULATION
0.14 −0.12 −0.01 0.43 −0.33 HALLMARK_IL2_STAT5_SIGNALING
0.03 0.03 −0.1 0.41 −0.3 GOBP_ADAPTIVE_IMMUNE_RESPONSE
0.22 −0.42 −0.09 0.41 −0.15 HALLMARK_GLYCOLYSIS
0.27 −0.02 −0.11 0.4 −0.44 HALLMARK_ALLOGRAFT_REJECTION
0.13 −0.06 −0.06 0.39 −0.36 GOBP_LEUKOCYTE_MEDIATED_IMMUNITY
0.05 0.05 −0.07 0.39 −0.31 GOBP_INFLAMMATORY_RESPONSE
0.24 0.13 0.24 0.32 −0.6 HALLMARK_INTERFERON_GAMMA_RESPONSE
Program-5 cell proliferation
0.05 −0.58 0.43 −0.27 0.59 GOBP_METAPHASE_ANAPHASE_TRANSITION_OF_CELL_CYCLE
−0.07 −0.5 0.38 −0.3 0.57 GOBP_CHROMOSOME_SEPARATION
−0.16 −0.52 0.41 −0.22 0.56 GOBP_REGULATION_OF_CHROMOSOME_SEGREGATION
−0.33 −0.53 0.46 −0.07 0.54 HALLMARK_G2M_CHECKPOINT
−0.05 −0.56 0.44 −0.25 0.51 HALLMARK_E2F_TARGETS
−0.36 −0.4 0.38 −0.13 0.5 GOBP_DNA_PACKAGING
−0.16 −0.47 0.38 −0.23 0.49 GOBP_SISTER_CHROMATID_SEGREGATION
−0.13 −0.48 0.39 −0.22 0.49 GOBP_MITOTIC_SISTER_CHROMATID_SEGREGATION
−0.14 −0.42 0.33 −0.24 0.47 GOBP_NUCLEAR_CHROMOSOME_SEGREGATION
−0.16 −0.43 0.3 −0.14 0.46 GOBP_MITOTIC_NUCLEAR_DIVISION
−0.14 −0.42 0.35 −0.25 0.46 GOBP_CHROMOSOME_SEGREGATION
0.13 −0.58 0.25 0 0.46 HALLMARK_MYC_TARGETS_V2
−0.34 −0.37 0.39 −0.11 0.45 GOBP_DNA_CONFORMATION_CHANGE
0.31 −0.62 0.29 −0.16 0.37 HALLMARK_MYC_TARGETS_V1
0.07 −0.01 −0.1 −0.23 0.25 HALLMARK_SPERMATOGENESIS
Program−1 Program−2 Program−3 Program−4 Program−5
GBM
−0.5 0 0.5
gene set enrichement score

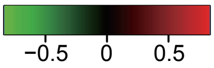


| Group | Gene set | Program−1 | Program−2 | Program−3 | Program−4 |
|---|---|---|---|---|---|
| subtypes defined by bulk RNA-seq | Basal | 0.51 | −0.66 | −0.38 | 0.56 |
| | Mesenchymal | 0.06 | −0.35 | 0.26 | 0.28 |
| | Atypical | −0.53 | 0.43 | −0.21 | 0.45 |
| | Classical | 0.18 | 0.58 | −0.21 | −0.46 |
| gene programs defined by scRNA-seq | Cell Cycle | 0.22 | −0.03 | 0.42 | −0.69 |
| | partial EMT | 0.67 | −0.68 | 0.12 | 0.04 |
| | Epithelial_differentiation_1 | −0.41 | −0.54 | −0.32 | 0.8 |
| | Epithelial_differentiation_2 | −0.25 | −0.59 | 0.48 | 0.44 |
| | Stress | −0.1 | −0.49 | −0.11 | 0.51 |
| | Hypoxia | 0.23 | −0.38 | −0.19 | 0.2 |
| Program-1 tissue remodeling, partial EMT | GOBP_HEMIDESMOSOME_ASSEMBLY | 0.89 | −0.62 | −0.27 | −0.07 |
| | GOBP_REGULATION_OF_VITAMIN_D_BIOSYNTHETIC_PROCESS | 0.82 | −0.75 | 0.06 | −0.14 |
| | GOBP_ENTRY_OF_BACTERIUM_INTO_HOST_CELL | 0.66 | −0.2 | −0.6 | 0.08 |
| | GOBP_WOUND_HEALING_SPREADING_OF_EPIDERMAL_CELLS | 0.61 | −0.42 | −0.63 | 0.32 |
| | GOBP_POSITIVE_REGULATION_OF_EXTRINSIC_APOPTOTIC_SIGNALING_PATHWAY_VIA_DEATH_DOMAIN_REC | 0.59 | −0.64 | −0.15 | 0.22 |
| | GOBP_BASEMENT_MEMBRANE_ORGANIZATION | 0.53 | −0.42 | −0.19 | −0.01 |
| | GOBP_CELL_SUBSTRATE_JUNCTION_ORGANIZATION | 0.52 | −0.36 | −0.43 | 0.31 |
| | GOBP_CELL_ADHESION_MEDIATED_BY_INTEGRIN | 0.5 | −0.31 | −0.27 | 0 |
| | HALLMARK_MITOTIC_SPINDLE | 0.5 | −0.1 | −0.43 | −0.12 |
| | HALLMARK_EPITHELIAL_MESENCHYMAL_TRANSITION | 0.48 | −0.43 | 0.03 | 0.14 |
| | GOBP_FOCAL_ADHESION_ASSEMBLY | 0.48 | −0.35 | −0.44 | 0.34 |
| | HALLMARK_PROTEIN_SECRETION | 0.48 | −0.28 | −0.41 | 0.05 |
| | GOBP_EPIBOLY | 0.47 | −0.36 | −0.43 | 0.13 |
| | HALLMARK_ANGIOGENESIS | 0.44 | −0.4 | −0.03 | 0.1 |
| | HALLMARK_G2M_CHECKPOINT | 0.4 | 0.14 | −0.24 | −0.49 |
| | HALLMARK_UV_RESPONSE_DN | 0.4 | −0.11 | −0.44 | 0.31 |
| | HALLMARK_TGF_BETA_SIGNALING | 0.39 | −0.44 | −0.32 | 0.24 |
| | HALLMARK_E2F_TARGETS | 0.37 | 0.15 | 0.21 | −0.62 |
| | HALLMARK_IL6_JAK_STAT3_SIGNALING | 0.35 | −0.52 | 0.04 | 0.23 |
| | HALLMARK_APICAL_JUNCTION | 0.31 | −0.37 | −0.18 | 0.15 |
| | HALLMARK_TNFA_SIGNALING_VIA_NFKB | 0.28 | −0.5 | −0.03 | 0.27 |
| | HALLMARK_MYC_TARGETS_V1 | 0.27 | −0.23 | 0.27 | −0.53 |
| Program-2 metabolic process, Classical-like | GOBP_FLAVONOID_METABOLIC_PROCESS | −0.36 | 0.75 | −0.02 | 0.14 |
| | GOBP_SKELETAL_MUSCLE_SATELLITE_CELL_DIFFERENTIATION | −0.05 | 0.71 | −0.37 | 0.09 |
| | GOBP_ETHANOL_OXIDATION | −0.62 | 0.7 | 0.01 | 0.05 |
| | GOBP_CELLULAR_GLUCURONIDATION | −0.26 | 0.69 | 0 | −0.02 |
| | GOBP_LATERAL_SPROUTING_FROM_AN_EPITHELIUM | 0.07 | 0.68 | −0.34 | 0.2 |
| | GOBP_URONIC_ACID_METABOLIC_PROCESS | −0.39 | 0.61 | 0.05 | 0.09 |
| | GOBP_DETECTION_OF_CHEMICAL_STIMULUS_INVOLVED_IN_SENSORY_PERCEPTION_OF_TASTE | −0.53 | 0.51 | 0.18 | 0.04 |
| | GOBP_CILIUM_MOVEMENT | −0.33 | 0.36 | −0.04 | −0.1 |
| | GOBP_HISTONE_H4_ACETYLATION | −0.03 | 0.3 | −0.08 | −0.33 |
| | HALLMARK_SPERMATOGENESIS | −0.14 | 0.28 | −0.06 | −0.24 |
| | GOBP_INTERSTRAND_CROSS_LINK_REPAIR | 0.18 | 0.22 | −0.03 | −0.54 |
| Program-3 respiration, Myc activation, cell cycle, interferon signaling | GOBP_MITOCHONDRIAL_ELECTRON_TRANSPORT_NADH_TO_UBIQUINONE | −0.59 | −0.44 | 0.68 | 0.14 |
| | GOBP_NADH_DEHYDROGENASE_COMPLEX_ASSEMBLY | −0.61 | −0.32 | 0.65 | −0.14 |
| | GOBP_ATP_SYNTHESIS_COUPLED_ELECTRON_TRANSPORT | −0.55 | −0.28 | 0.65 | −0.06 |
| | GOBP_MITOCHONDRIAL_RESPIRATORY_CHAIN_COMPLEX_ASSEMBLY | −0.55 | −0.27 | 0.63 | −0.27 |
| | GOBP_RESPIRATORY_ELECTRON_TRANSPORT_CHAIN | −0.51 | −0.28 | 0.6 | −0.17 |
| | GOBP_OXIDATIVE_PHOSPHORYLATION | −0.49 | −0.32 | 0.6 | −0.07 |
| | GOBP_CYTOCHROME_COMPLEX_ASSEMBLY | −0.44 | −0.12 | 0.56 | −0.53 |
| | GOBP_ELECTRON_TRANSPORT_CHAIN | −0.45 | −0.3 | 0.54 | −0.06 |
| | GOBP_REGULATION_OF_CELLULAR_AMINO_ACID_METABOLIC_PROCESS | 0.08 | −0.31 | 0.5 | −0.29 |
| | HALLMARK_INTERFERON_ALPHA_RESPONSE | 0.24 | −0.71 | 0.49 | 0.35 |
| | HALLMARK_OXIDATIVE_PHOSPHORYLATION | −0.43 | −0.16 | 0.46 | −0.11 |
| | GOBP_CELLULAR_RESPIRATION | −0.42 | −0.19 | 0.44 | −0.1 |
| | HALLMARK_MYC_TARGETS_V2 | 0.27 | −0.21 | 0.44 | −0.62 |
| | HALLMARK_DNA_REPAIR | −0.03 | −0.24 | 0.41 | −0.41 |
| | HALLMARK_REACTIVE_OXYGEN_SPECIES_PATHWAY | −0.4 | −0.06 | 0.23 | 0.17 |
| | HALLMARK_FATTY_ACID_METABOLISM | −0.29 | 0.07 | 0.12 | 0.12 |
| Program-4 immune/stress-response, epithelial differentiation | GOBP_PEPTIDE_CROSS_LINKING | 0.01 | −0.53 | −0.27 | 0.77 |
| | GOBP_CORNIFICATION | −0.2 | −0.51 | −0.29 | 0.76 |
| | GOBP_KERATINIZATION | −0.25 | −0.48 | −0.24 | 0.73 |
| | GOBP_REGULATION_OF_WATER_LOSS_VIA_SKIN | 0.03 | −0.44 | −0.4 | 0.64 |
| | GOBP_KERATINOCYTE_DIFFERENTIATION | 0.08 | −0.42 | −0.35 | 0.62 |
| | GOBP_EPIDERMAL_CELL_DIFFERENTIATION | 0.03 | −0.32 | −0.31 | 0.55 |
| | GOBP_SKIN_DEVELOPMENT | 0.24 | −0.27 | −0.35 | 0.51 |
| | GOBP_EPIDERMIS_DEVELOPMENT | 0.18 | −0.25 | −0.31 | 0.48 |
| | GOBP_POSITIVE_REGULATION_OF_MACROAUTOPHAGY | 0.29 | −0.34 | −0.47 | 0.46 |
| | GOBP_WATER_HOMEOSTASIS | 0.02 | −0.11 | −0.27 | 0.44 |
| | HALLMARK_INFLAMMATORY_RESPONSE | 0.35 | −0.55 | −0.02 | 0.41 |
| | HALLMARK_INTERFERON_GAMMA_RESPONSE | 0.26 | −0.58 | 0.31 | 0.35 |
| | HALLMARK_ALLOGRAFT_REJECTION | 0.11 | −0.49 | 0.25 | 0.29 |
| | HALLMARK_COMPLEMENT | 0.24 | −0.39 | 0.06 | 0.28 |
| | HALLMARK_KRAS_SIGNALING_UP | 0.23 | −0.26 | −0.11 | 0.26 |
| | HALLMARK_ADIPOGENESIS | −0.24 | −0.05 | 0.07 | 0.11 |

HNSCC

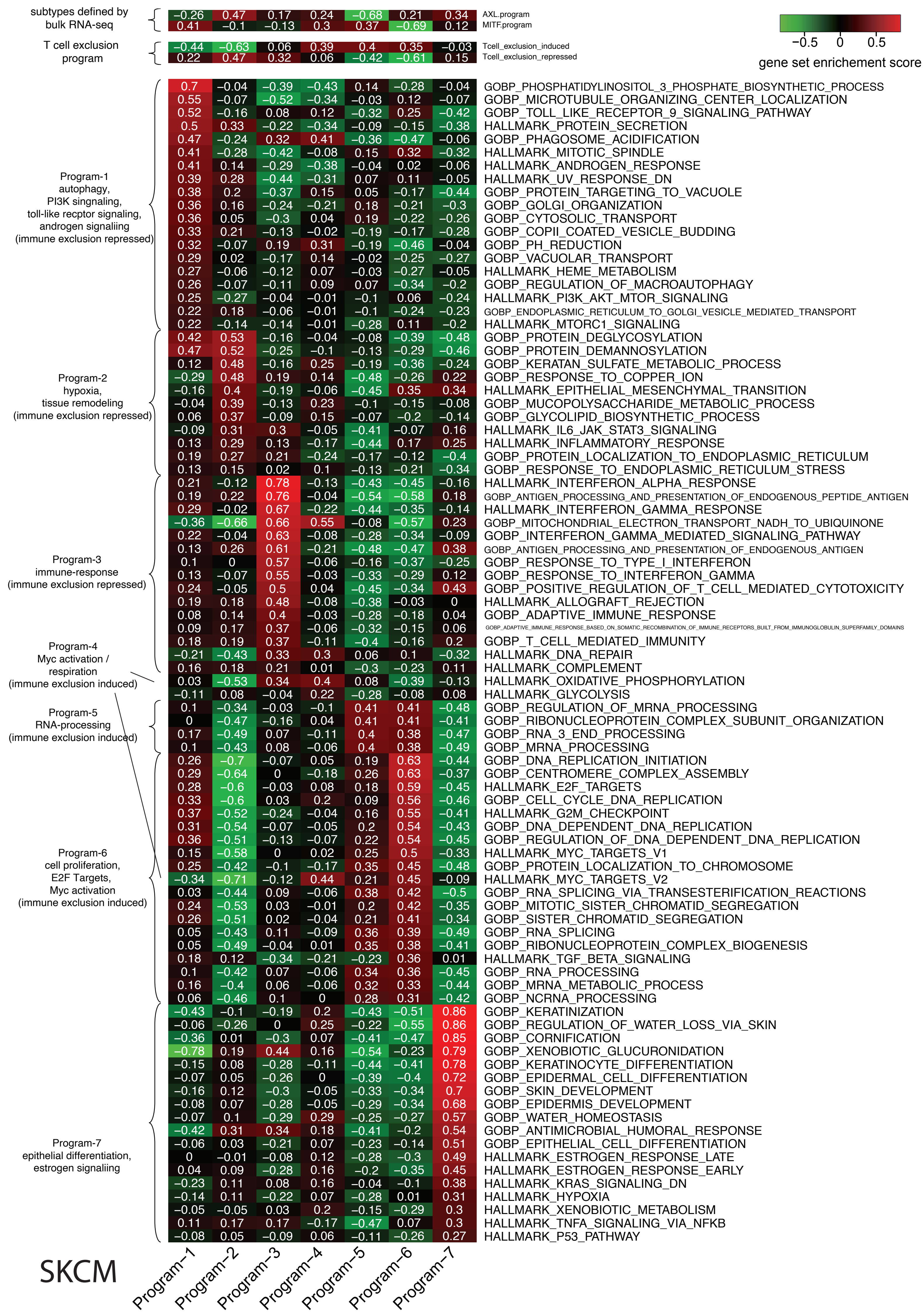


**Fig. S23 | Heatmap shows the gene set enrichment score by GSVA for subtype marker genes and MgSigDB biological process for each gene program of TCGA-GBM (a), TCGA-HNSCC (b), and TCGA-SKCM (c) inferred by BayesPrism.**

**a**

### TCGA-GBM

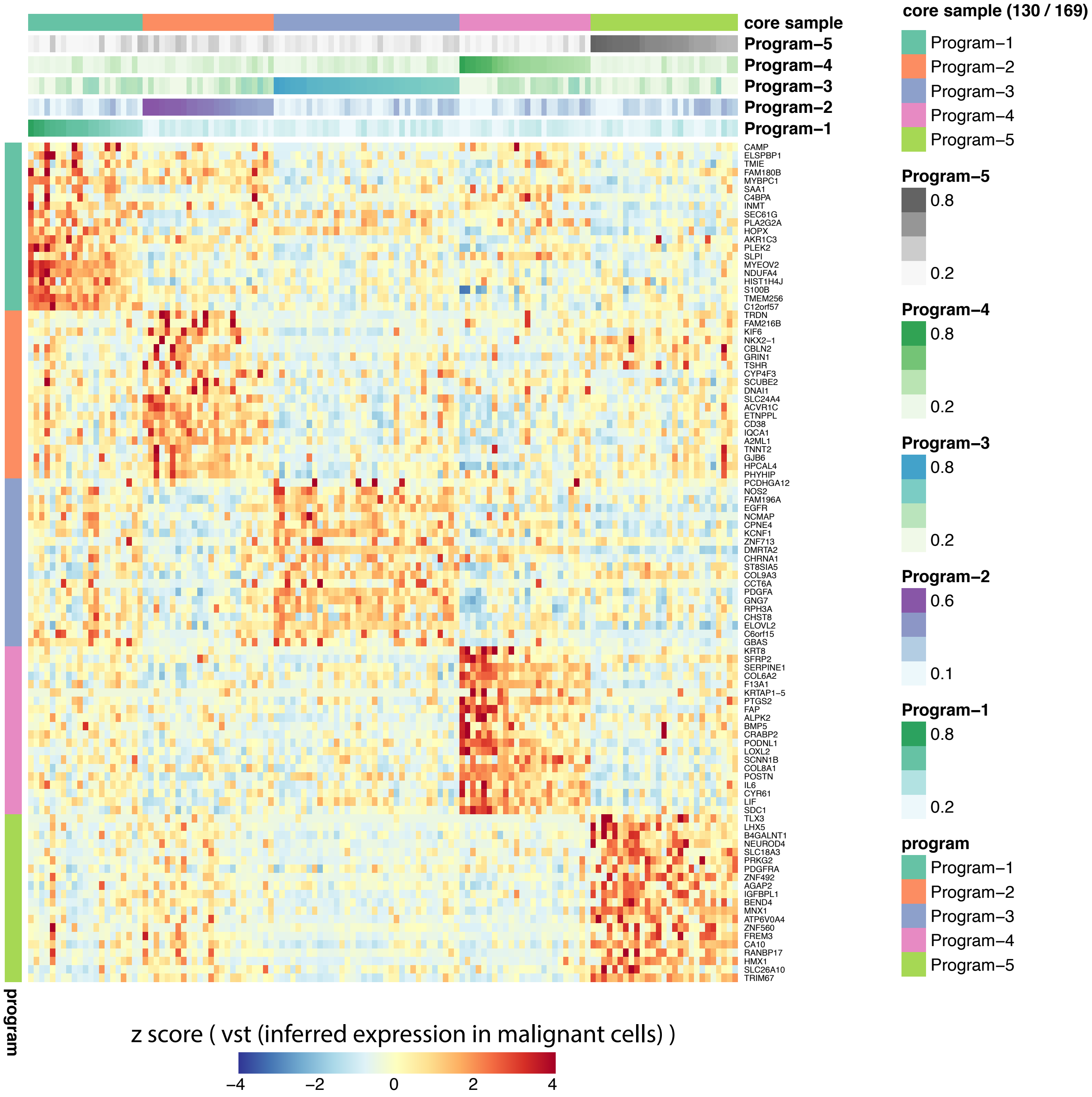
core sample
Program−5
Program−4
Program−3
Program−2
Program−1
CAMP
ELSPBP1
TMIE
FAM180B
MYBPC1
SAA1
C4BPA
INMT
SEC61G
PLA2G2A
HOPX
AKR1C3
PLEK2
SLPI
MYEOV2
NDUFA4
HIST1H4J
S100B
TMEM256
C12orf57
TRDN
FAM216B
KIF6
NKX2−1
CBLN2
GRIN1
TSHR
CYP4F3
SCUBE2
DNAI1
SLC24A4
ACVR1C
ETNPPL
CD38
IQCA1
A2ML1
TNNT2
GJB6
HPCAL4
PHYHIP
PCDHGA12
NOS2
FAM196A
EGFR
NCMAP
CPNE4
KCNF1
ZNF713
DMRTA2
CHRNA1
ST8SIA5
COL9A3
CCT6A
PDGFA
GNG7
RPH3A
CHST8
ELOVL2
C6orf15
GBAS
KRT8
SFRP2
SERPINE1
COL6A2
F13A1
KRTAP1−5
PTGS2
FAP
ALPK2
BMP5
CRABP2
PODNL1
LOXL2
SCNN1B
COL8A1
POSTN
IL6
CYR61
LIF
SDC1
TLX3
LHX5
B4GALNT1
NEUROD4
SLC18A3
PRKG2
PDGFRA
ZNF492
AGAP2
IGFBPL1
BEND4
MNX1
ATP6V0A4
ZNF560
FREM3
CA10
RANBP17
HMX1
SLC26A10
TRIM67
program
z score ( vst (inferred expression in malignant cells) )
−4
−2
0
2
4
core sample (130 / 169)
Program−1
Program−2
Program−3
Program−4
Program−5
Program−5
0.8
0.2
Program−4
0.8
0.2
Program−3
0.8
0.2
Program−2
0.6
0.1
Program−1
0.8
0.2
program
Program−1
Program−2
Program−3
Program−4
Program−5

**b**

### TCGA-HNSCC

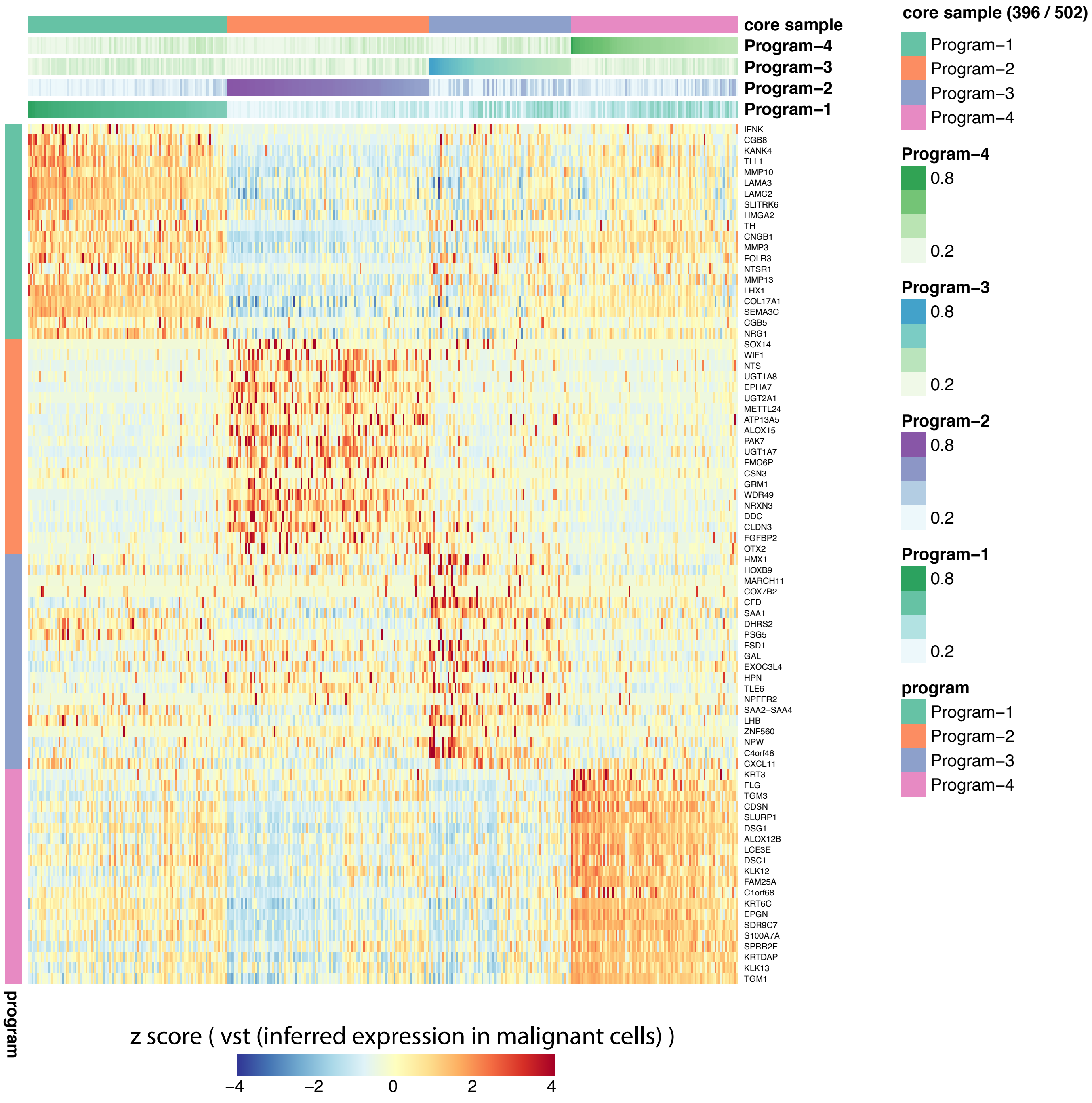

core sample
Program−4
Program−3
Program−2
Program−1
IFNK
CGB8
KANK4
TLL1
MMP10
LAMA3
LAMC2
SLITRK6
HMGA2
TH
CNGB1
MMP3
FOLR3
NTSR1
MMP13
LHX1
COL17A1
SEMA3C
CGB5
NRG1
SOX14
WIF1
NTS
UGT1A8
EPHA7
UGT2A1
METTL24
ATP13A5
ALOX15
PAK7
UGT1A7
FMO6P
CSN3
GRM1
WDR49
NRXN3
DDC
CLDN3
FGFBP2
OTX2
HMX1
HOXB9
MARCH11
COX7B2
CFD
SAA1
DHRS2
PSG5
FSD1
GAL
EXOC3L4
HPN
TLE6
NPFFR2
SAA2−SAA4
LHB
ZNF560
NPW
C4orf48
CXCL11
KRT3
FLG
TGM3
CDSN
SLURP1
DSG1
ALOX12B
LCE3E
DSC1
KLK12
FAM25A
C1orf68
KRT6C
EPGN
SDR9C7
S100A7A
SPRR2F
KRTDAP
KLK13
TGM1
program
core sample (396 / 502)
Program−1
Program−2
Program−3
Program−4
Program−4
0.8
0.2
Program−3
0.8
0.2
Program−2
0.8
0.2
Program−1
0.8
0.2
program
Program−1
Program−2
Program−3
Program−4
z score ( vst (inferred expression in malignant cells) )
−4
−2
0
2
4

**c**

### TCGA-SKCM

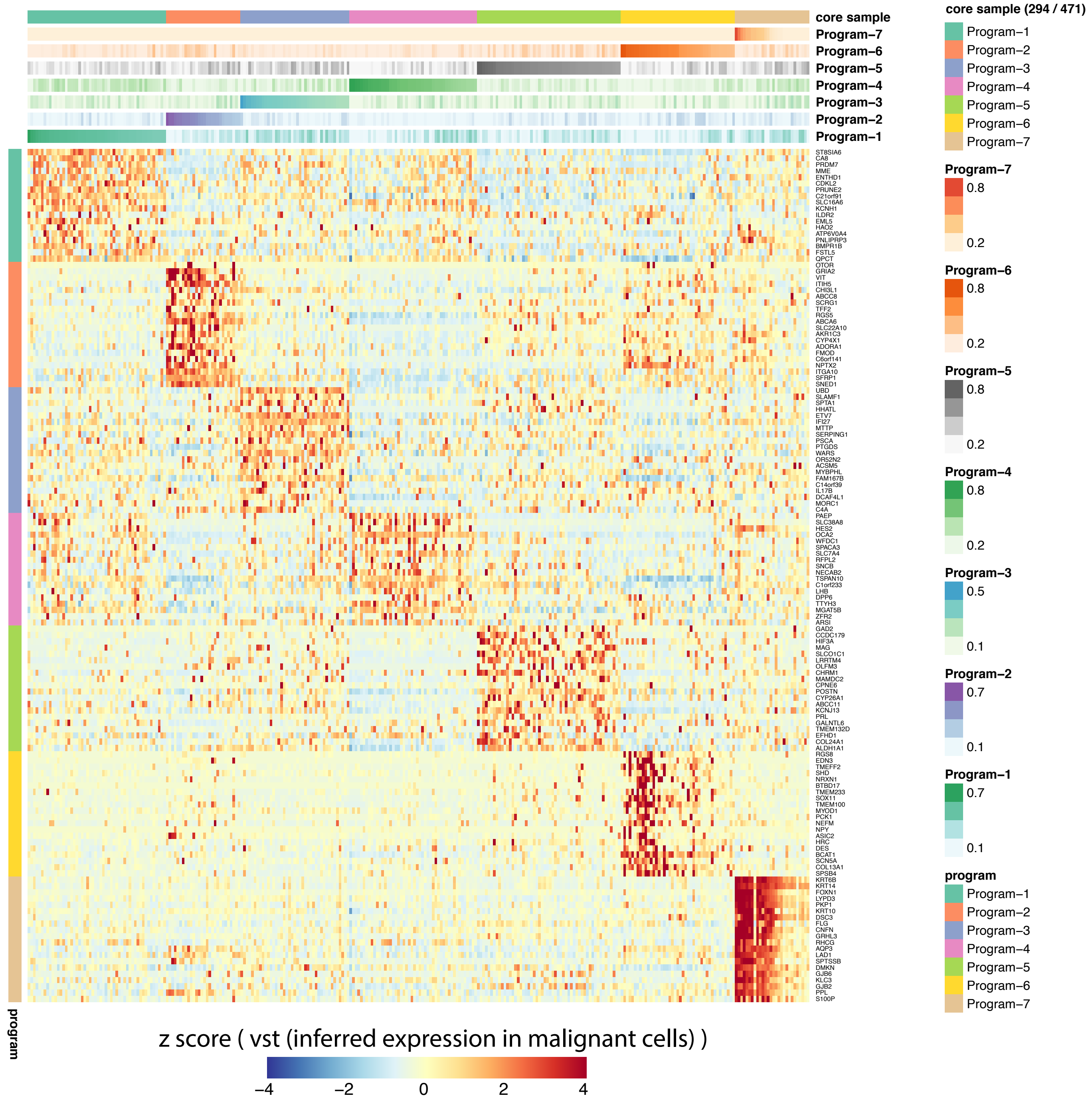


**Fig. S24 | Heatmap shows the relative expression of top 20 differentially expressed genes of each gene program across a set of TCGA bulk samples representing each gene program.** Each row represents a gene, while each column represents a TCGA bulk sample. The expression level was colored by the z score of variance-stabilizing transformed expression in malignant cells deconvolved by BayesPrism. Bulk samples were grouped by their affiliation to their gene program and ordered by normalized program weights (top annotation heatmaps). Genes were selected and ordered from high to low by the $\log_2$ fold change compared to samples affiliated to other gene programs.

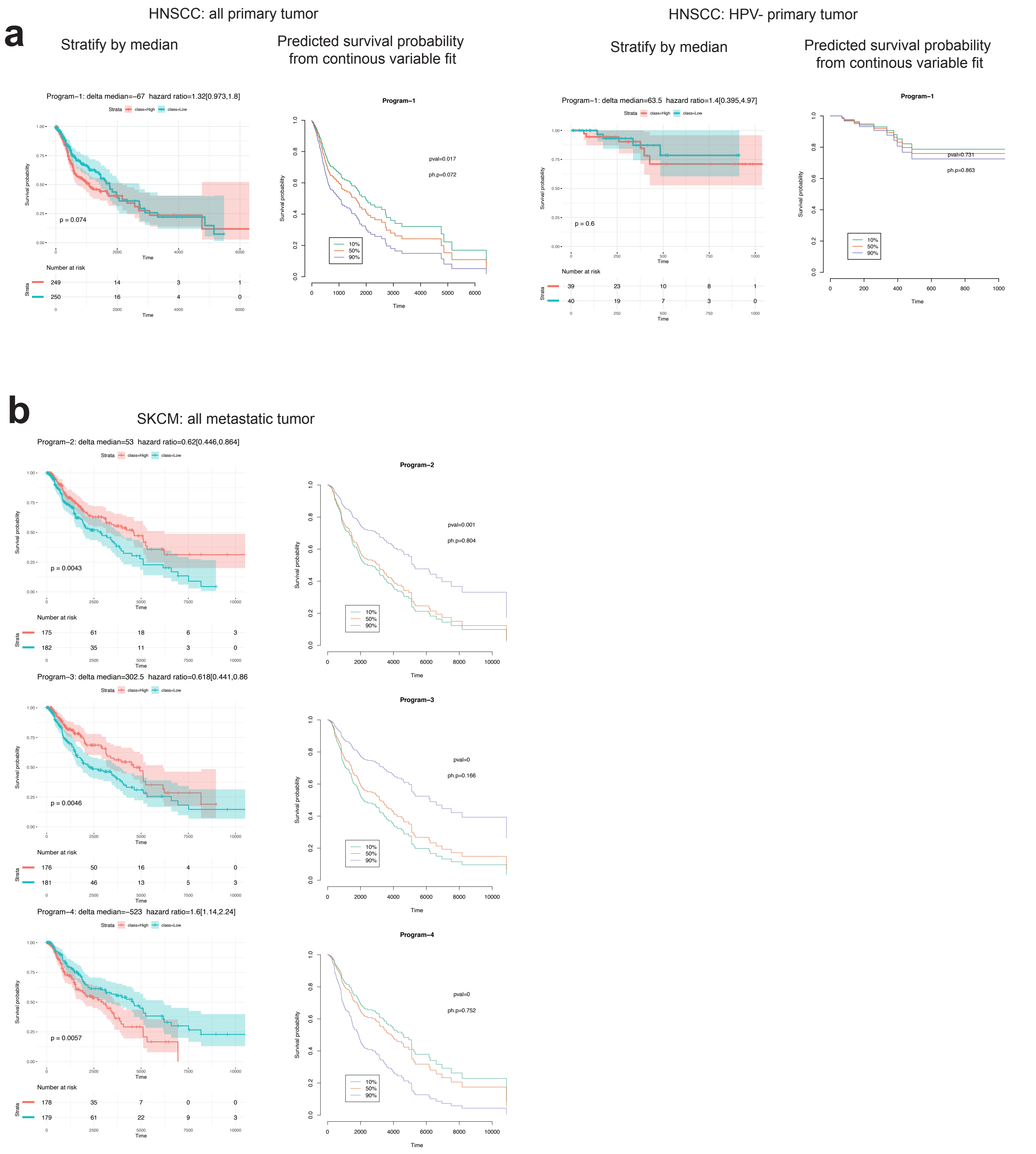


**Fig. S25 | KM plots of gene programs in malignant cells with significant association with survival by at least one model.** Two models of regression were shown: stratifying patients using the median cutoff, or treating the normalized program weights as continuous variables in the Cox proportional-hazards model. The log-rank test was used for the median-cutoff model. The survival curves in the right column were plotted by conditioning on the program weights at its 10%, 50% and 90% percentile, and then predicting survival probabilities based on the continuous variable Cox regression model, which was only used for the purpose of visualization. Two p values were computed for the continuous variable Cox regression model: “pval” was generated by the Wald test indicating the statistical significance of survival association, while “ph.p” was generated by the chi-squared test for scaled Schoenfeld residuals to check the proportional hazards assumption.
