## Supplementary Notes for "Bayesian cell-type deconvolution and gene expression inference reveals tumor-microenvironment interactions"

### Supplementary Note 1

#### Overview: BayesPrism (Bayesian) vs. frequentist regression-based approaches

Frequentist approaches, including most regressions and their regularized versions, assume the accuracy of reference expression matrix (under some error distribution) to derive a point estimate of cell type compositions. This assumption is often violated in the deconvolution of transcription profiles due to technical or biological variation between reference and bulk data of interest. This problem is particularly pronounced in cancer, where the transcription profile of malignant cells is highly heterogeneous across patients, a phenomenon largely attributed to the unique landscape of somatic mutations in individuals. Therefore, a reference expression matrix does not generalize to malignant cells in unobserved patients. In addition, batch effects between single cell reference and bulk tumor samples makes the reference matrix inaccurate.

To overcome these difficulties, BayesPrism uses φ, the reference expression matrix observed from scRNA-seq, as prior information which together with the prior on μ induces a prior distribution on the cell state-specific expression matrix U. The joint posterior distribution on μ and U is updated when X, the bulk RNA-seq information is observed. Note that the inference on U is not possible in frequentist approaches, where U either equals φ or not. By explicitly modeling the expression profile U in the bulk sample, BayesPrism accounts for the inaccuracy of the reference matrix φ.

#### 1. Gibbs sampling on cell state composition and gene expression

The inference of $P(\mu, U \mid \varphi, X; \alpha)$ in the first sampling of BayesPrism resembles sub-problems in the inference of latent Dirichlet allocation[1]. For readers who are familiar with work of topic models such latent Dirichlet allocation Blei et. al., each read in bulk RNA-seq is equivalent to a word; each bulk RNA-seq sample is equivalent to a document; each cell state is equivalent to a topic; each gene is equivalent to a vocabulary. The difference between BayesPrism and LDA is that in LDA the topic-word distribution, φ, is unknown and to be inferred from the posterior, while BayesPrism assumes it is known from the expression profile of scRNA-seq. The traditional sampling-based approach for LDA inference relies on the use of Gibbs sampling to approximate the posterior distribution of the topic distribution of each word[2], from which the topic-word distribution and document-topic distribution can be derived. This approach is computationally costly for inferring RNA-seq, where the word (read) number is typically at the scale of $10^8$ for one document (bulk RNA-seq sample). Besides, biological applications need not to know the posterior for each word, but are rather interested the expression level of each gene. Zhu and Lei et al. [3] recently improved the computational efficiency by sampling at the gene level through the introduction of augmented latent variables, such that the time complexity is only a function of number of genes, which are usually between 10K-50K, and is independent of sequencing depth. We follow their derivation to get the formula for the first Gibbs sampling of BayesPrism's, and provide a more detailed step-by-step derivation, and adding back missing terms in their likelihood function.

The full model specification is as follows. Each bulk sample $n \in \{1, \ldots, N\}$ is measured across G annotated genes, and the bulk RNA-seq data is represented by a matrix $X \in \mathbb{R}^{N \times G}$. As the genes that each read aligned to is also observed, each row of X, denoted by $X_n$, can be expanded to $\tilde{X}_{n,r} \in \mathbb{R}^G$ to denote the gene that the r^th^ read in the n^th^ sample aligned to. We assume that the cell states of each cell of the scRNA-seq dataset are known. The reference expression profiles of a total S cell states estimated from the scRNA-seq data denoted by $\varphi \in \mathbb{R}^{S \times G}$. We model the distribution of read counts of scRNA-seq in each cell state using multinomial distribution, and hence each row of $\varphi_s \in \mathbb{R}^G$ is an maximum likelihood estimator of the multinomial distribution event probability parameters, such that $\sum_{g=1}^{G} \varphi_{s,g} = 1$, for $\forall s \in \{1, \dots, S\}$. To avoid having zeros in φ, for each $\varphi_s$ we compute the pseudo count such that the $\min(\varphi_{s,g}) = 10^{-8}$ after renormalizing to one. $\mu \in \mathbb{R}^{N \times S}$ denotes the fraction of reads assigned to the $s^{th}$ cell state in the $n^{th}$ bulk RNA-seq sample. $Y_{n,r}$ denotes the latent variable describing the cell state that the $r^{th}$ read of the $n^{th}$ sample belongs to. α is the hyper-parameter of the Dirichlet prior on μ. We set $\alpha = 10^{-8}$ to represent a weak and non-informative prior, such that the posterior is mainly driven by the likelihood. The augmented cell state expression tensor $U \in \mathbb{R}^{N \times S \times G}$ is defined as $U_{n,s,g} = \sum_{\{r:X_{r,n}=g\}} I_{\{Y_{n,r}=s\}}$, which denotes the number of reads assigned to the $g^{th}$ gene in $s^{th}$ cell state of the $n^{th}$ bulk sample.

The generative process is described as follows.

1. Generate fractions for tumor and environmental cells:
   $\mu_n \sim$ Dirichlet (α), i.i.d. for $n \in \{1, \dots, N\}$, and $\alpha > 0$.

2. Generate the reads for bulk RNA-seq
   $Y_{n,r} \mid \mu_n \sim$ Categorical ($\mu_n$), independently for $n \in \{1, \dots, N\}$, and $r \in \{1, \dots, R_n\}$
   $\tilde{X}_{n,r} \mid Y_{n,r}, \varphi \sim$ Categorical ($\varphi_{Y_{n,r}}$), i.i.d. for $r \in \{1, \dots, R_n\}$
   $X_{n,g} = \sum_{r=1}^{R_n} I_{\{\tilde{X}_{n,r}=g\}}$
   $U_{n,s,g} = \sum_{\{r:X_{r,n}=g\}} I_{\{Y_{n,r}=s\}}$

   *Note that the distribution is written as multinomial distribution in Methods, which is an equivalent but more compact representation of the categorical distribution. We expand it here using the categorical distribution to facilitate understanding and downstream derivation.

To derive the posterior of P(μ, U | φ, X; α), we first write down the complete likelihood function:

$$p(X, Y, U, \mu \mid \varphi; \alpha) = p(\mu \mid \alpha)\, p(Y \mid \mu)\, p\,(\tilde{X} \mid Y)\, p(X \mid \tilde{X})$$

$$\propto \prod_{n=1}^{N} \left\{ \begin{array}{c} \Gamma(S \cdot \alpha) \prod_{s=1}^{S} \frac{\mu_{n,s}{}^{(\alpha-1)}}{\Gamma(\alpha)} \cdot \prod_{r=1}^{R_n} \prod_{s=1}^{S} \mu_{n,s}{}^{I_{\{Y_{n,r}=s\}}} \cdot \\ \prod_{s=1}^{S} \prod_{g=1}^{G} \varphi_{s,g}{}^{\sum_{r:\tilde{X}_{n,r}=g} I_{\{Y_{n,r}=s\}}} \cdot \prod_{g=1}^{G} I_{\{X_{n,g}=\sum_{r=1}^{R_n} I_{\{\tilde{X}_{n,r}=g\}}\}} \end{array} \right\} \quad (1)$$

Further observe that

$$\prod_{r=1}^{R_n}\prod_{s=1}^{S}\mu_{n,s}{}^{I_{\{Y_{n,r}=s\}}}$$
$$=\prod_{s=1}^{S}\mu_{n,s}{}^{\sum_{r=1}^{R_n} I_{\{Y_{n,r}=s\}}}$$
$$=\prod_{s=1}^{S}\mu_{n,s}{}^{\sum_{g=1}^{G}\sum_{r=1}^{R_n} I_{\{Y_{n,r}=s\ \&\ \tilde{X}_{n,r}=g\}}}$$
$$=\prod_{s=1}^{S}\mu_{n,s}{}^{\sum_{g=1}^{G} U_{n,s,g}}$$
$$=\prod_{g=1}^{G}\prod_{s=1}^{S}\mu_{n,s}{}^{U_{n,s,g}} \quad (2)$$

$$\prod_{s=1}^{S}\prod_{g=1}^{G}\varphi_{s,g}{}^{\sum_{r:\tilde{X}_{n,r}=g} I_{\{Y_{n,r}=s\}}}$$
$$=\prod_{g=1}^{G}\prod_{s=1}^{S}\varphi_{s,g}{}^{U_{n,s,g}} \quad (3)$$

and,

$$I_{\{X_{n,g}=\sum_{r=1}^{R_n} I_{\{\tilde{X}_{n,r}=g\}}\}}$$
$$=I_{\{X_{n,g}=\sum_{r=1}^{R_n}\sum_{s=1}^{S} I_{\{\tilde{X}_{n,r}=g\ \&\ Y_{n,r}=s\}}\}}$$
$$=I_{\{X_{n,g}=\sum_{s=1}^{S} U_{n,s,g}\}} \quad (4)$$

Combining 1-4, get

$$(1)=\prod_{n=1}^{N}\left\{\begin{matrix}\Gamma(S\cdot\alpha)\ \prod_{s=1}^{S}\frac{\mu_{n,s}{}^{(\alpha-1)}}{\Gamma(\alpha)}\cdot\prod_{g=1}^{G}\prod_{s=1}^{S}(\mu_{n,s}\varphi_{s,g})^{U_{n,s,g}}\cdot \\ \prod_{g=1}^{G} I_{\{X_{n,g}=\sum_{s=1}^{S} U_{n,s,g}\}}\end{matrix}\right\}$$

Therefore,

$$p(\mu_{n,\cdot}\mid X,U,\varphi;\alpha)\ \propto\ \prod_{s=1}^{S}\mu_{n,s}{}^{(\sum_{g=1}^{G} U_{n,s,g}+\alpha-1)},$$
$$p(U_{n,\cdot,g}\mid \mu,X,\varphi;\alpha)\ \propto\ \prod_{s=1}^{S}(\mu_{n,s}\varphi_{s,g})^{U_{n,s,g}}\ I_{\{X_{n,g}=\sum_{s=1}^{S} U_{n,s,g}\}} \quad (5)$$

Their corresponding distributions can then be read off from (5):

$$\mu_{n,\cdot}\mid X,U,\varphi;\alpha\sim Dirichlet\ (\alpha+\ \textstyle\sum_{g=1}^{G} U_{n,\cdot,g}\ ),$$
$$U_{n,\cdot,g}\mid \mu,X,\varphi;\alpha\sim Multinomial\ (\frac{\mu_{n,\cdot}\odot\varphi_{\cdot,g}}{\sum_{s=1}^{S}\theta_{n,s}\varphi_{s,g}},X_{n,g}),$$

where $\odot$ is element-wise multiplication. (6)

We initiate $\mu$ to a value of $1/S$. As each sample is conditionally independent, they can be sampled in parallel. Empirically, the Gibbs chain converges fairly fast. This is mainly due to the high read depth of bulk RNA-seq samples, which gives a fairly concentrated posterior distribution on $\mu$. The default setting for Gibbs sampling is as follows: length of chain = 1000; burn in = first 500; thinning =2. BayesPrism reports the mean of posterior, E[U] and E[$\mu$]. In practice, the Gibbs chain converges very fast due to the large number of reads in the bulk (Fig. SN1):

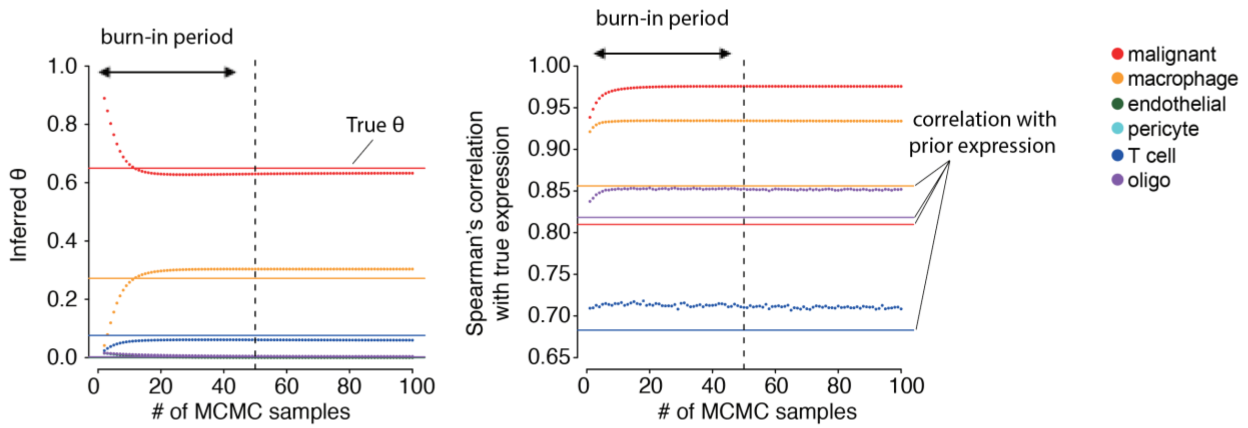


**Fig. SN1. MCMC diagnostic plots for cell type fractions (left) and Spearman's correlations between the inferred gene expression and true gene expression.** X axis marks the number of MCMC samples, i.e. chain length. The horizontal lines in the **left** panel mark the true cell type fractions, while those on the **right** panel mark the Spearman's correlations between the gene expression of the scRNA-seq prior and the true gene expression. The posterior quickly reaches a concentrated and unimodal stationary distribution after around 20 MCMC cycles.

#### 2. update the reference matrix $\psi$ for malignant and non-malignant cells

To estimate the malignant expression reference in each patient, we model the likelihood using multinomial distribution with the parameter $\psi_{mal_n} \in \mathbb{R}^G$, and get the maximum likelihood estimator:

$$\underset{\psi_{mal_n}}{\operatorname{argmax}} \; p\left(Z_{n,t,\cdot} \mid \psi_{mal_n}\right) = \frac{Z_{n,t,g}}{\sum_{g=1}^{G} Z_{n,t,g}} \;, where\ t = malignant \tag{7}$$

We also use the multinomial distribution to model the likelihood function for the expression profiles of non-malignant cells shared across all patients. In this step we pooled the observations of Z across all bulk RNA-seq samples for each non-malignant cell to borrow the statistical strength across samples. To prevent pathological cases where subsets of non-malignant cells have a close to zero fraction in all bulk RNA-seq, in which case the Z will be close to zero and hence dominated by the sampling noise, we put a zero-mean log normal-distributed prior on the fold change with respect to their corresponding reference φ' (the scRNA-seq reference matrix defined over cell types, similar to φ), and use the maximum a posteriori (MAP) estimator for $\psi_{env}$

$$D_t = log\left( p\left( \psi_{env_t} \middle| Z_{\cdot,t,\cdot} \,, \varphi'_t ; \sigma\right)\right) \tag{8}$$

$$= \sum_{n=1}^{N} log\left(p\left( Z_{n,t,\cdot} \mid \psi_{env_t}\right)\right) + log\left( p\left(\psi_{env_t} \mid \varphi'_t ; \sigma \right)\right) + C$$

$$= \sum_{n=1}^{N} \sum_{g=1}^{G} Z_{n,t,g} \log\left(\psi_{env_{t,g}}\right) - \frac{1}{2\sigma^2} \sum_{g=1}^{G} \log\left(\frac{\psi_{env_{t,g}}}{\varphi'_{t,g}}\right)^2 + C', where\ t \in \{malignant\}^c$$

The MAP estimator of $D_t$ has no closed form solution and needs to be optimized numerically. Directly optimizing over $\psi$ is more difficult, due to the constraint that $\sum_{g=1}^{G} \psi_{env_{t,g}} = 1$. We therefore used the change of variables method, by letting $\gamma_{t,g} = \log\left(\frac{\psi_{str_{t,g}}}{\varphi'_{t,g}}\right)$, and $\psi_{env_{t,g}} = \frac{\varphi'_{t,g} \cdot \gamma_{t,g}}{\sum_{g=1}^{G} \varphi'_{t,g} \cdot \gamma_{t,g}}$, to make the optimization unconstrained and numerically more stable (see below).

$$D_t = \sum_{g=1}^{G} \left(\sum_{n=1}^{N} Z_{n,t,g}\right) \log\left(\frac{\varphi_{env_{m,g}} \cdot \gamma_{m,g}}{\sum_{g=1}^{G} \varphi_{env_{m,g}} \cdot \gamma_{m,g}}\right) - \frac{1}{2\sigma^2} \sum_{g=1}^{G} \gamma_{t,g}{}^2 + C' \quad (9)$$

$\sum_{n=1}^{N} Z_{n,t,g}$ can be computed before optimization, and let $Z_{t,g} = \sum_{n=1}^{N} Z_{n,t,g}$. With some algebra, the partial derivative of the posterior can be derived as

$$\frac{\partial D_t}{\partial \gamma_{t,g}} = Z_{t,g} - \psi_{env_{t,g}} \left(\sum_{g=1}^{G} Z_{t,g}\right) - \frac{1}{\sigma^2} \gamma_{t,g} \quad (10)$$

As $\frac{\partial D_t}{\partial \gamma_{t,g}}$ is a function of only $\gamma_t$, they can be optimized in parallel. We use the conjugate gradient method written by the Rcgmin package[4] with minor modifications by adding a stopping criterion to prevent it making more than 500 consecutive non-productive iterations. All $\gamma$ are initiated at 0. By default, we set $\sigma = 2$, for all genes, which is around the typical range of log fold change between two batches of RNA-seq dataset and is a very weak prior compared to the likelihood. Users may also supply their own σ based on their prior knowledge, for example the standard deviation of log fold change from differential expression analysis between a pair of matched samples, or the closest scRNA-seq reference and bulk RNA-seq samples.

#### 3. Learning latent embeddings for malignant gene programs.

##### 3.1 Model specification

The goal of learning the latent embeddings is to approximate the expression of malignant cells across a cohort of N bulk RNA-seq samples as a linear combination of a small number of K bases, with K<<N. Complete factorization approaches, such as NMF and LDA, aim at factorizing the bulk RNA-seq down to a linear combination of K bases (gene programs), with K<<N. However, gene programs inferred by these approaches will be confounded by the expression of non-malignant cells. Additionally, expression in malignant cells is highly heterogenous (i.e., in practice, K=N for this problem), and hence reducing K to a number significantly less than N is a lossy compression. To overcome these limitations, the strategy used by BayesPrism differs in that it conditions on fraction and expression profile of non-malignant cells, i.e. $\theta_{env}$ and $\psi_{env}$ , inferred by the deconvolution module, and learns the embeddings of malignant cells.

We denote the initial K tumor basis as $\eta_0 \in \mathbb{R}^{K \times G}$, and a set of perturbations from $\eta_0$ to be inferred as $\lambda \in \mathbb{R}^{K \times G}$, their associated weights as $\omega \in \mathbb{R}^{N \times K}$ . The total number of components in the embedding learning is then $M = K + T - 1$ .

The probabilistic graphical model is shown by **Fig. SN2** below:

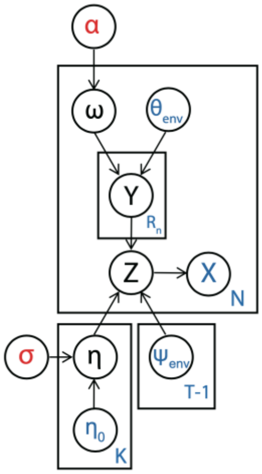


**Fig. SN2. Graphical model illustrates the statistical dependencies and the generative process for the observed bulk RNA-seq data, X.** Red text marks hyper-parameters; blue marks observed variables; black marks latent variables.

The generative process is as follows.

1. Generate weights for tumor basis:

   $\kappa_n \sim$ Dirichlet ($\alpha$), i.i.d. for n∈{1, ..., N}, where $\alpha \in \mathbb{R}^K$ and $\alpha > 0$.

   $\omega_n = \tau_n \kappa_n$, where $\tau_n = 1 - \sum_{t\in\{malignant\}^c} \theta_{n,t}$

   Concatenating the columns of $\omega$ and $\theta_{env}$ , we get $\nu \in \mathbb{R}^{N\times M}$ , where $\sum_{m=1}^{M} \nu_{n,m} = 1$ .

   *Note that $\alpha$ need not be the same as the one in equation (1). By default, however, BayesPrism uses the same $\alpha$ for a non-informative sparse prior.

2. Generate the full expression profiles:

   log ($\lambda_{k,g}$) ~ Normal (0, σ), for g∈{1, ..., G} and k ∈ {1, ..., K}

   *Note that σ need not be the same as the one in equation (8). By default, however, BayesPrism uses the same σ.

   $\eta_{k,g} = \frac{\eta_{0_{k,g}} \cdot \lambda_{k,g}}{\sum_{g=1}^{G} \eta_{0_{k,g}} \cdot \lambda_{k,g}}$ for k ∈ {1, … , K }

   Concatenating the rows of η and $\psi_{env}$, we get the total expression profile $\zeta \in \mathbb{R}^{M\times G}$

3. Generate the reads for bulk RNA-seq:

   $Y_{n,r} \mid \nu_n \sim$ Categorical ($\nu_n$), independently for n∈{1, ..., N}, and r∈{1, ..., $R_n$ }

   $\tilde{X}_{n,r} \mid Y_{n,r}, \zeta \sim$ Categorical ($\zeta_{Y_{n,r}}$ ), i.i.d. for r ∈{1, ..., $R_n$}

   $X_{n,g} = \sum_{r=1}^{R_n} I_{\{\tilde{X}_{n,r}=g\}}$

   $V_{n,m,g} = \sum_{\{r:X_{r,n}=g\}} I_{\{Y_{n,r}=m\}}$

#### 2.2 Model inference using Expectation-maximization (EM)

As ω, η, and Z are all latent variables and are marginally dependent on each other, direct inference can be difficult. Considering the advantage of marginalizing nuance variables[5], we use the Expectation-maximization (EM) algorithm to optimize η while marginalizing $\omega$ and V.

The EM algorithm is formulated as follows. In the M step the posterior we would like to maximize $E_Q[\log(p(\lambda,\ V \mid X, \eta_0,\ \psi_{env}, \theta_{env};\ \sigma))]$, with the expectation taken over $Q = p(\ V \mid \psi_{env}, \theta_{env}, \eta^{old};\ \sigma, \alpha)$, i.e. the posterior sampled by Gibbs sampling in the E step, which constitutes the Gibbs-EM algorithm[6].

##### 2.2.1 The E step

The complete likelihood function for the E step is

$$p(\omega, Y, \tilde{X}, X \mid \eta^{old}, \psi_{env}, \theta_{env};\ \sigma, \alpha) = p(\omega \mid \tau;\alpha)\, p(Y \mid \nu, \zeta)\, p(\tilde{X} \mid Y)\, p(X \mid \tilde{X})$$
$$= \prod_{n=1}^{N}\{p(\omega_n \mid \tau_n;\alpha) \prod_{r=1}^{R_n} p(Y_r \mid \nu,\ \zeta)\, p(\tilde{X}_r \mid Y_r)\, p(X_r \mid \tilde{X}_r)\}$$

Observe that the posterior of $p(V_{n,\cdot,g} \mid \nu, X, \zeta; \alpha)$ follows the same form as the $p(U_{n,\cdot,g} \mid \mu, X, \varphi; \alpha)$ in (5). Hence we have:

$$p(V_{n,\cdot,g} \mid \nu, X, \zeta; \alpha) \sim Multinomial\ (\frac{\nu_{n,\cdot} \odot \zeta_{\cdot,g}}{\sum_{m=1}^{M} \nu_{n,m}\zeta_{m,g}}, X_{n,g}),$$

where ⊙ is element-wise multiplication.

Sampling from $p(\omega_{n,\cdot} \mid X, V, \zeta; \alpha)$ is slightly different from (5), due to the scaling factor $\tau_n$. Since $\omega$ is deterministic of κ, sampling $\omega$ is trivial when one can sample from the posterior of κ, i.e. $p(\kappa_{n,\cdot} \mid X, V, \zeta; \alpha)$, which we derive as follows:

$$p(\kappa_{n,\cdot};\ \alpha) \propto \prod_{k=1}^{K} \kappa_{n,k}^{\alpha-1}$$

$$p(L_{n,\cdot} \mid \kappa_{n,\cdot}, \theta_{n,\cdot}) \propto \prod_{k=1}^{K} (\tau_n \kappa_{n,k})^{L_{n,k}} \prod_{t\in\{malignant\}^c} \theta_{n,t}^{O_{n,t}} = \prod_{k=1}^{K} \kappa_{n,k}^{L_{n,k}} \cdot C$$

, where $L_{n,k}=\sum_{g=1}^{G} V_{n,k,g}$, for k∈{1,…,K}, and $O_{n,t}=\sum_{g=1}^{G} V_{n,t,g}$, for t∈{K+1,…,T-1}

Therefore, $p(\kappa_{n,\cdot} \mid X, V, \zeta; \alpha) \propto p(\kappa_{n,\cdot};\ \alpha) \cdot p(L_{n,\cdot} \mid \kappa_{n,\cdot}, \theta_{n,\cdot}) \propto \prod_{k=1}^{K} \kappa_{n,k}^{L_{n,k}+\alpha-1}$.

The conditional distribution follows $\kappa_{n,\cdot} \mid X, V, \zeta; \alpha \sim Dirichlet\ (\alpha +\ L_{n,\cdot})$.

##### 2.2.1 The M step

In the M step, we aim to find $\underset{\eta}{\text{argmax}}\ E_Q[\log(\ p(\eta, V\ |X, \eta_0, \psi_{env}\ , \theta_{env};\ \sigma))]$, with Q being the Gibbs samples describing the posterior of V sampled at the E step. $\underset{\eta}{\text{argmax}}\ E_Q[\log(\ p(\eta, V, |X, \eta_0, \psi_{env}\ , \theta_{env};\ \sigma, \alpha))]$ is a constrained optimization, with $\sum_{g=1}^{G} \eta_{k,g} = 1$ for all ∀ k∈ {1, … , K }. We again turn this into an un-constrained optimization by optimizing its equivalent form $\underset{\lambda}{\text{argmax}}\ E_Q[\log(\ p(\lambda, V, |X, \eta_0,\ \psi_{env}\ , \theta_{env};\ \sigma))]$.

The complete log posterior written in the augmented form is

$$
\begin{aligned}
& E_Q[log(p(\ \lambda, V\ |X, \eta_0, \psi_{env}, \theta_{env};\ \sigma))] \\
&= E_Q[log(p(\ V, X\ |\ \lambda, \eta_0, \psi_{env}, \theta_{env}))]\ + log(p(\lambda;\ \sigma))\ + C \\
&= E_Q[log(p(\ V\ |\ \lambda, \eta_0, \psi_{env}, \theta_{env}))] + log(p(\lambda;\ \sigma))\ + C \\
&= \sum_{g=1}^{G} \sum_{k=1}^{K} E_Q[log(p(\ V_{n,k,g} |\ \lambda_{m,g}, \eta_{0_{m,g}}))]\ \} + \sum_{g=1}^{G} \sum_{k=1}^{K} log(p(\lambda_{k,g};\ \sigma))\ + C' \\
&= \sum_{n=1}^{N} \sum_{g=1}^{G} \sum_{k=1}^{K} E_Q\left[V_{n,k,g} \log\left(\zeta_{k,g}\right)\right] + \sum_{g=1}^{G} \sum_{k=1}^{K} \log\left(p\left(\lambda_{k,g};\ \sigma\right)\right)\ + C''
\end{aligned}
$$

The constants C, C' and C" are with respect to the parameter λ.

Similar to the derivation of (9) and (10), the posterior and its derivative are

$$
D_k = \sum_{g=1}^{G} \left(\sum_{n=1}^{N} E_Q[V_{n,k,g}]\right) \log\left(\frac{\eta_{0_{k,g}} \cdot \lambda_{k,g}}{\sum_{g=1}^{G} \eta_{0_{k,g}} \cdot \lambda_{k,g}}\right) - \frac{1}{2\sigma^2} \sum_{g=1}^{G} {\gamma_{k,g}}^2 \tag{12}
$$

Since $\sum_{n=1}^{N} E_Q[V_{n,k,g}]$ can be computed before optimization, and we can let $V_{k,g} = \sum_{n=1}^{N} E_Q[V_{n,k,g}]$, with some algebra, the partial derivative of the posterior can be derived as

$$
\frac{\partial D_k}{\partial \gamma_{k,g}} = V_{k,g} - \eta_{k,g}\left(\sum_{g=1}^{G} V_{k,g}\right) - \frac{1}{\sigma^2} \gamma_{k,g} \tag{13}
$$

As $\frac{\partial D_k}{\partial \gamma_{k,g}}$ is a function of only $\gamma_k$, they can be optimized in parallel for each k. The optimization was implemented by the conjugate gradient method in the same way as for $\psi_{env}$. σ=2 is a very weak prior compared to the likelihood, and mainly serves as a way to prevent zero probability in $\eta$ during EM. There are several ways to set $\eta_0$ and K. $\eta_0$ can be set identically for all K by taking an average of the tumor expression profiles, $\psi_{mal}$, estimated in equation (7). Despite being initialized equally, $\gamma_k$will eventually diverge and pick up a gradient after 10-20 EM cycles, due to the randomness of Gibbs-sampling. In practice this approach takes more EM cycles to converge. Alternatively, user may use the NMF to factorize $\psi_{mal}$ , and pick the optimum K based on criteria used for consensus clustering, e.g. cophenetic correlation, dispersion, silhouette score, etc [7]. This approach converges much faster than a uniform initialization, and typically converges within 50 cycles (**Fig. SN3**). Biological prior knowledge can also be used to initiate $\eta_0$ using the expression of known subtypes or origin cells of malignant cells.

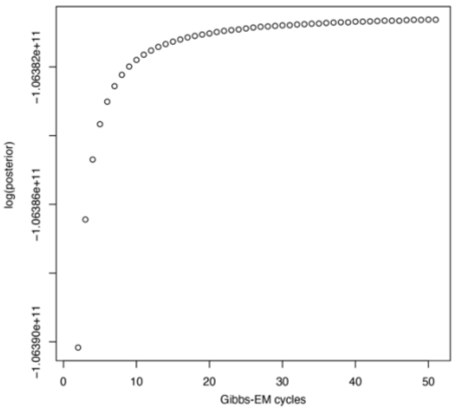


**Fig.SN3. log(posterior) of the model as a function of EM cycles.**

#### Supplementary Note 2: Benchmarking computational efficiency

As CIBERSORTx is based on SVR which scales cubically ($O(N^3)$) as the number of genes, and does not parallelize for each bulk sample, it does not scale to datasets with large numbers of marker genes. In contrast, BayesPrism scales linearly ($O(N)$), and takes advantage of parallelization (see figure below). To make the benchmark reasonably computable, we downsampled the ⅕ of the 1,350 pseudo-bulks for comparison in **Fig.1c-d** and **Supplementary Fig. 6**.

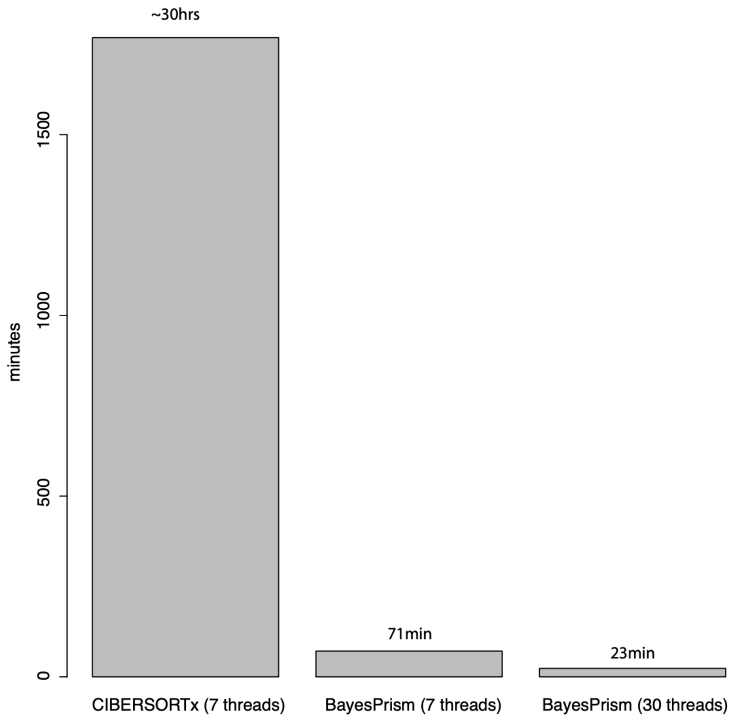


**The estimated computing time of the online version of CIBERSORTx and BayesPrism.** The computing time of BayesPrism was tested on Intel(R) Xeon(R) CPU E5-4620 v2 @ 2.60GHz.

#### Supplementary Note 3: Mathematical intuition of the robustness of BayesPrism

In this supplementary note, we discuss the intuition of the robustness of BayesPrism by demonstrating that BayesPrism is mathematically invariant to two types of noise that represent idealized cases of technical batch effects and biological variation. The proof of invariance follows from the formula used to perform Gibbs sampling, as derived in the equation (6) in the **Supplementary Note 1**, and copied below.

$$\mu_{n,\cdot} \mid X, U, \varphi; \alpha \sim Dirichlet\ \left(\alpha + \sum_{g=1}^{G} U_{n,\cdot,g}\right),$$

$$U_{n,\cdot,g} \mid \mu, X, \varphi; \alpha \sim Multinomial\ \left(\frac{\mu_{n,\cdot} \odot \varphi_{\cdot,g}}{\sum_{s=1}^{S} \theta_{n,s}\varphi_{s,g}}, X_{n,g}\right),$$

where $\odot$ is element-wise multiplication. (6)

The first type of noise is linear multiplicative noise, where the true reference in the bulk RNA-seq differs from the observed reference in scRNA-seq by a multiplicative constant:

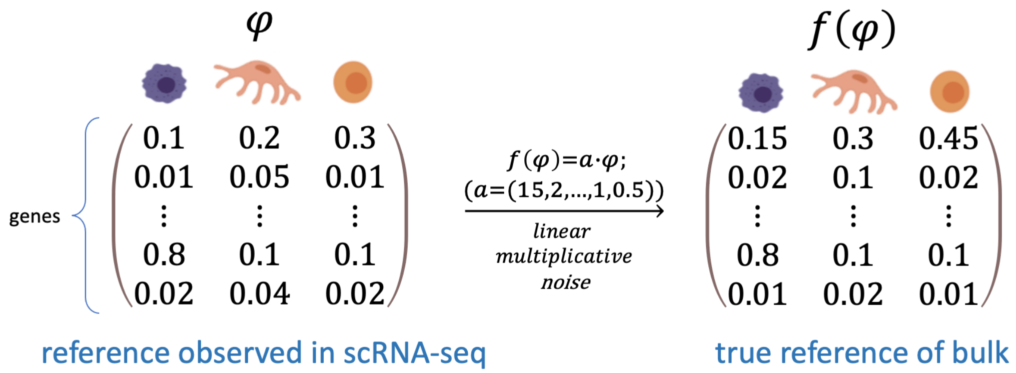


Notice that under such noise, the relative ratios of the expression of each gene between cell types from the observed reference are the same as the true reference, and hence this noise mimics the technical batch effects where genes are captured at different efficiencies by two experimental / sequencing methods. By replacing $\varphi$ with $a \cdot \varphi$ in (6), it is obvious that the posterior of U remains the same, as the constant cancels out. Therefore, BayesPrism is invariant to such noise. **Supplementary Fig. 2** simulates the performance of BayesPrism and other deconvolution tools under such a noise model.

The second type of noise is any arbitrary transformation on marker genes expressed exclusively in one cell type, provided that the expression in other cells remains zero after the transformation:

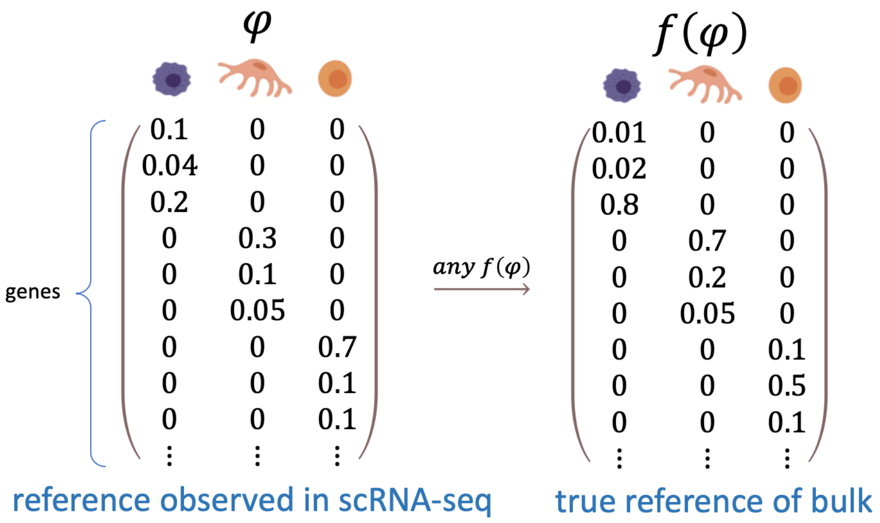


Under such noise, it is also obvious that the posterior of U remains the same, and reads from the marker genes of a particular cell type will be exclusively assigned to that cell type, resulting in a Dirac posterior, and hence a rapid convergence of the Gibbs chain in one sampling step. This noise model mimics highly cell type specific genes that retain their cell type specificity under both technical batch effects and biological variation. It is likely that the combination of the invariance to the two types of noise contributes to the robustness of BayesPrism.

#### Supplementary Note 4: Robustness against an undersampled reference

In this supplementary note, we discuss the details of how an undersampled reference can affect the accuracy of deconvolution. An undersampled reference falls into one of the following three categories: 1) reference with one or few cell types missing, 2) reference with low numbers of cells, 3) reference with limited number of patients (in the tumor setting). We discuss how each of them may affect the accuracy of deconvolution.

Some cell types can be missing from the single cell reference due to the scarcity or low capture efficiency, hereafter referred to as “missing cell types”. In the absence of tumor heterogeneity, the expression of all cell types or a subset of missing cell types, can be recovered using matrix factorization-based approaches, such as NMF and LDA, given a large amount of bulk RNA-seq samples each containing different fractions of cell types and the presence of signature genes. In the presence of heterogeneous tumors, the *de novo* modeling of the expression of missing cell types becomes an unidentifiable problem. In practice, the expression of missing cells will be redistributed into the reference cells by all deconvolution methods, according to the similarity with respect to each reference cell type. We therefore empirically tested how missing cell types may possibly affect BayesPrism by excluding T cell, a rare cell type in GBM, from the refGBM8 reference, and tested the performance in deconvolving the GBM28 pseudo-bulks (N=1,350) containing T cells. We found that the cell type-level correlations are robust up to 40-50% of missing T cells (**Supplementary Fig. 7, left**). The MSE, however, goes up quickly as the missing fraction, indicating that the missing cell fraction becomes redistributed into other cell types (**Supplementary Fig. 7, right**). Macrophages are the most affected, due to its higher similarity in expression with T cells. Nevertheless, this is less problematic for references of missing cell types due to scarcity in the tissue. For instance, T cells are present at < 5% in both GBM scRNA-seq datasets, at which level the deconvolution is not affected. Subsetting on genes that are not (or less) expressed in the missing cell types can also ameliorate this issue.

Number of cells in the scRNA-seq reference also affects the deconvolution. The extent to which this affects accuracy depends on the cell types to be inferred and the sequencing depth of the single cell platforms. As a rule of thumb, to resolve highly similar cell types, e.g. subtypes of T cells, larger numbers of cells (and hence more information) are needed. The higher the heterogeneity the tumor is, the more patients are needed. The shallower the sequencing depth, e.g. 10X vs SMART-seq2, the greater the numbers of cells are needed. It is impossible to determine the minimum number of cells for each task a priori. Therefore, we recommend to run a leave-one-out test to estimate the actual performance. To estimate the ballpark, we downsampled the cells in each cell type to a particular size N (without replacement). As the number of cells in the scRNA-seq reference vary between cell types, we are only allowed to downsample cell types with a higher number of cells than N. Therefore, N represents the maximum number of cells of all cell types. BayesPrism approaches its optimal performance when the maximum number of cell types contain between 100 (for SMART-seq2) to 500 cells (for platforms with depth close to 10X) (**Supplementary Fig. 8b-e**). Taken together, our analyses suggest that BayesPrism is generally robust to reference with undersampled cell numbers.

In the case of tumor deconvolution, the number of patients / number of tumor sub-clusters in the reference may also affect the accuracy. The extent to which this affects accuracy depends on the heterogeneity of the tumor of interest. We recommend planning the experiments by running a leave-one-out test to estimate the number of tumor patients needed. It is often ideal to have a scRNA-seq reference that represents all major tumor cell states or subtypes. To test empirically, we subsampled the malignant cells from individual patients from the 8 GBM scRNA-seq reference dataset. BayesPrism achieved saturating levels of performance deconvolving GBM28 pseudo-bulk data with as few as 3-4 patients (**Supplementary Fig. 8a**). Taken together, the analysis suggests that BayesPrism can account for tumor heterogeneity with relatively small numbers of cancer patients.

#### Supplementary Note 5: Effects of ribosomal protein coding genes and mitochondrial genes on deconvolution

When benchmarking deconvolution for several of the datasets under a marker-free mode, we found that the performance of all methods including BayesPrism improved when mitochondrial and ribosomal protein coding genes were excluded. These housekeeping genes were ubiquitously transcribed in all reference scRNA-seq datasets. The expression level of ribosomal protein coding genes ranges from 12% to 38% of the total mapped reads (human PBMC data[42]). They are often not informative in distinguishing cell types and can be a source of large spurious variance[43]. Cell types that were transcriptionally similar, such as CD4+ and CD8+ T cells, were more sensitive to these genes, as fewer genes were informative in distinguishing them. The accuracy of devonovolution is less sensitive to the expression level of mitochondrial genes, as their expression level is lower than that of ribosomal protein coding genes, ranging from 3% to 13% of total mapped reads (human PBMC data[42]). However, considering the level of mitochondrial genes often reflect technical bias of the scRNA-seq experiments, such as the survival of cells, we also recommend excluding them from the scRNA-seq reference.

Additionally, as BayesPrism estimates the fraction of reads from each cell type, and malignant cells may systematically upregulate the ribosomal protein coding genes[44], the inclusion of these genes may result in an overestimation of the fraction of malignant cells by a multiplicative constant.

In addition to ribosomal genes, we also advise users to look out for highly expressed genes, such as MALAT1 and other nuclear lincRNAs, actin and hemoglobin, as recommended by some scRNA-seq processing pipelines. As minor differences in the expression level of these genes may cause global changes in normalization when constructing the reference matrix.

#### Supplementary Note 6: Rich correlation structure between stromal cells

To determine how non-malignant cell types co-varied with each other, we examined the pairwise correlations between each cell type in the TCGA cohort. In GBM, pericytes and endothelial cells showed strong correlation (Spearman's rank correlation [ρ] = 0.34; **Supplementary Fig. 16a**), consistent with their combined presence in vascular structures[7]. Macrophages also show strong correlation with pericytes in GBM ([ρ] = 0.43; **Supplementary Fig. 16a**), potentially driven by hypoxia-induced necrosis. However, correlations were weaker overall in GBM than in other cancer types. In HNSCC, the proportion of immune cell types were highly correlated with each other (**Supplementary Fig. 16b**). We also noted a high correlation between most immune cell types and endothelial cells. In melanoma, endothelial cells had a relatively strong positive association with fibroblasts (ρ = 0.62; **Supplementary Fig. 16c**), which may be consistent with reports that fibroblast ECM remodelling promotes angiogenesis in melanoma[8]. We also noted two separate submodules of highly correlated immune cells, one consisting of CD4+ T-cells and B cells, and the other consisting of CD8+ T-cells and macrophages (**Supplementary Fig. 16c**). This finding suggests that melanoma patients may have two distinct types of immune response, mediated by humoral and cellular immunity, to varying degrees between patients.

#### Supplementary Note 7: Validating correlations.

Discovering genes involved in tumor-microenvironment interactions is a major unresolved problem. Discovering candidate interactions in which gene expression in malignant cells is correlated with fractions of non-malignant cells is a potentially powerful strategy that has identified strong candidates which have been validated using single-cell RNA-seq studies. However, single cell RNA-seq studies are underpowered to discover these associations due to small sample size, providing a motivation to use the large quantities of existing bulk-RNA-seq. The main challenge in this problem is that correlations between gene expression and fractions of query cell types in bulk data are prone to false positives when a gene is expressed highly and specifically in the query cell type. We reasoned that BayesPrism recovered malignant cell expression accurately enough to avoid this type of false positive. To test this hypothesis, we examined the distribution of marker genes that were specifically expressed in each cell type based on independently derived data. Whereas marker genes had systematically higher correlations in the bulk data, these were reduced to a median near 0 when using the estimates of expression in malignant cells produced by BayesPrism (**Supplementary Fig. 19**). This suggests that BayesPrism expression estimates are effective in removing many trivial false positives.

**Supplementary Note 8: Concordance between the tumor-microenvironment correlation identified using gene programs and using individual genes.**

Two approaches have been used to summarize the correlation between the gene expression in malignant cells and cell type fractions of non-malignant cells. In the first approach, we computed the genewise correlation between its inferred expression in malignant cells and the fraction of a non-malignant cell type, and then performed gene set enrichment analysis over the rank order of the correlation coefficient for each non-malignant cell type (**Fig. 3d** and **Supplementary Fig. 21**). In the second approach, we used the embedding learning module of BayesPrism to factorize the expression of malignant cells into the linear combination of multiple malignant bases. Each tumor sample was assigned with a group of weights over these bases. We then perform differential gene expression over a set of representative core samples for each tumor basis (**Supplementary Table 5**, **Supplementary Fig. 24**), followed by gene set enrichment analysis over the Wald statistics of the differential expression (**Fig. 4e-g**, **Supplementary Table 4**). Two approaches tackle the problem from different angles. The genewise approach studies the correlation with a particular non-malignant cell type of interest, while the gene program level approach focuses on the selection of a group of co-expressed genes, regardless of the correlation with non-malignant cell types. Although the embedding learning approach does not directly look for correlations with the fraction of non-malignant cells, we found that it yielded many observations that were concordant with the results obtained from the genewise approach, which are summarized below.

| | Direction of correlation with cell type fractions | Consistently enriched Biological processes |
|---|---|---|
| GBM | | |
| program-1 | endothelial(-) | respiration / oxidative phosphorylation pathways |
| program-2 | oligodendrocyte(+) | neuronal processes |
| program-3 | oligodendrocyte(-) | DNA conformational change and RNA processing |
| program-4 | macrophage(+); endothelial(+); pericyte(+) | hypoxia, immune response, EMT and remodeling of extracellular matrix |
| program-5 | macrophage(-) | Proliferation, DNA replication and cell cycle |
| HNSCC | | |
| program-1 | B cell(-); T cell(-) | EMT |
| program-2 | endothelial(+); T cell(+) | Hallmark E2F targets (DNA replication) |

| | | |
|---|---|---|
| program-3 | fibroblast(-); macrophage(+); mast cell(-) | Respiration [fibroblast(-)]; Interferon [fibroblast(-) & macrophage(+)]; DNA repair [mast cell(-)] |
| program-4 | endothelial(-); fibroblast(-); T cell(-); dendritic cell(-); macrophage(-); mast cell(+) | keratinization |
| SKCM | | |
| program-2 | fibroblast(+); endothelial(+) | EMT, NFkB, hypoxia, cell adhesion, extracellular matrix organization, vesicle transport/budding, and ER stress |
| program-3 | CD4+ T cell(+); CD8+ T cell(+); NK cell(+); macrophage(+); B cell(+) | Immune response |
| program-5 | macrophage(-); fibroblast(-) | RNA processing [macrophage(-)]; Respiration [fibroblast(-)] |
| program-6 | endothelial(+) | EMT |

#### Supplementary Note 9: Spatial heterogeneity in cell type composition in GBM

To understand the spatial distribution of cells within a tumor, we applied BayesPrism to 122 bulk RNA-seq samples from the Ivy Glioblastoma Atlas Project (IVY GAP) that interrogate laser microdissected tissue from GBM[9]. Data was available for ten tumors microdissected into five structures: leading edge (LE), infiltrating tumor (IT), cellular tumor (CT), microvascular proliferation (MVP) and pseudopalisading cells around necrosis (PAN) (**Fig. 5a and Supplementary Table 3b**).

As we expected normal brain cells, including neurons, at high abundance in the leading edge and infiltrating tumor structures[9], we deconvolved all samples using a reference scRNA-seq dataset by combining refGBM8 with adult human neurons collected using Fluidigm[10]. As batch effects exist between refGBM8 and Fluidigm, the absolute fraction of cell types can be affected. In particular, as the neuron cell reference collected by Fluidigm was sequenced to a much higher total sequencing depth and detected more genes than refGBM8, it may absorb more reads than other cell types in refGBM8, and hence its absolute fraction is likely to be overestimated. This is similar to what we observe for T cells in the pseudo-bulk deconvolution of GBM28 using refGBM8 (see **Supplementary Fig.6o**). We observed that due to the significantly lower number of total reads caused by the scarcity of T cells in GBM, the inferred absolute fraction was off by a linear factor, while the relative fraction across samples remained accurate. By the same token, the relative cell type fractions of the deconvolved IVY-GAP data shall also remain accurate, and hence the results from statistical tests remain valid.

BayesPrism revealed several striking features of GBM regional cellular heterogeneity (**Fig. 5b**). First, pericytes and endothelial cells were significantly enriched in regions of microvascular proliferation ($p$ < 1e-4, linear mixed model, see Methods), and comprised nearly 60% of cells in these regions. Second, oligodendrocytes and adult neurons were enriched in the leading edge and infiltrating tumor ($p$ < 1e-4, linear mixed model), with a relative magnitude that matches H&E stained sections from these same patients[9]. Third, pseudopalisading cells around necrosis showed a depletion of endothelial cells ($p$ = 0.0499, linear mixed model) and enrichment for T cells ($p$ = 0.0048, linear mixed model) and macrophages ($p$ = 0.0140, linear mixed model). Fourth, leading edges were enriched for macrophages ($p$ = 0.0138, linear mixed model) and depleted of pericytes ($p$ = 0.0423, linear mixed model). Taken together, BayesPrism revealed a rich and highly heterogeneous picture of cell type composition.
